# Phylogenomics resolves a 200-year-old puzzle: a revised tribal classification of Afro-Eurasian dung beetles (Coleoptera: Scarabaeinae)

**DOI:** 10.64898/2026.07.22.740134

**Authors:** Giulio Montanaro, Fernando Lopes, Nicole L. Gunter, Clarke Scholtz, Adrian L. Davis, Federica Losacco, Michele Rossini, Conrad P.D.T. Gillett, Natalie A. Saxton, Rachel L. Stone, Gimo M. Daniel, Sergei Tarasov

**Affiliations:** Finnish Museum of Natural History, University of Helsinki, Pohjoinen Rautatiekatu 13, 00100 Helsinki, Finland; State University of Campinas, Institute of Biology, R. Monteiro Lobato, 255 - Cidade Universitária, Campinas, Brazil. 13083-862; Biodiversity and Geosciences Program, Queensland Museum Kurilpa, South Brisbane, Queensland, Australia; Department of Biology, Case Western University, Cleveland, Ohio, USA; Department of Zoology and Entomology, University of Pretoria, P/B X28, Hatfield, South Africa; Instituto Nacional de Coleoptera (INCol), Conselho Nacional de Desenvolvimento Científico e Tecnológico (CNPq), Brazil; Laboratório de Scarabaeoideologia, Departamento de Biologia e Zoologia, Universidade Federal de Mato Grosso, Cuiabá, Brazil; Department of Terrestrial Invertebrates, National Museum, 36 Aliwal Street, Bloemfontein, South Africa; Department of Biological & Environmental Sciences, Walter Sisulu University, Mthatha, South Africa

## Abstract

**Background:** The tribal classification of scarab dung beetles (Coleoptera: Scarabaeinae) is currently largely incomplete due to the lack of robust phylogenetic evidence supporting the assignment of many Afro-Eurasian and American genera to tribes.

**Methods:** We used ultraconserved elements (UCEs) to infer phylogenetic relationships across most Afro-Eurasian dung beetle lineages, including 25 of the 27 extant genera currently *incertae sedis*.

**Results:** We recovered full support for the monophyly of previously recognised tribes and for several phylogenetically and morphologically clearly delimited new tribal-level clades, allowing us to propose a complete tribal classification for all Afro-Eurasian dung beetle genera. Thirteen new tribes are described and diagnosed: Aphengoecini ***trib. nov.***, Bohepilissini ***trib. nov.***, Catharsiini ***trib. nov.***, Chalconotini ***trib. nov.***, Circelliini ***trib. nov.***, Dwesasilvasedini ***trib. nov.***, Haroldiini ***trib. nov.***, Heliocoprini ***trib. nov.***, Janssensantini ***trib. nov.***, Macroderini ***trib. nov.***, Nesovinsoniini ***trib. nov.***, Pycnopanelini ***trib. nov.*** and Tanzanolini ***trib. nov***. The tribe Coprini ***sensu novo*** is redefined as comprising three subtribes: Coprina, Onychothecina ***subtrib. nov.*** and Pedariina ***subtrib. nov***. The tribe Odontolomini is downranked to a subtribe of Endroedyolini ***sensu novo***, becoming Endroedyolini Odontolomina ***stat. nov***. The tribe Onthophagini ***sensu novo*** is redefined and divided into three subtribes: Helictopleurina ***stat. nov.***, Oniticellina ***stat. nov.*** and Onthophagina; the remaining former subtribes of Oniticellini (Attavicinina, Drepanocerina and Liatongina) are synonymised with Oniticellina; the subtribe Alloscelina of Onthophagini is synonymised with Onthophagina (***syn. nov.***). The tribe Panelini ***stat. rev.*** and ***sensu novo***, comprising the single genus *Panelus*, is revalidated and redefined. Updated morphological diagnoses of Elassocanthonini, Gymnopleurini, Onitini and Scarabaeini are provided. The genus *Phaedotrogus* is synonymised with *Haroldius* (Haroldiini) (***syn. nov.***). An identification key to all Afro-Eurasian dung beetle tribes is provided.

**Discussion:** Our results establish a robust phylogenetic framework and revised tribal classification for Afro-Eurasian Scarabaeinae dung beetles, providing a foundation for future taxonomic, comparative and macroevolutionary research.

## INTRODUCTION

Dung beetles (Coleoptera: Scarabaeidae: Scarabaeinae) form a spectacular radiation, comprising ∼300 genera and nearly 7,000 described species worldwide (Schoolmeesters, 2026). Their ecological importance and their role as bioindicators is widely recognised, and they are well-established models in ecology and evolutionary biology because of their exceptional diversity, habitat specialisations, horn evolution, and reproductive strategies (Hanski and Cambefort, 2014; Scholtz et al., 2009; Simmons and Ridsdill-Smith, 2011). This extraordinary richness is paralleled by a long and turbulent systematic history resulting from extensive morphological convergence, with many longstanding uncertainties only beginning to be resolved through phylogenetic approaches in recent years (Daniel and Davis, 2024; Gunter et al., 2026; Tarasov and Dimitrov, 2016).

For much of its history, Scarabaeinae classification was dominated by the deceptively simple dichotomy of “rollers”, typically long-legged species with a ball-rolling behaviour, *versus* “tunnelers”, short-legged species burying their provisions into subterranean tunnels dug under the dung. These two morphogroups were further subdivided into several tribes and subtribes based on morphology (Hanski and Cambefort, 2014; Janssens, 1949; Philips, 2011; Scholtz, 2009). However, while some such taxa were phenotypically homogeneous groups of closely related genera (*e.g.*, Scarabaeini, Gymnopleurini, and Onitini), others were artificial assemblages defined by overall body shape and a few homoplastic characters. The most striking examples are the historical concepts of Canthonini, comprising more than 100 genera sharing a general roller morphology, and the tunneler tribes Ateuchini, Coprini and Dichotomiini, among which numerous genera shifted back and forth throughout their taxonomic history (Montreuil, 1998; Tarasov and Dimitrov, 2016; Vaz-De-Mello, 2008; Lopes et al., 2024).

The advent of molecular phylogenetics and phylogenomics demonstrated that the traditional roller/tunneler dichotomy was largely artificial, while several historically recognised “dumping ground” tribes were largely polyphyletic, with similar morphology being a result of convergence (Gunter et al., 2016, 2026; Lopes et al., 2024; Mlambo et al., 2015; Sole and Scholtz, 2010; Tarasov, 2017; Tarasov and Dimitrov, 2016). The most taxonomically comprehensive analysis to date restricted the definition of some tribes to include only phylogenetically and morphologically closely related taxa, resulting in more than 100 genera becoming *incertae sedis* (Tarasov and Dimitrov, 2016). The Canthonini were redefined to include only 22 genera closely related to *Canthon* Hoffmansegg, 1817 and *Deltochilum* Eschscholtz, 1822, and are now recognised under the senior synonym Deltochilini (Bouchard et al., 2011) (in the text, we use Canthonini or canthonines to refer to their historical concept). Ateuchini, Coprini and Dichotomiini underwent a similar restructuring. Subsequently, several recent works contributed to the definition of additional tribes, bringing their number to 24 and reducing that of *incertae sedis* genera to 45 (Davis et al., 2019, 2025; Génier and Darling, 2024; Gunter et al., 2026; Lopes et al., 2024; Rossini et al., 2022; Scholtz et al., 2025; Tarasov, 2017; see supplementary file S5 for a full list).

Afro-Eurasia—uniting the Afrotropical, Indomalayan and Palearctic biogeographical realms (Olson et al., 2001)—has the greatest richness of scarabaeine genera (167) and tribes (14), and accounts for the majority of extant *incertae sedis* genera (30). Over the past eight years, several well-supported clades of historical Afro-Eurasian canthonines and ateuchines were transferred to newly established or redefined pre-existing tribes following the work by Tarasov and Dimitrov (2016): Parachoriini (Tarasov, 2017); Elassocanthonini, Endroedyolini and Odontolomini (Davis et al., 2019, 2025); Epactoidini (Rossini et al., 2022); Coprini (Lopes et al., 2024); and Coptorhinini and Paraphytini (Scholtz et al., 2025). On the other hand, isolated positions in the phylogeny and poorly supported affinities with other lineages precluded any formal new tribal accommodation for the remaining taxa (Davis et al., 2019; Scholtz et al., 2025).

Afro-Eurasian *incertae sedis* genera are concentrated in the Afrotropical realm, of which 23 are endemic. Many of them are species-poor genera of former canthonines with narrow, relictual distributions centered in southern and eastern Africa (Davis et al., 2008, 2020). To this group belong forest specialists found in South Africa (*Bohepilissus* Paulian, 1975 and *Dwesasilvasedis* Deschodt & Scholtz, 2008), southern and eastern Africa (*Gyronotus* Van Lansberge, 1874), and the Eastern Arc Mountains and South East Africa Montane Archipelago (*Janssensantus* Paulian, 1976 and *Tanzanolus* Scholtz & Howden, 1987—although the former also comprises a species from miombo in Zambia (Josso, 2022; Montanaro *et al*., in preparation)). Other relictual canthonines are restricted to coastal sandveld of South Africa (*Aphengoecus* Péringuey, 1901 and *Circellium* Latreille, 1829), in coastal shrublands of Mozambique (*Canthodimorpha* Davis, Scholtz & Harrison, 1999) and in arid regions of southwestern Africa (*Hammondantus* Cambefort, 1978). *Panelus* Lewis, 1895 and *Pycnopanelus* Arrow, 1931 show disjunct Afro-Indomalayan distributions, with the former comprising one Afrotropical species and many Indomalayan ones, and the latter having three species scattered through southwestern and northeastern Africa and India. Only one genus, *Chalconotus* Dejean, 1833, has a truly pan-African distribution, comprising the most common and widespread canthonines of the continent. Outside of Africa, the monotypic *Nesovinsonia* Martinez & Pereira, 1959 is restricted to Mauritius in the Mascarene island archipelago (Lopes et al., 2023).

On the other hand, the Afro-Indomalayan myrmecophilous genus *Haroldius* Boucomont, 1914 and its poorly known close relative *Phaedotrogus* Paulian, 1985 had a rather troubled classificatory history, being placed in Ateuchini, Canthonini or Onthophagini (including its synonym Alloscelina) by different authors (Janssens, 1949; Krikken and Huijbregts, 2006; Montreuil, 2010; Philips, 2016; Krell, 2010).

The remaining *incertae sedis* taxa are tunnellers. *Pedaria* Castelnau, 1832, restricted to the Afrotropical realm, was historically considered a member of Ateuchini *sensu lato*, as it did not fit into either the subtribe Ateuchina or the Scatimina (Vaz-de Mello, 2007). *Macroderes* Westwood, 1842 and *Xinidium* Harold, 1869, two southern African endemics, were either placed in Coprini or Dichotomiini (Cambefort, 1985; Montreuil, 1998). The charismatic *Catharsius* Hope, 1837, currently one of the most species-rich genera of horned tunnellers in the world, and its relatives *Catharsiocopris* Balthasar, 1967, *Copridaspidus* Boucomont, 1920 and *Metacatharsius* Montreuil, 1998, were historically placed in Coprini (Balthasar, 1967; Montreuil, 1998). The same applies to *Heliocopris* Hope, 1837 and *Synapsis* Bates, 1868, which were long considered close coprine relatives (Montreuil, 1998), although some authors suggested their placement in the Dichotomiini (Edmonds and Halffter, 1978). Lastly, *Lobateuchus* Montreuil, Génier & Nel, 2010, from the Lower Eocene (Oise amber), is the only fossil genus reliably attributable to Scarabaeinae that currently lacks a tribal placement (Montreuil et al., 2010). Montreuil et al. (2010) and Tarasov et al. (2016) suggested that it may belong to the Ateuchini, which would make it the only known Afro-Eurasian representative of an otherwise entirely American tribe. However, the difficulty of scoring morphological characters from the sole known specimen renders this assignment uncertain.

Overall, morphological data have often proven insufficient to resolve deeper phylogenetic relationships and establish a stable higher-level classification in dung beetles. In contrast, recent advances in systematics across diverse organisms have been driven by phylogenomic approaches based on genome-scale data (Delsuc et al., 2005; Dunn et al., 2008; Lee and Palci, 2015; Steenwyk et al., 2023). Within this context, ultraconserved elements (UCEs) have emerged as a powerful marker system for phylogenetic inference, combining highly conserved core regions with more variable flanking sequences that provide informative signal across both shallow and deep evolutionary timescales (Baca et al., 2017; Faircloth et al., 2012; Gustafson et al., 2020; Zhang et al., 2019; Hellemans et al., 2022). The development of a scarabaeoid-specific probe set (Gustafson et al., 2023) has already enabled substantial progress in dung beetle systematics, resolving several long-standing classification challenges (Génier et al., 2025; Gunter et al., 2026; Lopes et al., 2024). UCE-based phylogenomic analyses provide a robust framework for establishing a global dung beetle classification and are therefore employed in this study.

Our phylogeny includes all Afro-Eurasian dung beetle lineages (*i.e.*, isolated genera or monophyletic groups of genera highly supported by previous evidence) except for *Aphengoecus* (*incertae sedis*) and Parachoriini. Our analyses provide strong support for the delimitation of 13 new tribes and the redefinition of nine existing ones, leading to a complete resolution of the tribal placement of all extant Afro-Eurasian genera. However, although phylogenetically resolved, we do not treat in detail two endemic Madagascan clades (the *Epilissus* and *Nanos* generic groups), whose discussion is left for a forthcoming study (Rossini *et al*., in preparation). All treated tribes are diagnosed based on unique combinations of morphological characters, and an identification key is provided to facilitate their identification. This study represents a turning point in dung beetle systematics, proposing a classification that robustly resolves more than 200 years of uncertainty and leaving only 22 genera, mostly from the Americas, to be placed at the tribal level.

This manuscript conforms to the requirements of the amended International Code of Zoological Nomenclature (ICZN). Please note that this work is not issued for the purpose of zoological nomenclature and it should be considered not published within the meaning of the code (ICZN code, Section 8.2). The nomenclatural acts and new taxon names described herein should therefore be considered as not available.

## MATERIALS AND METHODS

### Molecular dataset construction

We obtained UCE sequence data from 316 taxa (see supplementary file S1), comprising an ingroup of 288 Scarabaeinae and an outgroup of 28 taxa in the wider superfamily Scarabaeoidea and the family Silphidae. These data represent ∼48% of Afro-Eurasian and ∼29% of global dung beetle generic diversity. Representatives of all described Afro-Eurasian tribes, with the exception of Parachoriini Tarasov, 2017, were included, in addition to the American Phanaeini and the Australasian Mentophilini.

The dataset comprises 25 out of 29 valid extant Afro-Eurasian *incertae sedis* genera, omitting only *Aphengoecus*, *Canthodimorpha* (a well-supported sister taxon to *Chalconotus*), *Catharsiocopris* and *Phaedotrogus*. The latter genus is herein synonymised with *Haroldius*, whereas *Catharsiocopris* is an undoubted but as yet unpublished synonym of *Catharsius*, based on morphological and molecular evidence (Takano, 2018). Hence, we ultimately cover 25 out of 27 extant *incertae sedis* genera. Sequences were newly generated (191 taxa) or were retrieved from the published literature (125 taxa; Gunter et al., 2026; Gustafson et al., 2023; Lopes et al., 2024; Losacco et al., ress) (see supplementary file S1). Notably, we obtained the first molecular data for the rare Mauritian endemic *Nesovinsonia*, the Madagascan *Apterepilissus*, and the Indomalayan *Synapsis*. Specimens from Madagascar and Mauritius were collected under authorisation from the Malagasy Institut pour la Conservation des Écosystèmes Tropicaux (MICET) (permit #443/21/MED/SG/DGGE/DAPRNE/SCBE.Re) and the Mauritian National Parks and Conservation Service (NPCS) (permit NP46/3V5), respectively.

DNA extraction was performed using the QIAamp DNA Micro Kit (QIAGEN) following the manufacturer’s instructions, except for an old dry-preserved specimen of *Apterepilissus ovalis* (Felsche, 1911) for which we utilised the archival DNA protocol by Lopes et al. (2024). UCEs were captured and sequenced using the Scarabaeinae probe set Scarab 3Kv1 (Gustafson et al., 2023). For data processing, data matrix construction and phylogenomic analyses, we followed the workflow and parameter settings of Lopes et al. (2024).

For phylogenomic analyses, we constructed data matrices in Phyluce 1.7.3 (Faircloth, 2016) with completeness of 50% (hereafter dataset 50p) and 70% (hereafter dataset 70p), allowing up to 50% and 30% missing taxa per locus, respectively. These completeness thresholds have previously generated reliable phylogenomic inferences (*e.g.*, Lopes et al., 2024; Gunter et al., 2026; Losacco et al., ress) and were therefore also applied in the present study.

### Phylogenetic inference

Phylogenetic analyses were performed in a maximum likelihood framework in IQ-TREE 2.0.7 (Minh et al., 2020). We inferred species trees from analysis of concatenated datasets using UCEs as independent loci (trees 50p-1 and 70p-1) and after partitioning the datasets with the entropy-based method PFinderUCE-Sliding-Window Site Characteristics (SWSC-EN) (Tagliacollo and Lanfear, 2018) (trees 50p-2 and 70p-2). Node support was estimated using 1,000 ultrafast bootstrap (UFBoot) replicates and the Shimodaira–Hasegawa approximate likelihood ratio test (SH-aLRT) (Goldman et al., 2000; Shimodaira and Hasegawa, 1999). Additionally, we inferred species trees from gene trees derived from both the UCE datasets (trees 50p-3 and 70p-3) and the SWSC-EN partitioned datasets (trees 50p-4 and 70p-4) using ASTRAL v5.7.1. ASTRAL implements the multispecies coalescent (MSC) model and estimates species trees by summarizing a set of independently inferred gene trees while accommodating discordance among loci. This approach accommodates discordance among the evolutionary histories of different genomic regions, frequently resulting from incomplete lineage sorting (Mirarab et al., 2014; Mirarab and Warnow, 2015; Zhang et al., 2018). Lastly, to account for heterogeneous evolutionary rates across sites (*i.e.*, heterotachy), we used the GHOST mixture model implemented in IQ-TREE (50p-5 and 70p-5) (Crotty et al., 2020). All the analyses are summarised in Table 1.

**Table 1.** Phylogenomic inferences performed in this study. Datasets with 50% and 70% completeness include only loci that are present in at least 50% or 70% of the sampled taxa, respectively. Dataset partitioning is either done considering all UCEs as single loci, or by grouping them in partitions inferred with the entropy-based method PFinderUCE-Sliding-Window Site Characteristics (SWSC-EN). Phylogenetic inference was performed in IQ-TREE using standard maximum likelihood (ML) or by employing the GHOST mixture model, or by using the multispecies coalescent model as implemented in ASTRAL.

| Tree | Dataset completeness | Dataset partitioning | Inference method |
| --- | --- | --- | --- |
| 50p-1 | 50% | UCEs as single loci | standard ML |
| 50p-2 | 50% | SWSC-EN partitions | standard ML |
| 50p-3 | 50% | UCEs as single loci | ASTRAL |
| 50p-4 | 50% | SWSC-EN partitions | ASTRAL |
| 50p-5 | 50% | UCEs as single loci | GHOST |
| 70p-1 | 70% | UCEs as single loci | standard ML |
| 70p-2 | 70% | SWSC-EN partitions | standard ML |
| 70p-3 | 70% | UCEs as single loci | ASTRAL |
| 70p-4 | 70% | SWSC-EN partitions | ASTRAL |
| 70p-5 | 70% | UCEs as single loci | GHOST |

We quantified topological differences among inferred trees using Robinson–Foulds (RF) distances (Robinson and Foulds, 1981) and visualised the pairwise distance matrix with multidimensional scaling (MDS) to assess relationships among the obtained topologies.

### Characterisation of new tribes

#### Criteria for the establishment of new taxa

We used molecular phylogenetic results, clade stability (100% support across at least all 70p analyses), and morphological congruence as criteria for establishing tribal-level groups. Earlier phylogenetic studies were taken into account to infer the placement of two genera not included in our dataset (*Aphengoecus* and *Canthodimorpha*). The fossil *Lobateuchus* was excluded from tribal delimitation owing to insufficient evidence for its phylogenetic placement.

We sought to optimise the assignment of either tribal or subtribal rank by evaluating the following criteria for each identified clade and its sister clade: 1) the depth of divergence; 2) the degree of morphological distinction; 3) the existence of substantial biogeographical and/or ecological differences between the clades; 4) classification consistency—*i.e.*, we sought to avoid unnecessary redefinition of previously well-established tribal concepts. Informal names were assigned to two well-supported clades consistently recovered in both our analyses and previous studies, thereby facilitating reference to these putative evolutionary lineages in future studies.

#### Morphological examination

Morphological diagnoses were based on examinations of the external and genital morphology of the included genera. For each (sub)tribe, we selected a unique combination of characters that unambiguously distinguishes the taxon from all others recognised herein. Firstly, we scored synapomorphies identified by Tarasov and Génier (2015) and used them to construct backbone diagnoses. These were supplemented with additional characters where necessary to ensure unambiguity. To verify that each diagnosis delimited only the intended group of genera, we used a custom R script to identify any other genera in Tarasov and Génier (2015)’s matrix that shared the same combination of character states. A few genera for which detailed morphological information was not available from previous studies (*Nesovinsonia*, *Panelus*, *Synapsis*) were examined *ex novo*.

Studied specimens were either fully disarticulated and cleaned following the protocol of Tarasov and Génier (2015), or only partially dissected when this was sufficient to observe the relevant characters. In heavily sclerotised species, the elytra and major body sclerites were bleached in hydrogen peroxide to facilitate visualisation of morphological structures. The examined material is housed primarily at the Finnish Museum of Natural History (Helsinki, Finland), which contains most of the fully disarticulated specimens, as well as in the additional institutional and private collections listed in the Acknowledgments.

#### Anatomical terminology

Anatomical terminology follows Tarasov and Génier (2015), Tarasov and Solodovnikov (2011), Lawrence and Slipinski (2013), Génier (2019) and Montanaro et al. (2024a). We use the term “uncus” (Lawrence and Slipinski, 2013) to refer to the cuticular spine on the distoventral margin of (usually male) protibia (for example, see Fig. 17C). This structure has received many names: “éperon” (Janssens, 1937), “finger-like process” (Krikken, 1977), “apical digit” (Gunter and Weir, 2019), “tooth” (Génier and Moretto, 2019), “internoapical tooth” (Génier and Moretto, 2017), “spur” (Scholtz and Howden, 1987), etc. This plethora of terms is also inconsistently applied to other protibial structures, such as the flattened protrusions on the dorsal margin or the articulated distal spine (which should more appropriately be referred to as teeth and spur, respectively). We therefore propose “uncus” as a standardised term in scarabaeine taxonomy to refer specifically and unambiguously to the structure defined above. Abbreviations for terms used in the text are provided in Table 2.

**Table 2.** Abbreviations of morphological terms used in the text.

| Abbreviation | Description |
| --- | --- |
| AS | Additional endophallite |
| BSc | Basal semicircular endophallite |
| Cav | Antennal cavity |
| Dep | Depression |
| EpC | Epipleural carina |
| EP | Epipleuron |
| TS | Spur-shaped lobe of A+?SA complex |
| MsEp | Mesanepesternum |
| MtEp | Metanepesternum |
| MtSc | Metathoracic scutellum |
| C | Elytral carina |
| F | Flagellomere |
| FLP | Fronto-lateral peripheral endophallite |
| LC | Lamella copulatrix |
| A | Axial endophallite |
| SA | Subaxial endophallite |
| SRP | Superior right peripheral endophallite |
| rsp | Raspula |
| PHC | Posterior hypomeral carina |
| AHC | Anterior hypomeral carina |
| LPC | Lateral pronotal carina |
| TFP | Trochanterofemoral pit |
| Tub | Tubercle |
| X | Sclerite of unclear homology |

## RESULTS

### Phylogenetic relationships

Trees obtained with different analyses (Figs. 1–2 and supplementary files S2 and S3) recovered partly different relationships. Those inferred with ASTRAL (50p-3, 50p-4, 70p-3 and 70p-4) showed lower support for several nodes and were particularly divergent from the IQ-tree trees according to Robinson– Foulds distances (see supplementary file S4). The 70p-1, 70p-2, 70p-5 and 50p-1 trees were more similar to each other and generally better supported, whereas the 50p-2 and 50p-5 reconstructions clustered together based on RF distances despite exhibiting some notable topological differences.

**Figure 1.**
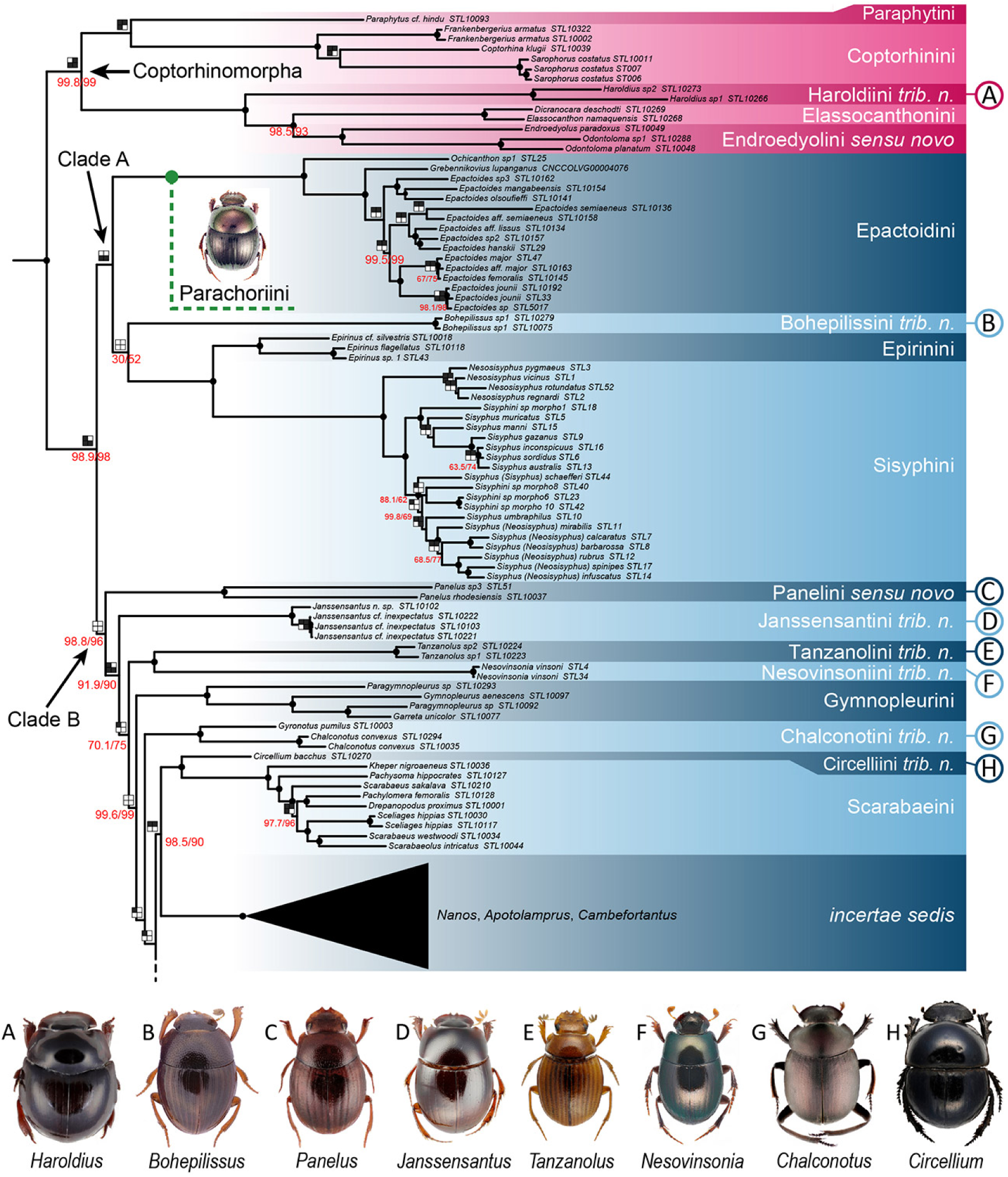
Phylogenetic tree 70p-1 (*partim*) showing new Scarabaeinae tribal classification. Tree 70p-1 was selected for visualization as it is better supported and more consistent with previous evidence. Node support in tree 70p-1 is compared to that in 70p-2, 70p-3, 70p-4 and 70p-5. For each node, black dots represent 100% support in all analyses; values in red are SH-aLRT (left) and bootstrap (right) support lower than 100% in tree 70p-1; 2×2 grids represent presence (black filling) or absence (white filling) of the node in 70p-2 (top left), 70p-3 (top right), 70p-4 (bottom left) and 70p-5 (bottom right) trees. See also the legend in Figure 2. The putative position of Parachoriini is based on the results by Tarasov (2017).

**Figure 2.**
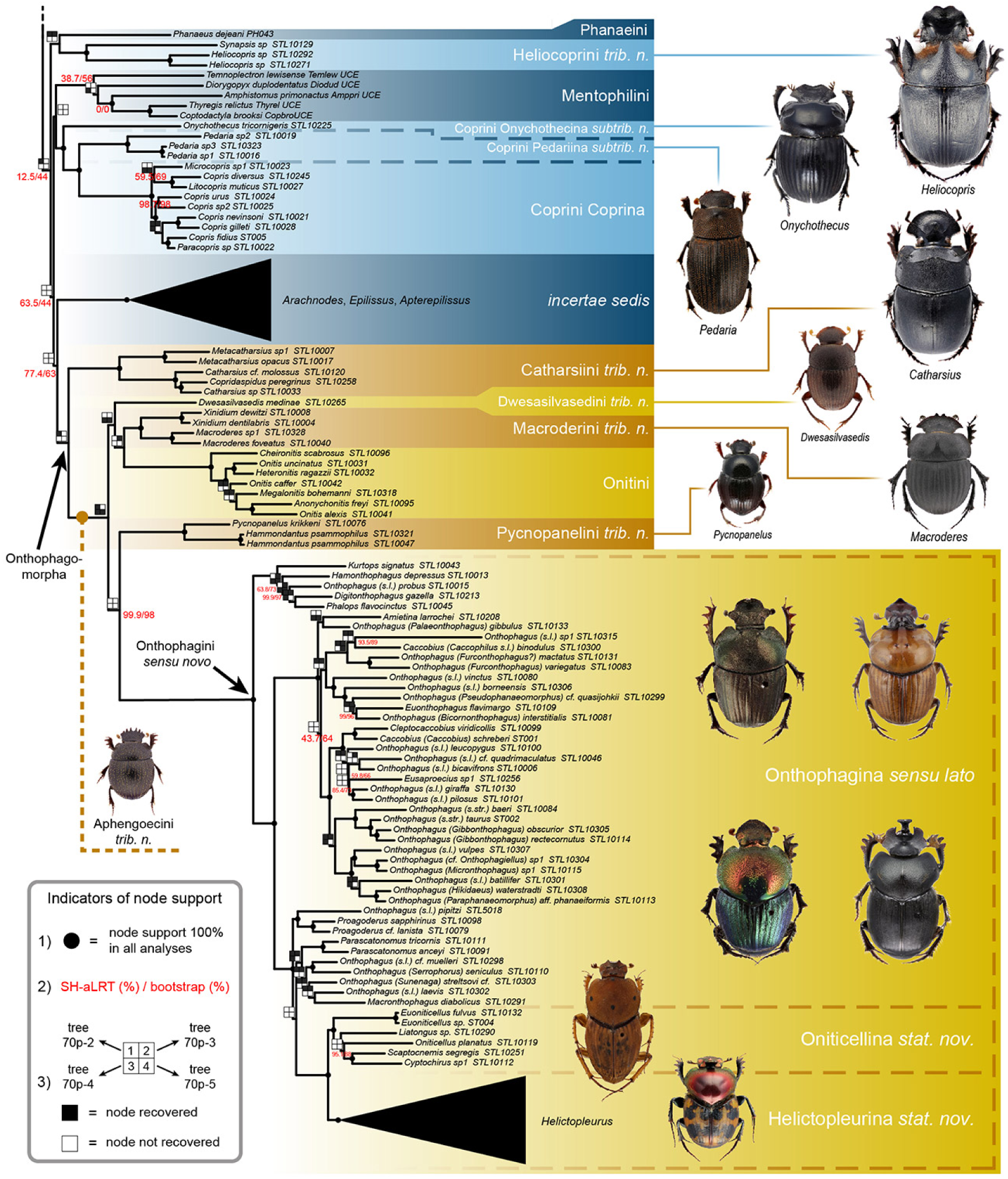
Phylogenetic tree 70p-1 (continuation) showing new Scarabaeinae tribal classification. Node support in tree 70p-1 is compared to that in 70p-2, 70p-3, 70p-4 and 70p-5. For each node, black dots represent 100% support in all analyses; values in red are SH-aLRT (left) and bootstrap (right) support lower than 100% in tree 70p-1; 2×2 grids represent presence (black filling) or absence (white filling) of the node in 70p-2 (top left), 70p-3 (top right), 70p-4 (bottom left) and 70p-5 (bottom right) trees. The putative position of Aphengoecini is based on the results by Tarasov and Dimitrov (2016).

The trees were largely consistent at intermediate levels of divergence, recovering congruent tribal-level groupings of genera (see supplementary files S2–S3). The well-established tribes Coptorhinini, Elassocanthonini, Endroedyolini, Epactoidini, Epirinini, Gymnopleurini, Mentophilini, Odontolomini, Onitini, Scarabaeini and Sisyphini were always monophyletic with 100% support. Oniticellini was found nested within a paraphyletic Onthophagini without exception, with the Madagascan genus *Helictopleurus* being sister to the remaining oniticellines. Full support was also recovered for *Chalconotus*+*Gyronotus*, *Hammondantus*+*Pycnopanelus*, *Macroderes*+*Xinidium*, and *Catharsius*+*Metacatharsius*, with *Copridaspidus* nested within *Catharsius*. The Madagascan endemic *incertae sedis* genera always formed two distinct, distantly related clusters, one including *Apotolamprus*+*Cambefortantus*+*Nanos* and one including *Apterepilissus*+*Arachnodes*+*Epilissus*. All analyses recovered 100% support for *Pedaria* nested within Coprini, with *Onychothecus* being the earliest-branching lineage of the clade. *Haroldius* was recovered with 100% support as sister to Elassocanthonini+(Endroedyolini+Odontolomini). The South African relict genus *Circellium* was consistently recovered as sister to Scarabaeini in all five 70p reconstructions, whereas the 50p trees placed it in varying positions, either as sister to Scarabaeini, Coprini or the *Nanos* group of genera. Similarly, all 70p trees found a sister relationship between *Synapsis* and *Heliocopris*, whereas only the 50p-3 and 50p-4 trees recovered the same topology. In contrast, the 50p-1 and 50p-5 trees placed *Synapsis* as sister to Onthophagini, while the 50p-2 tree recovered it as sister to a large heterogenous clade. The former canthonine genera *Bohepilissus*, *Janssensantus* and *Panelus* were consistently recovered in rather isolated positions, exhibiting deep divergence from their closest inferred relatives. In contrast, *Nesovinsonia* and *Tanzanolus* consistently formed a fully supported clade, despite the very deep divergence between the two.

At deeper levels, phylogenetic discordance among inferences was more pronounced. Several genera, especially *Janssensantus*, *Panelus* and *Tanzanolus*+*Nesovinsonia*, shifted across different positions depending on the analysis. Nonetheless, some higher-level clusters were consistently recovered across most of the trees. A clade comprising of *Dwesasilvasedis*, *Macroderes*+*Xinidium*, Onitini, Onthophagini+Oniticellini and *Pycnopanelus*+*Hammondantus*, was recovered in all 70p analyses except the GHOST inference (70p-5), as well as in 50p-1 and 50p-2 inferences, which additionally included *Synapsis*. The remaining 50p inferences (ASTRAL and GHOST) separated Onthophagini+Oniticellini from the other taxa. However, reciprocal relationships of lineages within the clade varied across reconstructions. Moreover, this large clade had a sister relationship with *Catharsius*+*Metacatharsius*+*Copridaspidus* in all except the ASTRAL analyses. Epactoidini, Epirinini and Sisyphini together formed a clade in all analyses, with *Bohepilissus* being either sister (50p-1, 70p-3 and 70p-4) or a nested member (70p-1) or only distantly related (in the remaining trees). The clade (Coptorhinini+Paraphytini)+(*Haroldius*+(Elassocanthonini+ (Endroedyolini+Odontolomini))) was recovered with an identical topology and with mostly 100% support across most analyses, usually as sister to the remaining scarabaeines. In a few cases, Paraphytini were inferred as basal, either to the remaining taxa (70p-4) or to all Scarabaeinae (50p-2), while in the 70p-2 analysis Coptorhinini+Paraphytini were inferred as basal to the rest of the subfamily. A sister relationship between Coprini (including *Pedaria*) and the *Epilissus* group of genera was recovered by several analyses (all 70p trees except 70p-1, as well as 50p-1, 50p-3 and 50p-4). *Chalconotus*+*Gyronotus* were also frequently inferred as sister to Scarabaeini, either directly or with *Circellium* intervening between the two clades. Overall, the 50p-5 and 70p-5 GHOST analyses resulted in the most pronounced outlier patterns in inter-tribal relationships, for example recovering Onthophagini (plus *Synapsis* in 50p-5) as sister to the remaining scarabaeines–—a topology strongly incongruent with all other trees except for 50p-4.

### Diagnosis of new taxa

The 70p-1 tree (Figs. 1 and 2) was used to identify tribal-level clades and consequently selected for visualisation of phylogenetic relationships because its topology was more consistent with previous studies (Gunter et al., 2026; Lopes et al., 2024; Mlambo et al., 2015; Tarasov and Génier, 2015; Tarasov and Dimitrov, 2016). In this section we summarise the new tribal treatments, which are further expanded in the Taxonomy section.

Most of the previously described, monophyletic Afro-Eurasian tribes have been characterised in detail in recent studies (Daniel et al., 2020b; Davis et al., 2019; Rossini et al., 2022; Scholtz et al., 2025; Tarasov, 2017; Tarasov and Dimitrov, 2016) and were therefore not the primary focus of our analyses. However, we provide updated diagnoses of Gymnopleurini (Fig. 13), Onitini (Fig. 19) and Scarabaeini (Fig. 23), three historically recognised groups that have not been revised recently, and we enrich the diagnosis of Elassocanthonini by emphasising several informative characters (endophallites composed solely of A and BSc sclerites; epipleuron very wide) not mentioned in the original descriptions. Moreover, we merge Endroedyolini and Odontolomini, establishing Endroedyolini *sensu novo* composed of the subtribes Endroedyolina and Odontolomina *stat. nov.* (Fig. 12). This choice is justified by their marked morphological similarity and relatively shallow phylogenetic divergence.

As to the genus *Haroldius*, its fully supported sister relationship with Elassocanthonini+Endroedyolini *sensu novo* justifies its placement in Haroldiini *trib. nov.* (Fig. 14). This taxon is well-characterised by the presence of a wide epipleuron, an endophallus comprising only A and BSc sclerites, eight antennomeres, as well as a very peculiar ecology and disjunct Afro-Indomalayan distribution. Additionally, examination of the external morphology of the monotypic genus *Phaedotrogus* Paulian, 1985 indicated that it is hardly distinguishable from *Haroldius*, with which it is synonymised. Elassocanthonini, Endroedyolini *sensu novo* and Haroldiini share two synapomorphies: the presence of only A and BSc endophallites and a very broad epipleuron. The same endophallic configuration is known only in the closely related Coptorhinini and Paraphytini, representing a non-homoplastic synapomorphy supporting the monophyly of these five tribes, consistently with molecular evidence. We informally refer to this clade as Coptorhinomorpha, from *Coptorhina*, one of its most emblematic genera, and -morpha (from Greek morphē, “form, shape”).

The *incertae sedis* genus *Pedaria* is placed in Coprini *sensu novo* together with *Onychothecus* and *Copris*-related genera. Although this placement is strongly supported by our phylogenetic analyses, no clear morphological synapomorphies diagnose the tribe as a whole. To accommodate this disparity, we subdivide Coprini into three subtribes: Coprina, Onychothecina *subtr. nov.*, and Pedariina *subtr. nov.* (Figs. 9 and 10).

The genus *Circellium* emerged as a fully supported sister of Scarabaeini in most phylogenies. Evaluation of its morphology indicates that it shares several characters with that tribe, namely the absence of protarsi, the shape of endophallites and aedeagus, and the elytral striation. Nevertheless, the two lineages appear morphologically distinct in other characters of the head and legs, and they show substantial phylogenetic divergence. Therefore, we place *Circellium* in Circelliini *trib. nov*.

Consistent with our molecular phylogeny, Onthophagini and Oniticellini cannot be reliably distinguished on morphological grounds and share multiple homoplastic synapomorphies (Fig. 20). We therefore treat them as a single tribe, Onthophagini *sensu novo*, comprising Helictopleurina *stat. nov.* (formerly a subtribe of Oniticellini), Oniticellina *stat. nov.*, and Onthophagina. Neither phylogenetic nor morphological evidence supports the recognition of the remaining former subtribes of Oniticellini, Attavicinina, Drepanocerina, and Liatongina, which are here synonymized with Oniticellina.

Phylogenetic analyses revealed a further 11 previously unrecognised tribal-level clades. Many are represented by single, relictual, and phylogenetically isolated genera with distinctive morphologies. For example, *Janssensantus* (Janssensantini *trib. nov.*; Fig. 16) and *Bohepilissus* (Bohepilissini *trib. nov.*; Fig. 5) are readily diagnosed by unique combinations of external and genital characters. Likewise, *Tanzanolus* (Tanzanolini *trib. nov.*; Fig. 24) is distinguished by its unique wing venation. In contrast, *Nesovinsonia* (Nesovinsoniini *trib. nov.*; Fig. 18) lacks obvious morphological autapomorphies and is diagnosed by a combination of homoplastic characters. Although Tanzanolini and Nesovinsoniini form a well-supported sister-group relationship, they share no apparent morphological synapomorphies. Another isolated lineage is the monogeneric tribe Panelini Lewis, 1895 *stat. rev.* and *sensu novo*, originally established for a heterogeneous assemblage of genera but here revalidated and redefined to include only *Panelus*. Although superficially similar to other small-bodied former canthonines, including Bohepilissini, Tanzanolini, and the mentophiline *Lepanus* Balthasar, 1966, *Panelus* is readily distinguished by characters of wing venation, elytral striation, and the endophallus.

As to the genera *Heliocopris* and *Synapsis*, morphology corroborates the sister relationship found by UCEs. The new tribe Heliocoprini *trib. nov.* (Fig. 15) is well delimited by the presence of three elytral striae (8–10) placed on the pseudoepipleuron, a feature present elsewhere only in the Australasian Boletoscapterini and Mentophilini.

Chalconotini *trib. nov.*, comprising *Canthodimorpha*, *Chalconotus* and *Gyronotus*, is diagnosed on a combination of elytral, genital and leg features. Although *Canthodimorpha* was not included in the present analyses, previous phylogenetic studies consistently recover it as the sister genus of *Chalconotus* (see Taxonomy).

A strongly supported molecular clade comprises *Catharsius*+*Copridaspidus*+*Metacatharsius*, *Dwesasilvasedis*, *Hammondantus*+*Pycnopanelus*, *Macroderes*+*Xinidium*, Onitini and Onthophagini *sensu novo* (Fig. 2). Within this group, *Catharsius* and allies form the earliest-branching clade, the Catharsiini *trib. nov.* (Fig. 6), well-characterised by a transversely carinated mesanepisternum, wing venation and similar elytral striation. Reciprocal relationships between the other lineages vary depending on the analysis, which, together with their significant degree of morphological differentiation, justifies their characterisation as separate tribes: Dwesasilvasedini *trib. nov.* for *Dwesasilvasedis* (Fig. 11), Macroderini *trib. nov.* for *Macroderes* and *Xinidium* (Fig. 17), and Pycnopanelini *trib. nov.* for *Pycnopanelus* and *Hammondantus* (Fig. 22). Pycnopanelini share a very similar shape of parameres, the elytral striation pattern, the presence of uncus and the body shape. *Macroderes* and *Xinidium*, while differing in the presence/absence of pseudoepipleural carina and strong sexual dimorphism, share features of parameres and endophallites that make the diagnosis of the tribe sufficiently robust. Here, we name the clade formed by these six tribes Onthophagomorpha, after *Onthophagus*, its most species-rich genus. The only clear synapomorphy of the group is the presence of unci, which are found in many, especially most basally-derived genera of Onthophagini (*Phalops*, *Digitonthophagus*, *Kurtops*, *Hamonthophagus*, *Proagoderus*, etc.) and in all species of the other tribes. Moreover, two main groups can be identified within Onthophagomorpha based on elytral morphology, corresponding to distinct lineages in the tree 70p-1. One comprises Dwesasilvasedini, Macroderini and Onitini and is characterised by 9 or 10 elytral striae and, except in *Xinidium* and a few onitines, presence of a pseudoepipleural carina between stria 8 and 9. The second group comprises Onthophagini and Pycnopanelini, which have a similar number of striae: 9 in Pycnopanelini, with the 8th either reduced or, in *Pycnopanelus parvicollis*, completely missing; and 8 in Onthophagini, although some *Phalops* have a trace of an additional stria very likely homologous to the 8th of the Pycnopanelini. Both tribes completely lack a pseudoepipleural carina. Lastly, Catharsiini, which are basal to the remaining onthophagomorphans, have elytra with 9 or 10 striae and various degrees of development of the pseudoepipleural carina between stria 8 and 9. Onthophagomorpha probably also includes *Aphengoecus*, a morphologically distinct genus here placed in Aphengoecini *trib. nov.* (Fig. 4). Although *Aphengoecus*’ morphology provides little evidence of affinities with other tribes, previous molecular studies strongly support its placement within Onthophagomorpha (see Taxonomy), to which it is tentatively assigned.

Another well-supported molecular clade (clade A; Fig. 1) comprises Epactoidini, Epirinini, Sisyphini, and, depending on the topology, Bohepilissini and Panelini. It probably also includes Parachoriini and possibly Eurysternini (see Discussion). As its morphological diagnosis remains unclear and its composition uncertain, we do not formally name this clade.

Lastly, the Madagascan canthonines of the *Nanos* and *Epilissus* groups of genera, each represent distinct evolutionary lineages. As their taxonomy requires separate treatment, they are not considered further here and will be revised in a separate study.

## TAXONOMY

### Aphengoecini *trib. nov*

(Figs. 4, 3F)

**Figure 3.**
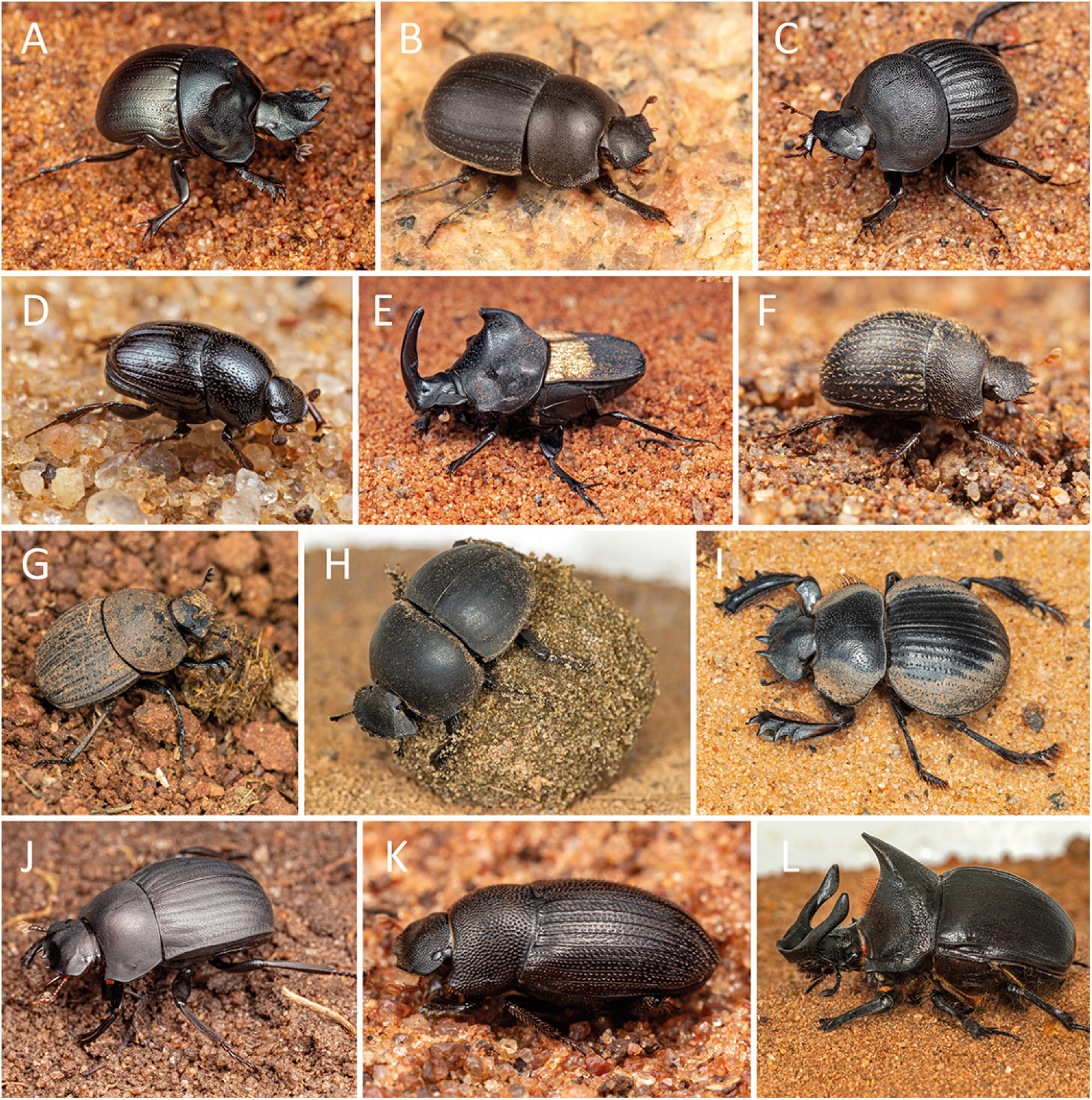
Live dung beetles representing different tribes. (A) *Coptorhina auspicata* Péringuey, 1901 (Coptorhinini); (B) *Versicorpus daures* Deschodt, 2019 (Elassocanthonini); (C) *Macroderes gifboomi* Abdalla & Deschodt, 2018 (Macroderini *trib. nov.*); (D) *Pycnopanelus krikkeni* Cambefort, 1978 (Pycnopanelini *trib. nov.*); (E) *Tragiscus dimidiatus* Klug, 1855 (Onthophagini Oniticellina *stat. nov.*); (F) *Aphengoecus multiserratus* Scholtz & Howden, 1987 (Aphengoecini *trib. nov.*); (G) *Epirinus asper* Péringuey, 1901 (Epirinini); (H) *Circellium bacchus* (Fabricius, 1781) (Circelliini *trib. nov.*); (I) *Pachysoma gariepinum* Ferreira, 1953 (Scarabaeini); (J) *Gyronotus mulanjensis* Davis, Scholtz & Harrison, 1999 (Chalconotini *trib. nov.*); (K) *Pedaria segregis* Péringuey, 1901 (Coprini Pedariina *subtrib. nov.*); (L) *Heliocopris andersoni* Bates, 1868 (Heliocoprini *trib. nov.*). Pictures courtesy of Hennie De Klerk.

**Type genus**: *Aphengoecus* Péringuey, 1901.

**Genus included:**

*Aphengoecus* Péringuey, 1901 (2 valid species; Afrotropical).

**Diagnosis**. The tribe is defined by the following unique combination of characters: 1) second elytral carina located on pseudoepipleuron (Fig. 4D); 2) first elytral carina and epipleural carina converging on posterior half (Fig. 4D); 3) medial margin of 2nd elytral carina adjoining stria 8 (Fig. 4D); 4) anterior head margin with 6 teeth (4 on clypeus, 1 on each gena) (Fig. 4A).

**Figure 4.**
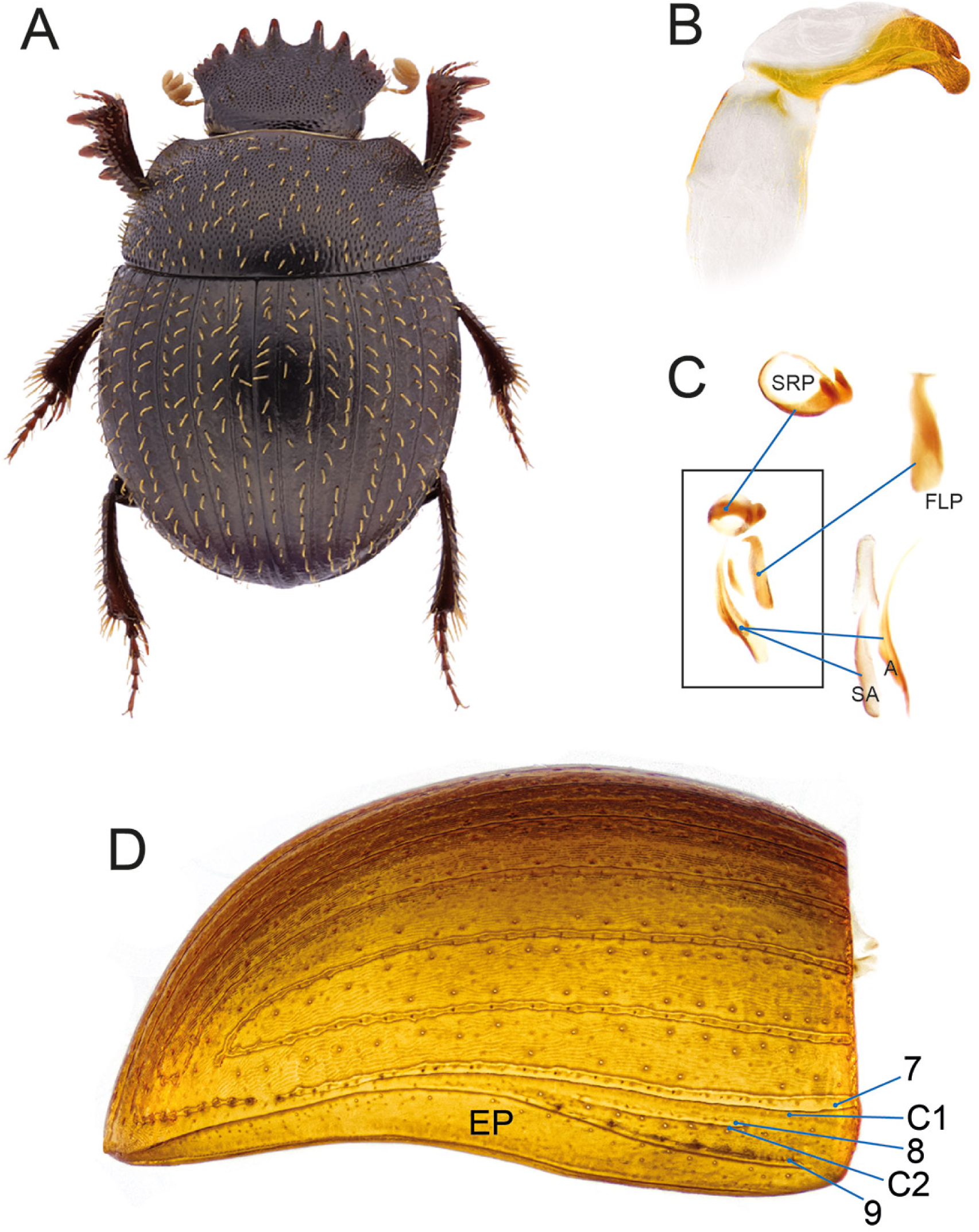
Diagnostic morphological characters of Aphengoecini *trib. nov*. *Aphengoecus multiserratus*: (A) habitus (picture by C. Deschodt); (B) aedeagus in left lateral view; (C) endophallites; (D) elytron showing lateralmost striae (7–9) and carinae (C1, C2; epipleural carina not marked).

**Description.**

Small sized beetles (4–5 mm); body oval, strongly convex; dorsal surface covered with short, yellowish brown setae.

**Table 3.**
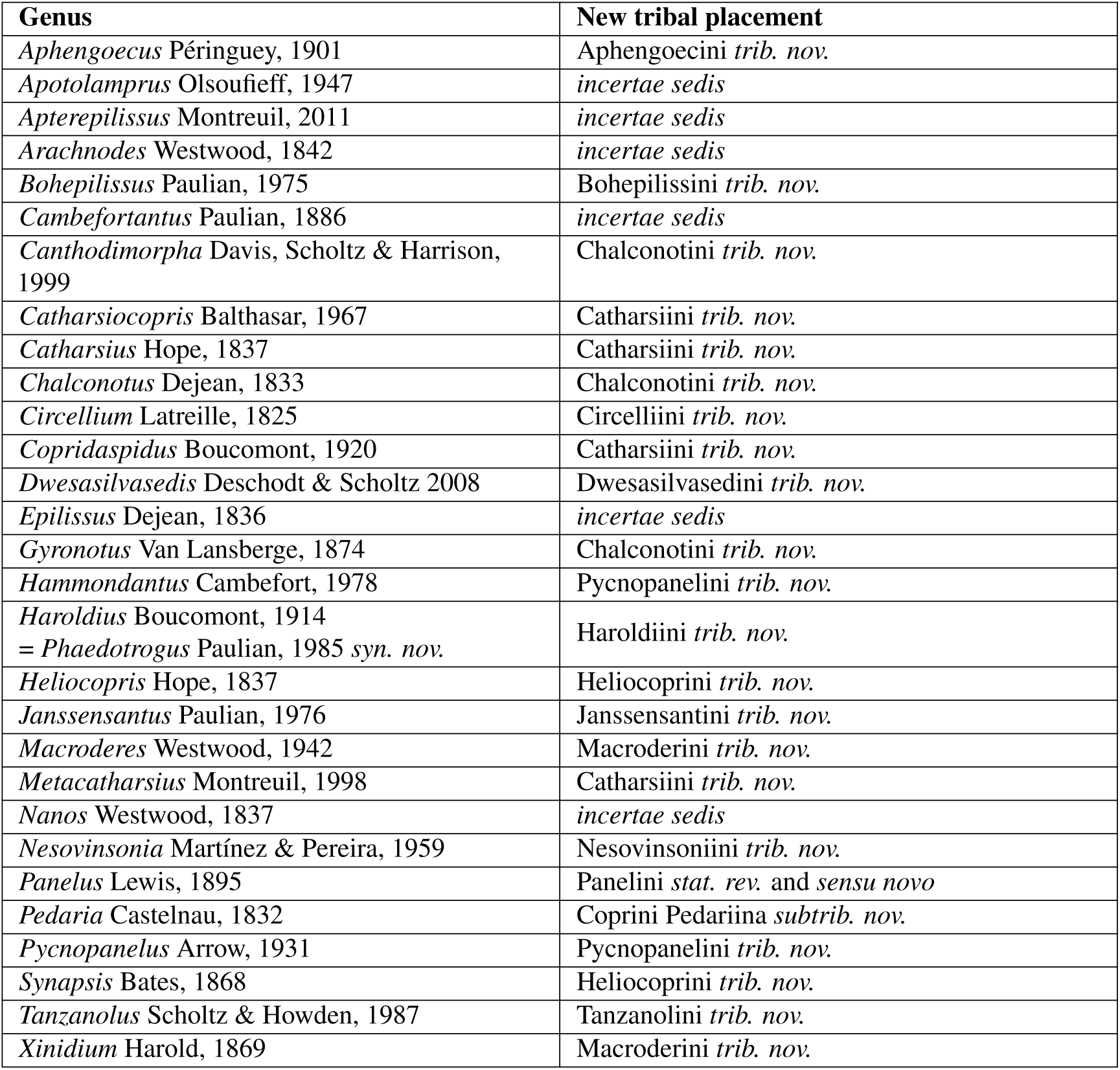
Revised tribal placement of Afro-Eurasian genera formerly treated as *incertae sedis*.

*Head*. Anterior margin of clypeus with 4 sharp teeth, genae with one sharp tooth; frontoclypeal sulcus absent. Eyes small. Antennae with 9 antennomeres.

*Pronotum*. Strongly and densely punctate; lateral pronotal carina entire.

*Elytra and wings*. Elytron with 9 visible striae; elytron carinate between striae 7–8 and 8–9; striae 8 and 9 placed on pseudoepipleuron, stria 8 and second elytral carina restricted to anterior half of elytron and not reaching elytral anterior margin; epipleuron wide, epipleural carina adjoining first elytral carina posteriorly. Mesoscutellum concealed. Metathoracic wings absent.

*Legs*. Protibia with 3 teeth with smaller denticles in between (number of denticles is species-specific); protibial distal end transversely truncated, male distoventral region feebly produced and bearing lamellalike setae. Meso- and metatibiae gradually expanded proximo-distally. Tarsal claws not toothed.

*Ventral body surface*. Anterior region of hypomeron depressed; anterior hypomeral carina stretching towards lateral margin of hypomeron; posterior longitudinal hypomeral carina absent.

*Tergite VIII*. Wider than long, evenly convex.

*Genitalia*. Parameres asymmetric (Fig. 4B). Endophallites (Fig. 4C): FLP present, elongated, simple; A and SA present; SRP ring-shaped.

**Distribution and biology**. *Aphengoecus* is restricted to deep coastal sands and coastal mountains in the Western Cape in South Africa. Its two species appear to be rather localised, with *A. clypeatus* Péringuey, 1901 being possibly extinct due to degradation of its original habitat. *A. multiserratus* Scholtz & Howden, 1987 is attracted to various excrements, meat and banana baits (Davis et al., 2008).

**Remarks**. *Aphengoecus* has long been considered a member of the Canthonini and later moved to *incertae sedis* by Tarasov and Dimitrov (2016). Unfortunately, we were not able to include this genus in our reconstruction due to the lack of material suitable for DNA analyses. However, most molecular phylogenies consistently recovered the genus as closely related to onthophagomorphan taxa: Mlambo et al. (2015) found *Aphengoecus* nested within Macroderini; Sole and Scholtz (2010) as sister to *Macroderes*+(*Dwesasilvasedis*+*Xinidium*); and Tarasov and Dimitrov (2016) and Tarasov (2017) as basal to all remaining Onthophagomorpha except Catharsiini. The only exception is Mlambo et al. (2014) who, based on three loci, found it as sister to Chalconotini.

We consider it unlikely that *Aphengoecus* is nested within Macroderini, which are united by a sound combination of genital and elytral morphological synapomorphies and which significantly differ from *Aphengoecus* in elytral carination, leg and head features, and symmetry of the parameres. *Aphengoecus* has several peculiar, apparently strongly derived characters (elytral striation and carination, absence of uncus, shape of clypeus) that make it difficult to further compare it with other Onthophagomorpha. Overall, it is likely that *Aphengoecus* falls somewhere between other onthophagomorphan tribes (Fig. 2), although for now its exact placement remains unclear. These observations strongly support the creation of Aphengoecini *trib. nov.* as a distinct tribe.

### Bohepilissini *trib. nov*

(Fig. 5)

**Figure 5.**
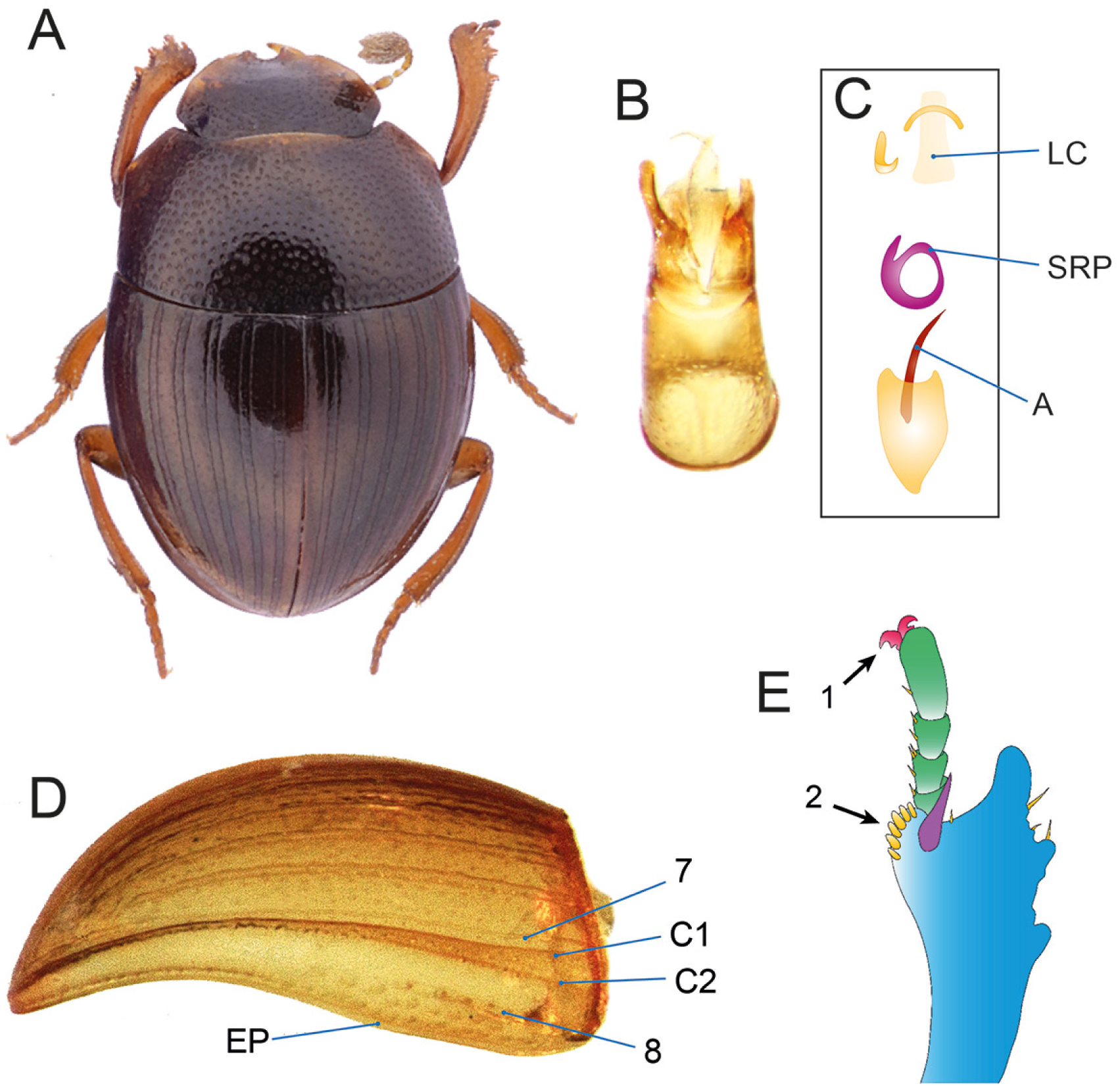
Diagnostic morphological characters of Bohepilissini *trib. nov*. (A) *Bohepilissus nitidus*, habitus (picture by C. Deschodt). *Bohepilissus* sp.: (B) aedeagus in dorsal view; (C) endophallites; (D) elytron showing lateral striae (7–8) and carinae (C1, C2; epipleural carina not marked); (E) male protibia showing toothed tarsal claws (arrow 1) and set of scale-like setae on distal margin (arrow 2).

**Type genus**: *Bohepilissus* Paulian, 1975.

**Genus included:**

*Bohepilissus* Paulian, 1975 (2 valid species; Afrotropical).

**Diagnosis**. The tribe is defined by the following unique combination of characters: 1) SA and FLP endophallites absent (Fig. 5C); 2) 1st elytral carina with two ridges (Fig. 5D); 3) male distoventral protibial apex processed and bearing lamella-like setae (Fig. 5E); 4) tarsal claws toothed (Fig. 5E).

**Description.**

Very small sized beetles (1.7–3.1 mm); body oval, moderately convex.

*Head*. Anterior margin of clypeus with two teeth, angulate laterally to teeth, genae slightly angulated; frontoclypeal sulcus absent. Antennae with 9 antennomeres.

*Pronotum*. Moderately convex, with sparse punctures; lateral pronotal carina entire.

*Elytra and wings*. Elytra with 8 visible striae, 8th stria reduced to anterior third of elytron (a hardly visible groove is present along the epipleuron, but it is unclear whether it is the trace of a 9th carina or not); elytron carinate laterally to stria 7, carina with two ridges; elytron slanted inwards laterally to carina; stria 8 placed on pseudoepipleuron. Mesoscutellum concealed. Metathoracic wings absent.

*Legs*. Protibia with 3 teeth; protibial distal end obliquely truncated, male distoventral region produced and bearing lamella-like setae; protarsus present. Meso- and metatibiae gradually expanded proximo-distally. Tarsal claws toothed.

*Ventral body surface*. Anterior region of hypomeron depressed; anterior hypomeral carina stretching towards lateral margin of hypomeron; posterior longitudinal hypomeral carina absent.

*Tergite VIII*. Wider than long, evenly convex.

*Genitalia*. Parameres either symmetrical or asymmetrical (Fig. 5B). Endophallites: LC and A present; FLP and SA absent; SRP ring-shaped.

**Distribution and biology**. The two described species of *Bohepilissus* are restricted to forest patches on the coastline and adjacent coastal mountains in southern South Africa (Davis et al., 2008, 2020). They seem to be leaf litter dwellers and are attracted to dung and carrion. Intriguingly, *Bohepilissus subtilis* Boheman, 1857 presents geographically fragmented and phenotypically divergent populations hinting at a strong, possibly species-level differentiation between them (Scholtz and Howden, 1987).

**Remarks**. *Bohepilissus* was formerly classified within the polyphyletic Canthonini (Scholtz and Howden, 1987) or Epilissini (Montreuil, 2010; currently deemed as an invalid tribe by Gunter et al. (2026)), and later placed in *incertae sedis* by Tarasov and Dimitrov (2016). Its phylogenetic placement is still enigmatic, as there is inconsistency among available phylogenies (Mlambo et al., 2014; Sole and Scholtz, 2010; Tarasov and Dimitrov, 2016; Tarasov and Génier, 2015). Our reconstructions mostly found it in various positions within or sister to the clade Epactoidini+(Sisyphini+Epirinini) or close to Panelini. Therefore, while it is likely that *Bohepilissus* is strictly related to these lineages, its exact placement was not reliably solved by our analyses.

Morphologically, the genus looks close to *Tanzanolus* (Tanzanolini) and *Panelus* (Panelini). All these genera include very small, relictual Afrotropical forest dwellers with oval-shaped body and digitiform protibial apex. In particular, *Tanzanolus* and *Bohepilissus* have a somewhat similar elytral configuration with two elytral carinae (yet a different number of striae) and reduced endophallites, with *Bohepilissus* lacking FLP and SA and *Tanzanolus* lacking all but the axial endophallite. Other characters set them apart, such as toothed (*Bohepilissus*) *vs*. not toothed (*Tanzanolus*) tarsal claws, absence (*Bohepilissus*) *vs*. presence (*Tanzanolus*) of posterior longitudinal hypomeral carina, elytral carinae fused posteriorly in *Bohepilissus* but not in *Tanzanolus*, and the shallowly (*Bohepilissus*) vs. clearly (*Tanzanolus*) 6-toothed head margin. Moreover, no phylogeny supports a close relationship between the two genera. *Panelus* may be more phylogenetically close to *Bohepilissus* based on molecular results, yet their elytral and genital morphologies are quite different. Overall, the phylogenetic isolation and morphological distinctiveness of *Bohepilissus* fully supports its accommodation in Bohepilissini *trib. nov*.

### Catharsiini *trib. nov*

(Fig. 6)

**Figure 6.**
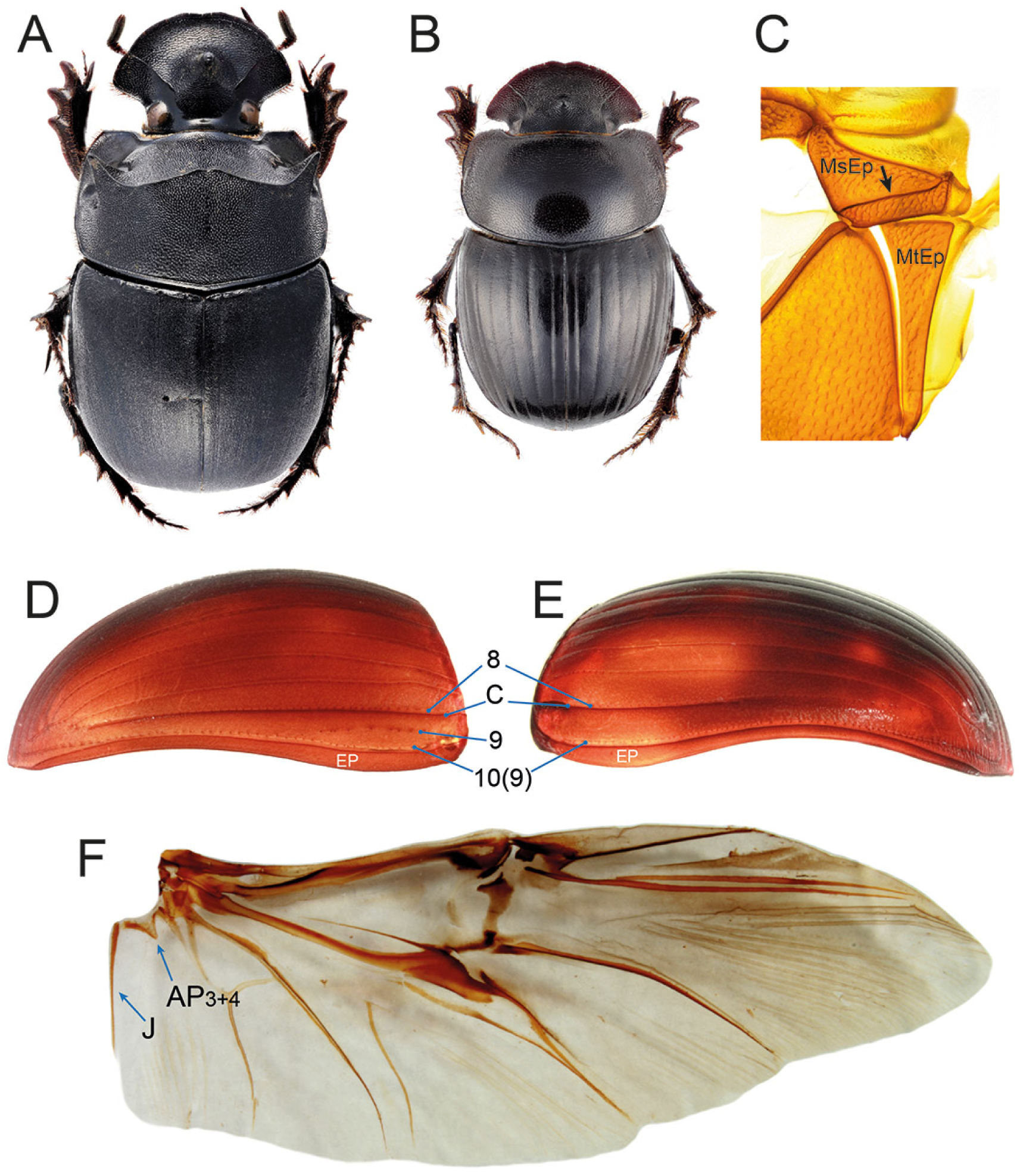
Diagnostic morphological characters of Catharsiini *trib. nov*. (A) *Catharsius molossus*, habitus; (B) *Metacatharisus* sp., habitus; (C) *Metacatharsius* sp., mesanepisternum and metanepisternum with arrow pointing at mesanepisternal carina; (D) *Metacatharsius* sp. and (E) *Catharsius* sp., elytra showing lateral striae (8–10, stria 10 of *Metacatharsius* homologous to 9 of *Catharsius*) and carina (C; epipleural carina not marked); (F) *Metacatharsius* sp., metathoracic wing showing well developed J and poorly developed AP3+4 veins.

**Type genus**: *Catharsius* Hope, 1837.

**Genera included:**

*Catharsiocopris* Balthasar, 1967 (1 valid species; Afrotropical);

*Catharsius* Hope, 1837 (107 valid species; Afrotropical, Indomalayan, Palaearctic);

*Copridaspidus* Boucomont, 1920 (1 valid species; Afrotropical);

*Metacatharsius* Montreuil, 1998 (65 valid species; Afrotropical, Palaearctic).

**Diagnosis**. The tribe is defined by the following unique combination of characters: 1) medial margin of 1st elytral carina adjoining stria 8 (Fig. 6D–E); 2) wing with J vein well developed and AP3+4 reduced (Fig. 6F); 3) mesanepisternum carinated dorsoventrally (Fig. 6C).

**Description.**

Small to very large sized beetles (4.8–44.6 mm); body from moderately flattened to very convex.

*Head*. Anterior margin of clypeus with or without 2 teeth; frontoclypeal sulcus present, carinated, tuberculated or horned. Antennae with 9 antennomeres.

*Pronotum*. Densely granulated or punctured, often with protrusions or carinae, particularly in *Catharsius*; mediolateral regions with a fovea; lateral pronotal carina entire, sometimes doubled posteriorly.

*Elytra and wings*. Elytra with 9 or 10 visible striae; elytron carinate immediately laterally to stria 8, carina more or less effaced posteriorly. Mesoscutellum concealed. Metathoracic wings present; J vein well developed and AP3+4 reduced.

*Legs*. Protibia with 3 or 4 teeth; protibial distal end obliquely truncated; male uncus present or absent. Meso- and metatibiae strongly expanded proximo-distally, dorsal surface with a transverse carina. Tarsal claws not toothed.

*Ventral body surface*. Anterior hypomeral carina present or not, if present stretching towards lateral margin of pronotum; posterior longitudinal hypomeral carina absent; mesanepisternum with a dorsoventrally oriented carina.

*Tergite VIII*. Wider than long, evenly convex, longer in males.

*Genitalia*. Parameres symmetrical, elongated, thin and sharp posteriorly. Endophallites: A, SA, FLP and SRP present; LC present.

**Distribution and biology**. The Catharsiini are a species-rich group of tunnellers widespread in tropical regions of Afro-Eurasia. *Catharsius* is particularly diverse in the Afrotropical realm, with several species found in the Indomalayan realm, while *Metacatharsius* is largely restricted to the Afrotropics, with only one species extending to the Palaearctic. *Catharsius* species inhabit rainforests to arid habitats and often exploit various types of food sources, from dung to carrion. *Metacatharsius* are more dry-adapted, being found from savanna to desert, and also show associations biased to either dung or carrion (Davis et al., 2008).

**Remarks**. Historically, *Catharsiocopris*, *Catharsius*, *Copridaspidus* and *Metacatharsius* were considered Coprini, mainly due to the presence of a transverse carina on the dorsal surface of the metatibia, one of the most important features defining the historical concept of the tribe (Balthasar, 1963; Davis et al., 2008; Montreuil, 1998). More recently, Tarasov and Dimitrov (2016) moved them—with the exception of *Catharsiocopris*, which was not treated—to *incertae sedis*, together with several other former coprine genera not strictly related to *Copris*. The need for the establishment of Catharsiini was proposed, yet not published, in the revision of Afrotropical *Catharsius* by Takano (2018).

The monophyly of Catharsiini as defined here is fully supported both by morphology (Tarasov and Génier, 2015; Montreuil, 1998) and by our molecular analyses, which are in substantial agreement with previous research (Tarasov and Dimitrov, 2016; Gunter et al., 2026; Lopes et al., 2024; Takano, 2018). Our trees support the placement of the monotypic genus *Copridaspidus* within *Catharsius* (Fig. 2), while we were not able to sample *Catharsiocopris*, also monotypic. However, both genera were found to be undoubted junior synonyms of *Catharsius* by Takano (2018). Therefore, although we retain them as valid taxa here, we expect them to be synonymised soon in light of more comprehensive morphological and phylogenetic analyses.

As to higher-level relationships, Catharsiini appear to be basal to the Onthophagomorpha clade (Fig. 2; see also Tarasov and Génier (2015) and Tarasov and Dimitrov (2016)), which include other uncus-bearing tribes of African origin (see Discussion).

### Chalconotini *trib. nov*

(Figs. 7, 3J)

**Figure 7.**
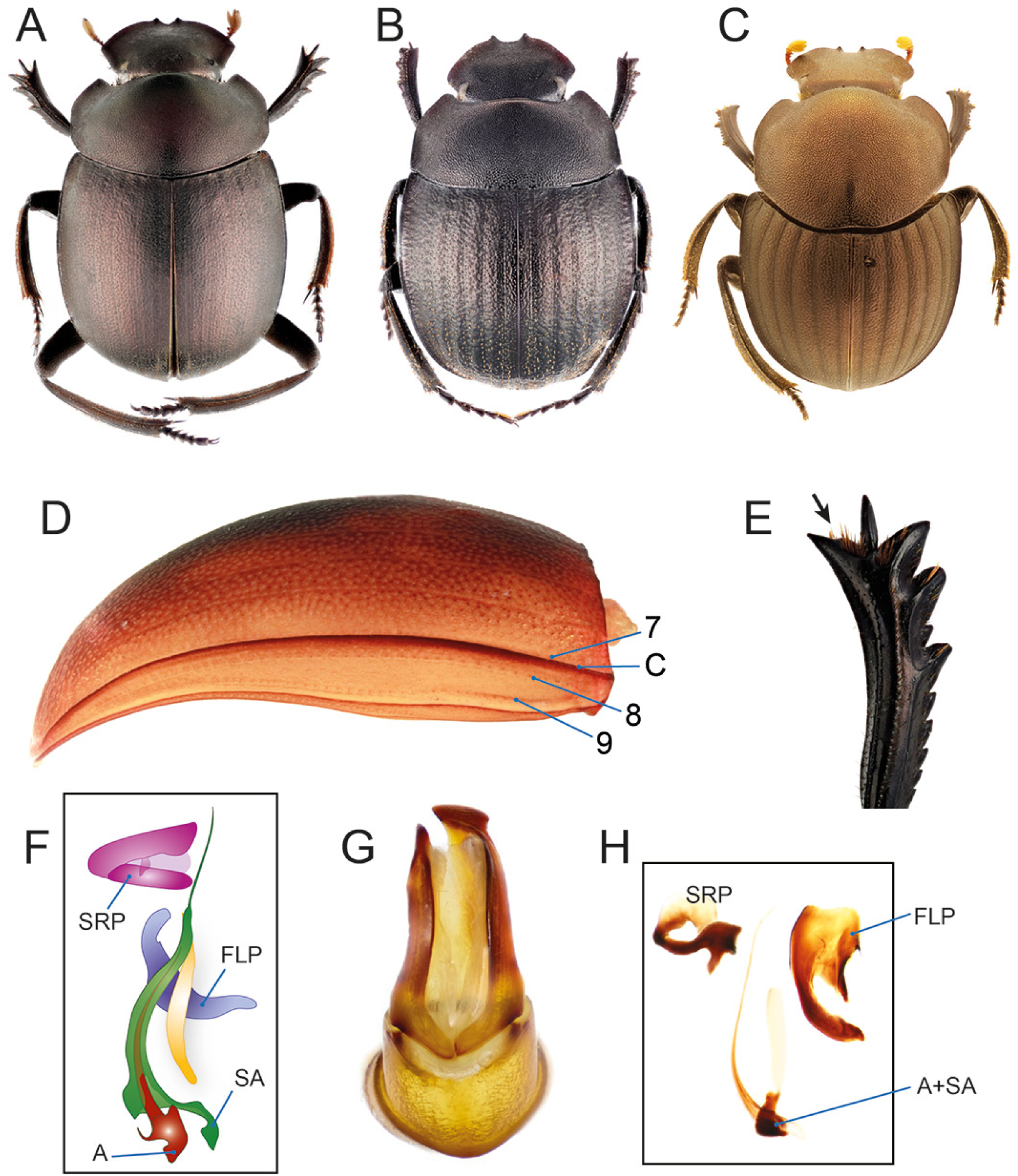
Diagnostic morphological characters of Chalconotini *trib. nov*. (A) *Chalconotus* sp., habitus; (B) *Gyronotus* sp., habitus; (C) *Canthodimorpha lawrencei*, habitus (picture by C. Deschodt); (D) *Chalconotus convexus*, elytron showing lateral striae (7–9) and carina (C; epipleural carina not marked); (E) *Chalconotus cupreus*, female protibia showing absence of protarsus (black arrow); (F) *C. convexus*, endophallites; (G) *Gyronotus* sp., aedeagus in dorsal view; (H) *Gyronotus* sp., endophallites.

**Type genus**: *Chalconotus* Dejean, 1833.

**Genera included:**

*Canthodimorpha* Davis, Scholtz & Harrison, 1999 (1 valid species; Afrotropical);

*Chalconotus* Dejean, 1833 (10 valid species; Afrotropical);

*Gyronotus* Lansberge, 1874 (17 valid species; Afrotropical).

**Diagnosis**. The tribe is defined by the following unique combination of characters: 1) SA endophallite tightly wrapped within axial endophallite, the two almost inseparable but recognizable (Figs. 7F, 7H); 2) protarsus absent both in males and females (Fig. 7E); 3) elytron with 9 visible striae (Fig. 7D); 4) presence of a single elytral carina lateral to stria 7 (Fig. 7D); 5) male uncus well developed.

**Description**.

Medium to very large sized beetles (11–35 mm); body oval (*Chalconotus*, *Canthodimorpha*, *Gyronotus*) to elongated (*Gyronotus*), convex (*Chalconotus*, *Canthodimorpha*, *Gyronotus*) to flattened (*Gyronotus*).

*Head*. Surface evenly, often densely punctate. Anterior margin of clypeus with two teeth, head margin evenly rounded or slightly sinuate at genoclypeal suture; frontoclypeal sulcus absent. Antennae with 9 antennomeres.

*Pronotum*. Strongly convex to flattened, in male *Canthodimorpha* and some species of *Chalconotus* more or less strongly produced anteriorly; evenly covered with simple punctures; lateral pronotal carina entire.

*Elytra and wings*. Elytron with 9 visible striae; elytron carinate laterally to stria 7, carina with one ridge; elytron slanted inwards laterally to carina; striae 8 and 9 placed on pseudoepipleuron. Mesoscutellum concealed. Metathoracic wings present (*Chalconotus*) or absent (*Canthodimorpha*, *Gyronotus*); if present, RP1 with wide posterior sclerite.

*Legs*. Protibia with 3 teeth, distal margin oblique; in males, uncus well developed, rounded to sharp; protarsus absent in both sexes. Meso- and metatibiae poorly and evenly expanded proximo-distally. Tarsal claws not toothed.

*Ventral body surface*. Anterior hypomeral carina absent or extremely reduced (some *Gyronotus*) or present and stretching towards lateral margin of hypomeron; posterior longitudinal hypomeral carina absent. Metaventrite evenly convex; mesometaventral sulcus strongly rounded to transverse.

*Tergite VIII*. Evenly convex, longer in males; medial region swollen ventrally to ventral sulcus in male *Chalconotus* and some *Gyronotus*.

*Genitalia*. Parameres either symmetrical (*Canthodimorpha*) or asymmetrical (*Chalconotus*, *Gyronotus*; Fig. 7G). Endophallites (examined only in *Gyronotus* and *Chalconotus*): SA endophallite tightly wrapped within axial endophallite, the two almost inseparable but recognizable; FLP and SRP present; LC absent.

**Distribution and biology**. The tribe is endemic to the Afrotropical realm. *Chalconotus* is widespread throughout the entire realm and inhabits both forest and savannah habitats, with a few species being very common. *Gyronotus* and *Canthodimorpha* show relictual distributions in forests from eastern to southern Africa and in coastal shrublands of Mozambique, respectively (Davis, 1999; Davis et al., 2020).

**Remarks**. Chalconotini *trib. nov*. is a morphologically rather homogeneous group composed of former Canthonini members. Its monophyly is strongly supported both by morphology and molecules. Tarasov and Génier (2015) found evidence for a sister relationship between *Chalconotus* and *Gyronotus* based on both external and genital phenotypic characters. This was corroborated by several molecular studies based both on nuclear and mitochondrial genes (Mlambo et al., 2014; Sole and Scholtz, 2010; but see also Mlambo et al., 2015; Tarasov and Dimitrov, 2016), and is finally confirmed with full support by our results (Fig. 1). Although we were not able to include *Canthodimorpha* in our molecular analyses, the genus is consistently recovered as sister to *Chalconotus* across various studies (Mlambo et al., 2014, 2015; Tarasov and Dimitrov, 2016; Sole and Scholtz, 2010), which is broadly confirmed by morphology.

As to phylogenetic relationships between Chalconotini and other tribes, UCEs-based phylogenies consistently recovered a sister relationship with Scarabaeini+Circelliini (most of our trees; Gunter et al., 2026; Lopes et al., 2024). The three tribes share several features: the absence of protarsi in both sexes, a very similar elytral striation and (mostly) asymmetric parameres, strongly corroborating the hypothesis of their close affinity.

### Circelliini *trib. nov*

(Figs. 8, 3H)

**Figure 8.**
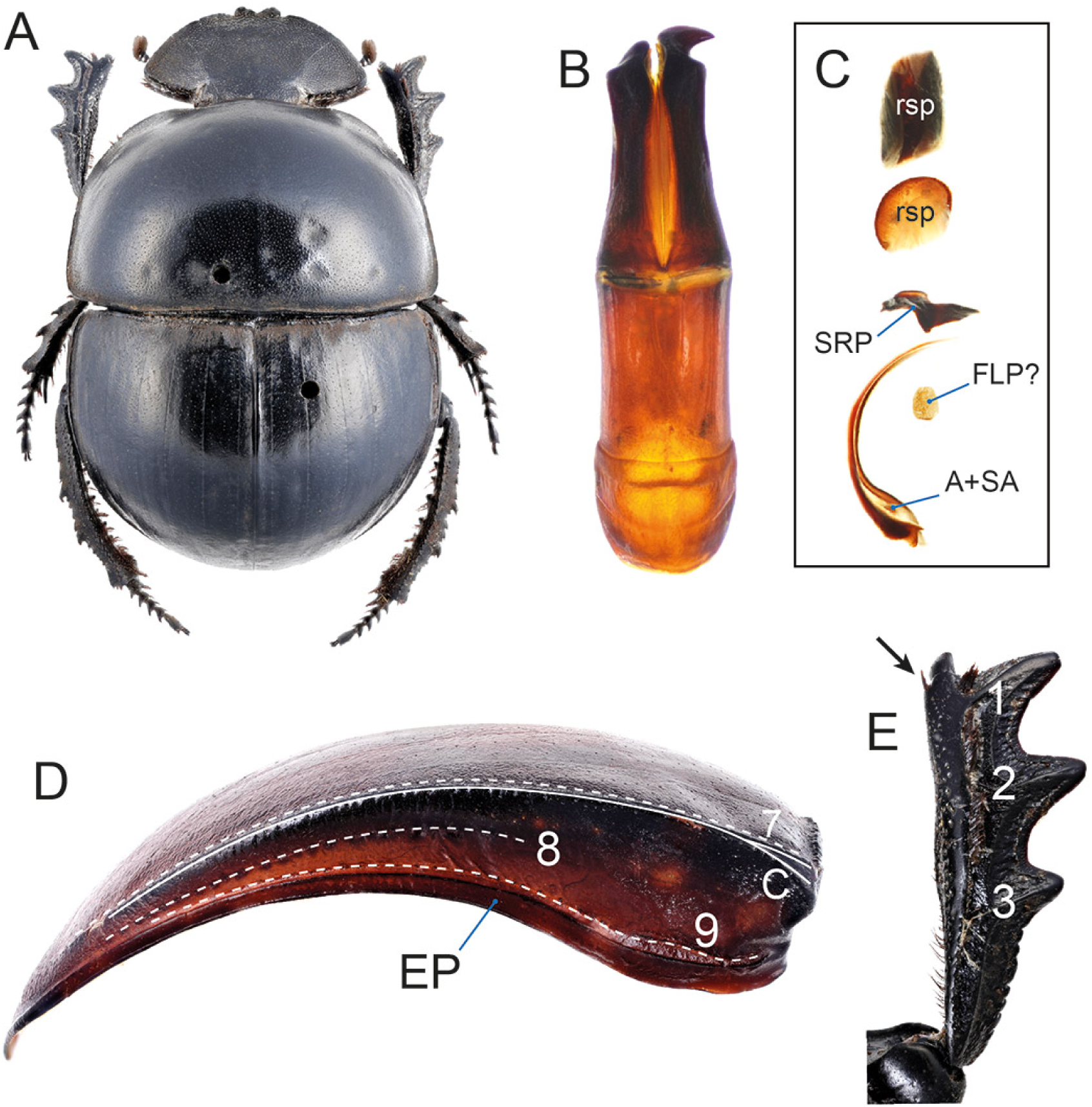
Diagnostic morphological characters of Circelliini *trib. nov.* *Circellium bacchus*: (A) habitus; (B) aedeagus in dorsal view; (C) endophallites; (D) elytron showing lateral striae (7–9) and carina (C); (E) protibia showing number of lateral teeth and absence of tarsus (black arrow).

**Type genus**: *Circellium* Latreille, 1829.

**Genus included:**

*Circellium* Latreille, 1829 (1 valid species; Afrotropical).

**Diagnosis**. The tribe is defined by the following combination of characters: 1) elytron with 9 striae (Fig. 8D); 2) elytron carinated between striae 7–8 (Fig. 8D); 3) protarsus absent in both sexes (Fig. 8E); 4) protibia with 3 teeth (Fig. 8E); 5) subaxial and axial endophallites as long as half of endophallus (Fig. 8C).

**Description.**

Very large sized beetles (22.0–50.0 mm); body very convex, black.

*Head*. Clypeal margin with 2 teeth, sharply notched lateral to teeth; head surface devoid of carinae or protrusions; frontoclypeal sulcus absent. Antennae with 9 antennomeres.

*Pronotum*. Evenly convex, without protrusions; finely punctate dorsally, rugose laterally; lateral pronotal carina present.

*Elytra and wings*. Elytron short, approximately as long as 1.3 times length of pronotum; with 9 striae; carinated between striae 7 and 8. Mesoscutellum concealed. Metathoracic wings absent.

*Legs*. Protibia with 3 teeth; uncus absent; protarsi absent in both sexes. Meso- and metatibiae feebly expanded proximo-distally, dorsal surface with two denticles. Tarsal claws not toothed.

*Ventral body surface*. Anterior hypomeral carina absent; posterior longitudinal hypomeral carina absent. Metaventrite not bulging anteriorly.

*Tergite VIII*. Evenly convex, subtriangular.

*Genitalia*. Parameres asymmetrical. Endophallites: LC absent; A and SA endophallites present, as long as half of endophallus; putative FLP reduced to a small bristle-like sclerite; SRP ring-shaped; presence of two raspulae distally.

**Distribution and biology**. The only species in the tribe, *Circellium bacchus* (Fabricius, 1781), is restricted to the southern coast and hinterland of South Africa, between Port Elizabeth (Gqeberha) and Hermanus, where it lives in shrublands and open vegetation (Davis et al., 2008). It is a roller feeding on dung of various herbivores and, due to its fragmented populations especially in the western part of its range, it is currently classified as vulnerable by the IUCN Red List (Davis et al., 2020).

**Remarks**. *Circellium bacchus*, originally described in the genus *Scarabaeus*, was initially placed in the Scarabaeini due to the morphological resemblance with that tribe (Janssens, 1938; Ferreira, 1972). Later, it was moved to the Canthonini mainly due to the 3-toothed protibia (Cambefort, 1978; Scholtz and Howden, 1987), and eventually to *incertae sedis* (Tarasov and Dimitrov, 2016).

Both molecules (present analyses; Tarasov and Dimitrov, 2016) and morphology (Tarasov and Génier, 2015) fully support a sister relationship of *Circellium* with Scarabaeini. In fact, the two lineages share several morphological characters, some of which probably represent true synapomorphies: 1) absence of protarsi in both sexes; 2) elytron with 9 striae, striae 8–9 placed on pseudoepipleuron; 3) endophallus with very elongated A and SA endophallites, FLP often reduced and presence of raspulae; 4) asymmetrical aedeagus. On the other hand, the two groups show several differences: the shape of clypeus and genae (with 2 and 0 teeth, respectively, in *Circellium*; mostly with 4 and 2 teeth in Scarabaeini); the number of protibial teeth (3 in *Circellium*, 4 in *Scarabaeini*); and the presence of an elytral carina between striae 8 and 9 (absent in *Circellium*; present in Scarabaeini). Moreover, *Circellium* has a more convex, almost hemispherical body shape, while Scarabaeini tend to be more flattened.

The divergence between the two lineages is deep, exceeding that observed between closely related tribes such as Epirinini and Sisyphini or Elassocanthonini, Endroedyolini, and Haroldiini. Moreover, *Circellium* is biogeographically unique, being a monotypic relict genus occurring only in southern South Africa. Overall, this flagship genus is well distinguished from Scarabaeini under several perspectives, and we accommodate it in its own tribe, Circelliini *trib. nov*.

### Coprini Leach, 1815

(Figs. 9, 10)

**Figure 9.**
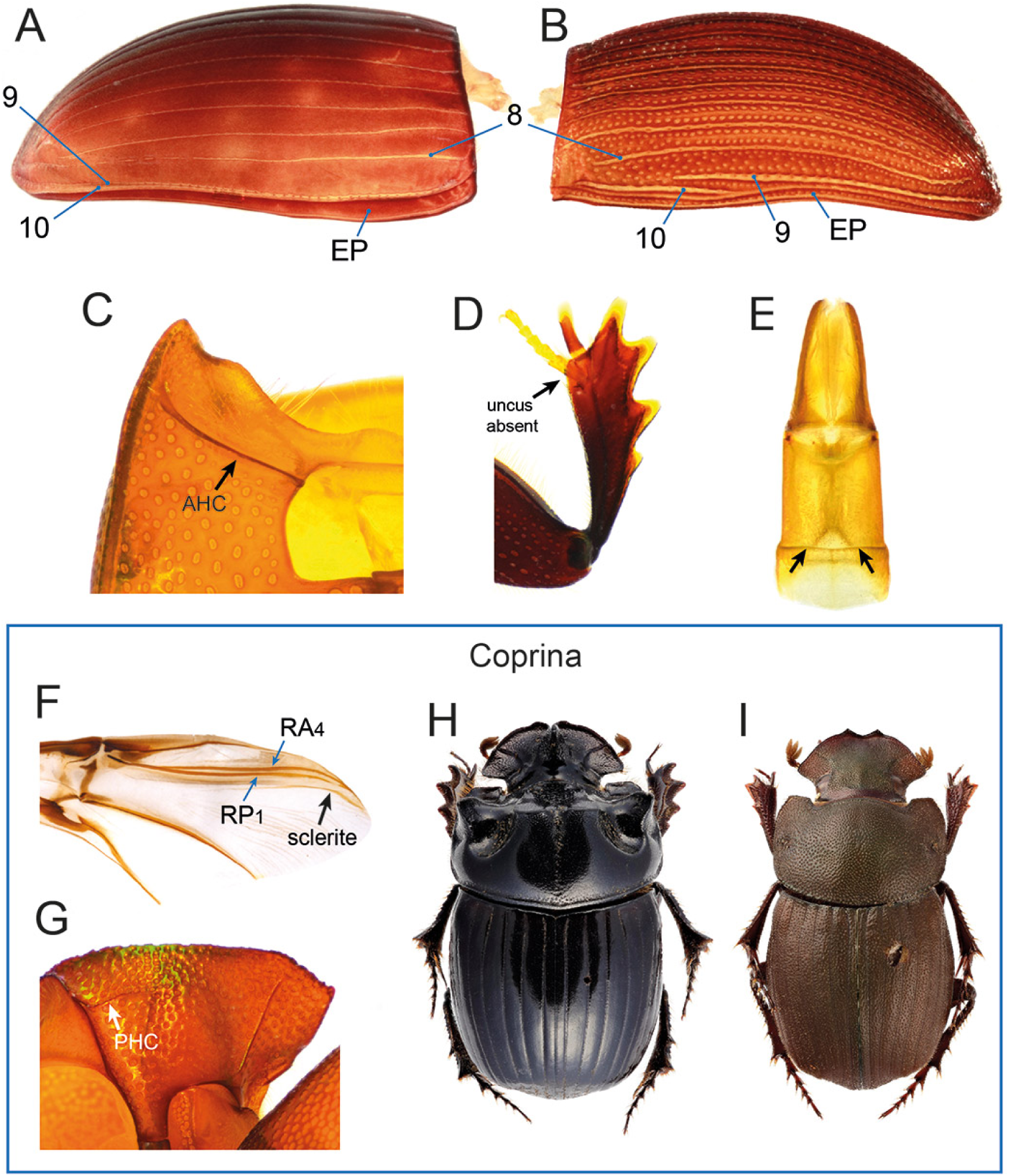
Diagnostic morphological characters of Coprini and Coprina. Coprini: (A) *Copris* sp. And (B) *Pedaria* sp., elytra with enumeration of lateral striae (8–10); (C) *Microcopris* sp., hypomeron showing anterior hypomeral carina; (D) *Copris* sp., protibia showing absence of uncus; (E) *Copris* sp., aedeagus in dorsal view showing absence of tubercles (black arrows). Coprina: (F) *Copris* sp., metathoracic wing showing posterior sclerite of RP1 vein; (G) *Pseudopedaria grossa*, hypomeron showing posterior hypomeral carina; (H) *Copris lunaris*, habitus; (I) *P. grossa*, habitus.

**Figure 10.**
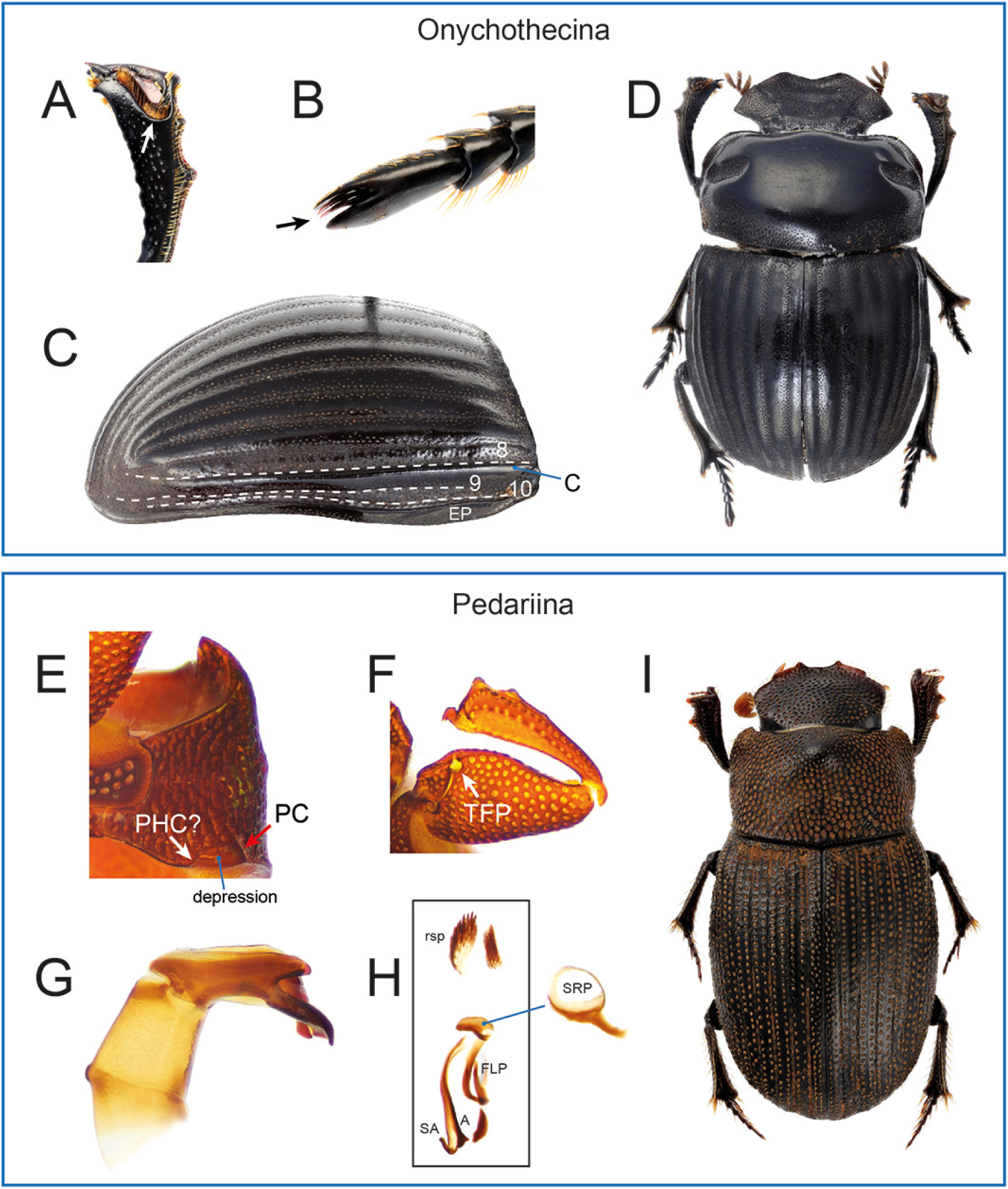
Diagnostic morphological characters of Onychothecina *subtrib. nov.* and Pedariina *subtrib. nov.* Onychothecina, *Onychothecus tridentigeris*: (A) protibia, arrow pointing at depression accommodating protarsus; (B) detail of mesotarsus, arrow pointing at claws protected by expansion of 5th tarsomere; (C) elytron showing lateral striae (8–10) and carina (C; epipleural carina not marked); (D) habitus of male. Pedariina: (E) *Pedaria alternans*, hypomeron showing the reduced pronotal carina and the putative posterior longitudinal hypomeral carina flanked by a depression; (F) *P. alternans*, ventral view of proleg showing trochanterofemoral pit; (G) *P. alternans*, aedeagus in left lateral view; (H) *Pedaria* sp., endophallites; (I) *P. biseria*, habitus.

**Type genus**: *Copris* Geoffroy, 1762.

**Subtribes included**:

Coprina Leach, 1815

Onychothecina *subtrib. nov*.

Pedariina *subtrib. nov*.

**Diagnosis**. The tribe is defined by the following unique combination of characters: 1) elytron with 10 striae (Figs. 9A–B, 10C); 2) absence of elytral carinae other than the epipleural one (Figs. 9A–B), or, if the elytron is carinated between stria 8 and 9 (Fig. 10C), protibia having an excavation accommodating protarsus (Fig. 10A); 3) anterior hypomeral carina present (Fig. 9C); 4) protarsus present in both sexes (Fig. 9D); 5) uncus absent in males (Fig. 9D); 6) phallobase without two tubercles proximo-dorsally (Fig. 9E); 7) FLP endophallite present (Fig. 10H).

**Description.**

Small to large beetles (ca. 5.0–30.0mm); body from moderately to very convex.

*Head*. Anterior margin of clypeus with or without 2 teeth; head often carinated, tuberculated or horned in correspondence with frontoclypeal sulcus (most Coprina), or devoid of protrusions (Pedariina, male Onychothecina and some Coprina), or with a clypeal protrusion (female Onychothecina). Antennae with 9 antennomeres.

*Pronotum*. Granulated or punctured, with or without conspicuous protrusions or carinae; mediolateral regions with a fovea; lateral pronotal carina either entire or variously effaced.

*Elytra and wings*. Elytron with 10 visible striae; not carinated except for epipleural carinae, except in Onychothecina which has a carina between striae 8 and 9. Mesoscutellum concealed. Metathoracic wings present in most species.

*Legs*. Protibia with 3 or 4 teeth; protibial distal end either obliquely or transversely truncated; male uncus absent; protarsus present in both sexes. Meso- and metatibiae strongly expanded proximo-distally; dorsal metatibial surface with (Coprina) or without (Onychothecina, Pedariina) transverse carina. Tarsal claws not toothed.

*Ventral body surface*. Anterior hypomeral carina present, stretching towards lateral margin of pronotum; posterior longitudinal hypomeral carina either present or absent, or replaced by a groove or depression.

*Tergite VIII*. Wider than long, evenly convex.

*Genitalia*. Parameres either symmetrical (Coprina) or asymmetrical (Onychothecina, Pedariina), elongated. Endophallites: A, SA, FLP, SRP present; LC present or absent.

**Remarks**. The generic composition of Coprini was long debated, with various similar-looking horned tunneller genera being assigned to it by different authors (see Montreuil (1998)). However, molecular and morphological phylogenies have widely shown the polyphyly of the tribe as historically conceived (Tarasov and Génier, 2015; Tarasov and Dimitrov, 2016; Gunter et al., 2016, 2026; Lopes et al., 2024). More recently, Tarasov and Dimitrov (2016) gave a more restrictive definition of Coprini to include only the genera closely related to *Copris*, leaving in *incertae sedis* taxa that are here accommodated in the new tribes Heliocoprini, Macroderini, Catharsiini, and some that were recently placed in the Mentophilini (Gunter et al., 2026).

Tarasov and Dimitrov (2016) and Tarasov (2017) also found a close relationship between Coprini and the genus *Pedaria*, until that moment considered a member of the Ateuchini (Vaz-de Mello, 2007). More recently, Lopes et al. (2024) found that a poorly known genus, *Onychothecus* Boucomont, 1912, is also closely related to Coprini *sensu* Tarasov and Dimitrov (2016), and accommodated it within the tribe after expanding its morphological concept. These findings are confirmed by our analyses (see also Gunter et al. (2026)), which found full support for the clade *Onychothecus*+(*Pedaria*+Coprini) (Fig. 2). Based on these results, we further expand the concept of Coprini by incorporating the genus *Pedaria*.

Morphologically, the group is relatively heterogeneous. *Onychothecus* (Fig. 10) shows some derived features (inverted sexual dimorphism, modified last tarsus and protibia), while at the same time sharing the general body shape of *Copris* and allies. *Pedaria* (Fig. 10) may seem even more morphologically derived, having a small, elongate, heavily punctate body, absence of head protrusions and pronotal horns, and strikingly asymmetrical male genitalia. Its habitus closely resembles that of the totally unrelated *Pedaridium* Harold, 1868 (Ateuchini Scatimina) and *Demarziella* Balthasar, 1961 (Mentophilini), with which it also shares the presence of a trochanterofemoral pit (Fig. 10F) (Gunter et al., 2026; Vaz-De-Mello, 2008). Lastly, the general habitus of *Copris* and closely related taxa (Fig. 9) is superficially similar to that of the unrelated genera *Ontherus* (*incertae sedis*), *Coptodactyla* (Mentophilini), *Catharsius* (Catharsiini) and *Xinidium* (Macroderini), all tunnellers characterised by strong sexual dimorphism with heavily armed males and elongated, usually medium- to large-sized body.

For these reasons, while we unite *Copris*, *Onychothecus* and *Pedaria* within Coprini, we define three subtribes in order to preserve the notion of their morphological disparity.

### Coprina Leach, 1815

(Figs. 9F–I)

**Type genus**: *Copris* Geoffroy, 1762.

**Genera included:**

*Copris* Geoffroy, 1762 (232 valid species; Afrotropical, Nearctic, Neotropical, Indomalayan, Palaearctic);

*Litocopris* Waterhouse, 1891 (4 valid species; Afrotropical);

*Microcopris* Waterhouse, 1891 (11 valid species; Indomalayan, Palaearctic);

*Paracopris* Balthasar, 1939 (26 valid species; Afrotropical, Indomalayan, Palaearctic);

*Pseudocopris* Ferreira, 1960 (2 valid species; Afrotropical);

*Pseudopedaria* Felsche, 1904 (6 valid species; Afrotropical);

*Sinocopris* Ochi, Kon & Bai, 2009 (24 valid species; Indomalayan, Palaearctic).

**Diagnosis**. The subtribe is defined by the following combination of characters, separating it from Onychothecina and Pedariina: 1) wing with posterior sclerite of RP1 present (Fig. 9F); 2) posterior longitudinal hypomeral carina usually present (Fig. 9G).

**Description.**

Medium to large sized beetles (ca. 8.5–30.0mm); body from moderately to very convex.

*Head*. Anterior margin of clypeus mostly with 2 teeth; head mostly carinated, tuberculated or horned in correspondence with frontoclypeal sulcus or on the frons (all genera), or devoid of protrusions (genera *Litocopris*, most *Pseudopedaria* and *Microcopris*).

*Pronotum*. Granulated or punctured; male and, to a lesser extent, female pronotum often horned or carinated on disc (most species of *Copris* and *Sinocopris*), or devoid of protrusions (remaining genera); mediolateral regions with a fovea; lateral pronotal carina entire.

*Elytra and wings*. Elytron with 10 visible striae, striae 9 and 10 closely adjacent and sometimes partially fused anteriorly; not carinated except for epipleural carina. Metathoracic wings present in most species.

*Legs*. Anterior leg without trochanterofemoral pit; protibia with 3 or 4 teeth; protibial distal end either transversely (*Litocopris*) or obliquely (remaining genera) truncated. Dorsal metatibial surface with transverse carina.

*Ventral body surface*. Posterior longitudinal hypomeral carina usually present.

*Genitalia*. Parameres symmetrical, elongated, rarely with spines; SRP not ring-shaped.

**Distribution and biology**. The subtribe is distributed in the Afrotropical, Nearctic, Neotropical, Indomalayan and Palaearctic realms. It comprises medium to large sized tunnellers feeding almost exclusively on dung (Davis et al., 2008).

**Remarks**. The diagnosis of Coprina corresponds to the concept of Coprini by Tarasov and Dimitrov (2016). It is a rather morphologically homogeneous clade on whose monophyly there is little doubt (present analyses; Tarasov and Dimitrov, 2016; Tarasov and Génier, 2015; Monaghan et al., 2007). The genera *Sinocopris* and *Pseudocopris* were never included in molecular analyses. However, morphology strongly supports their placement within the subtribe (Marchisio and Zunino, 2012).

Several genera of Coprina (*Litocopris*, *Microcopris*, *Paracopris* and *Sinocopris*) were historically considered subgenera of *Copris* or even synonyms of it (Marchisio and Zunino, 2012; Balthasar, 1963; Ferreira, 1961; Zidek, 2020). More recently, they were given full genus rank by some authors (Philips and Moretto, 2025; Zidek, 2020). For practicality, we chose to adopt the latter solution here. However, several sources of evidence suggest that *Copris* as currently defined is paraphyletic with respect to most genera. *Microcopris* and *Litocopris* were found to be nested within it in multiple molecular studies (Tarasov and Dimitrov, 2016; Monaghan et al., 2007), including the present one, and the same was suggested for *Litocopris*, *Pseudocopris* and *Pseudopedaria* based on morphology (Marchisio and Zunino, 2012). On the other hand, Marchisio and Zunino (2012) hinted at the possibility that the group of *Copris* species close to *C. bootes* could represent a distinct clade, perhaps more closely related to the Neotropical genus *Ontherus* (*incertae sedis*). This hypothesis is not supported by our reconstruction, which found a species of the same group (*C. nevinsoni* Waterhouse, 1891) deeply nested within *Copris*. Overall, a comprehensive phylogenetic survey of the subtribe is needed to improve its current generic classification.

One important feature defining Coprini, including Coprina, is the presence of 10 elytral striae. In the literature, this number was most often mistaken, as striae 9 and 10 were overlooked (e.g. Ferreira, 1961; Marchisio and Zunino, 2012; Janssens, 1946). This is due to the fact that striae 9 and 10 in Coprina are usually closely adjacent to the epipleural carina and sometimes tightly appressed or fused to one another on the anterior half, which makes their observation (particularly of stria 10) somewhat difficult. However, a careful examination after cleaning elytra in KOH is sufficient to reveal them, showing they are present in all species we examined.

### Onychothecina *subtrib. nov*

(Figs. 10A–D)

**Type genus**: *Onychothecus* Boucomont, 1912.

**Genus included:**

*Onychothecus* Boucomont, 1912 (4 species; Indomalayan, Palaearctic).

**Diagnosis**. The subtribe is defined by the following combination of characters, separating it from Coprina and Pedariina: 1) elytron carinated between stria 8 and 9 (Fig. 10C); 2) last tarsomere expanded and concealing tarsal claws (Fig. 10B); 3) protibia with a distal excavation accommodating protarsus (Fig. 10A).

**Description.**

Large sized beetles (18.0–26.8mm); body moderately convex.

*Head*. Anterior margin of clypeus with 2 teeth; female clypeus with medial horn-like protrusion; frontoclypeal sulcus absent.

*Pronotum*. Surface punctured; more or less gibbose anteriorly in the female; mediolateral regions with a fovea; lateral pronotal carina entire.

*Elytra and wings*. Elytron with 10 visible striae; with a carina between striae 8 and 9. Metathoracic wings present.

*Legs*. Anterior leg without trochanterofemoral pit; protibia with 3 teeth; protibial distal end transversely truncated. Dorsal metatibial surface without transverse carina.

*Ventral body surface*. Posterior longitudinal hypomeral carina absent.

*Genitalia*. Parameres asymmetrical (at least in *O. tridentigeris*); SRP not ring-shaped.

**Distribution and biology**. *Onychothecus* is found in the Indomalayan and Palaearctic (China) realms (Lopes et al., 2024). Nothing is known about its biology.

**Remarks**. *Onychothecus* was originally placed in the Scatonomini (now a synonym of Deltochilini) (Boucomont, 1912) and then in the Pinotini (now synonym of Dichotomiini) (Balthasar, 1963). The latter solution was also adopted by Vaz-de Mello (2007), who proposed a new monotypic ateuchine subtribe, Onychothecina, for this genus—the name is unpublished and therefore nomenclaturally unavailable. Based on morphological evidence, *Onychothecus* was subsequently declared *incertae sedis* because it does not fit any existing tribe (Tarasov and Dimitrov, 2016), and was most recently transferred into Coprini based on the first molecular data available for the genus (Lopes et al., 2024). UCE-based analyses fully support the placement of *Onychothecus* within Coprini, in which it represents the earliest-branching lineage (present analyses; Gunter et al., 2026; Lopes et al., 2024).

### Pedariina subtrib. nov

(Figs. 10E–I, 3K)

**Type genus**: *Pedaria* Laporte, 1832.

**Genera included:**

*Pedaria* Laporte, 1832 (68 valid species; Afrotropical).

**Diagnosis**. The subtribe is defined by the following combination of characters, separating it from Coprina and Onychothecina: 1) proleg with trochanterofemoral pit (Fig. 10F); 2) putative posterior longitudinal carina reduced and flanked by a lateral transverse depression (Fig. 10E); 3) parameres strongly asymmetrical (Fig. 10G); 4) SRP endophallite ring-shaped (Fig. 10H).

**Description.**

Small- to medium-sized beetles (ca. 5.0–15.0mm); body elongated, moderately convex.

*Head*. Anterior margin of clypeus usually with 2 teeth; head devoid of protrusions, frontoclypeal sulcus absent.

*Pronotum*. Densely covered with ocellate punctures; evenly convex, produced posteromedially into a spiniform process or produced anteromedially into a lobular process in some species; mediolateral regions with a small fovea; lateral pronotal carina variously developed, often restricted to the posterolateral part of pronotum.

*Elytra and wings*. Elytron with 10 striae, stria 9 not reaching elytral base; not carinated except for epipleural carina. Metathoracic wings present.

*Legs*. Anterior leg with trochanterofemoral pit; protibia with 3 teeth; protibial distal end transversely truncated; protibial spur expanded and bifid in males.

*Ventral body surface*. Anterior hypomeral region strongly depressed; putative posterior longitudinal hypomeral carina present, reduced, hypomeron with a groove or depression between the posterior hypomeral carina and the pronotal carina.

*Genitalia*. Parameres strongly asymmetrical. Endophallites: SRP ring-shaped; distal part of endophallus with raspulae at least in some species.

**Distribution and biology**. The tribe is restricted to the Afrotropical realm. Its species are known or suspected kleptoparasites of large tunnellers (Davis et al., 2008).

**Remarks**. The genus *Pedaria* was for a long time considered a member of the Pinotini (now Ateuchini) (Montreuil, 1998; Vaz-de Mello, 2007). Vaz-de Mello (2007) suggested its placement within the Demarziellini (now synonym of Mentophilini; Gunter et al., 2026), and later designated it as *incertae sedis* within the Ateuchini, as it did not fit any described ateuchine subtribe (Vaz-De-Mello, 2008). Tarasov and Dimitrov (2016) later moved it to Scarabaeinae *incertae sedis*.

Morphologically, *Pedaria* strongly resembles the genus *Demarziella* (Mentophilini), which led various authors to suggest relatedness between these taxa (Vaz-de Mello, 2007; Montreuil, 1998; Tarasov and Génier, 2015). A turning point was represented by recent molecular phylogenies, including the present one, which consistently recovered the genus as sister to *Copris* and related genera, showing that morphological similarity between *Pedaria* and Mentophilini is largely due to convergence (Tarasov and Dimitrov, 2016; Gunter et al., 2026; Lopes et al., 2024; Tarasov, 2017).

### Coptorhinini Scholtz, Davis & De Klerk, 2025

(Fig. 3A)

**Type genus**: *Coptorhina* Hope, 1833.

**Genera included:**

*Coptorhina* Hope, 1833 (7 species; Afrotropical);

*Delopleurus* Erichson, 1848 (10 valid species; Afrotropical, Indomalayan);

*Frankenbergerius* Balthasar, 1938 (7 valid species; Afrotropical);

*Sarophorus* Erichson, 1847 (12 valid species; Afrotropical).

**Remarks**. The tribe was recently described by Scholtz et al. (2025) based on solid molecular and morphological evidence, so we avoid discussing it again here. The groups falls within the Coptorhinomorpha clade, where it seems to be most closely related to Paraphytini (Fig. 1; see Discussion).

### Dwesasilvasedini *trib. nov*

(Fig. 11)

**Figure 11.**
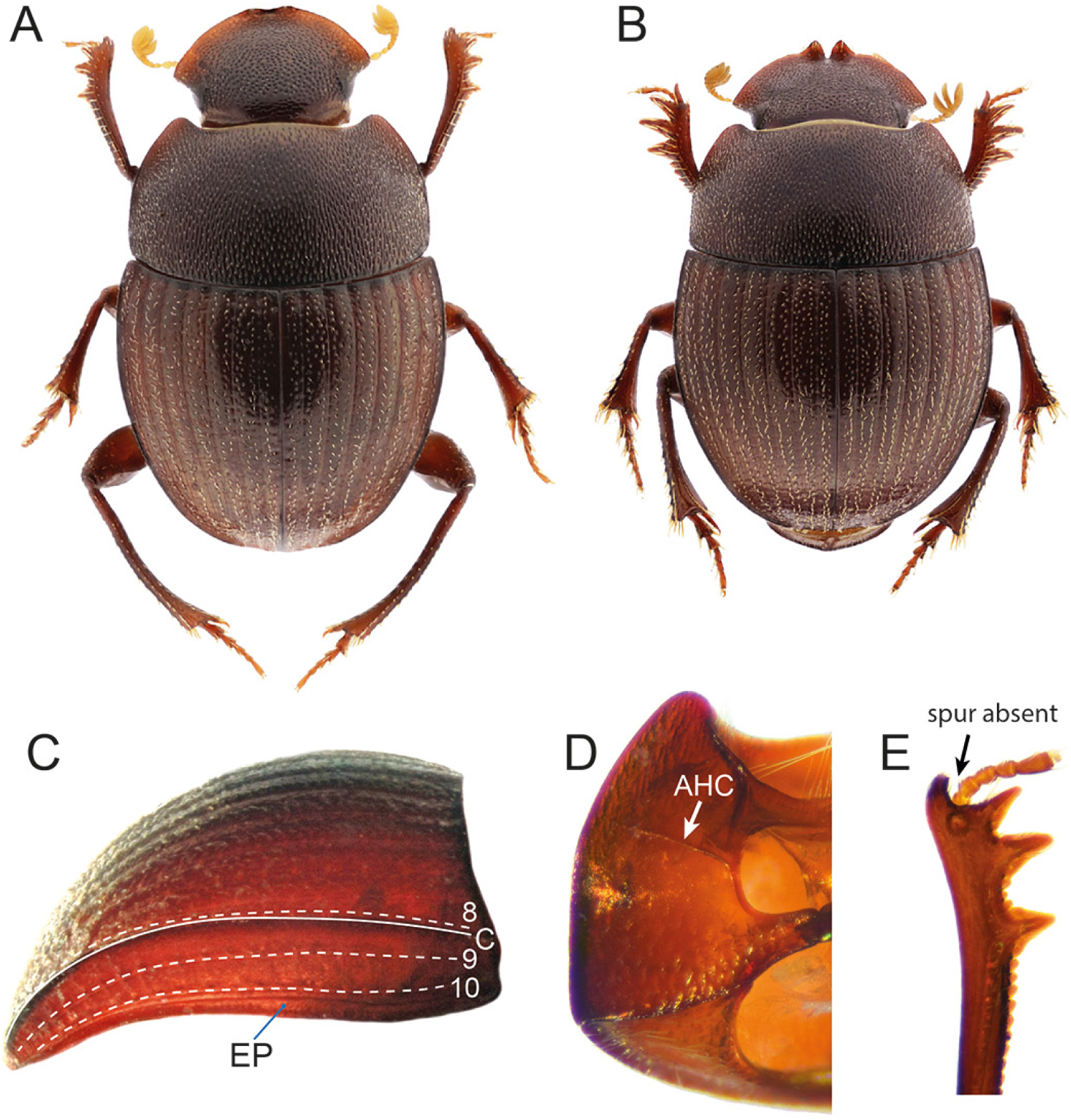
Diagnostic morphological characters of Dwesasilvasedini *trib. nov.* *Dwesasilvasedis medinae*: (A) male habitus (picture by C. Deschodt); (B) female habitus (picture by C. Deschodt); (C) elytron showing lateral striae (8–10) and carina (C; epipleural carina not marked); (D) hypomeron showing anterior hypomeral carina; (E) male protibia showing absence of spur.

**Type genus**: *Dwesasilvasedis* Deschodt & Scholtz, 2008.

**Genus included:**

*Dwesasilvasedis* Deschodt & Scholtz, 2008 (1 valid species; Afrotropical).

**Diagnosis**. The tribe is defined by the following unique combination of characters: 1) elytron with 10 visible striae, striae 9 and 10 placed on pseudoepipleuron (Fig. 11C); 2) presence of a single elytral carina lateral to stria 8 (Fig. 11C); 3) protibial spur absent in males (Fig. 11E); 4) male and female protarsus present (Fig. 11E); 5) anterior hypomeral carina stretching toward lateral margin of hypomeron (Fig. 11D).

**Description.**

Small sized beetles (4.2–4.8 mm); body oval, moderately convex, covered with short pale setae.

*Head*. Clypeus with two teeth, teeth more developed in females; head margin evenly rounded overall; frontoclypeal sulcus absent. Antennae with 9 antennomeres.

*Pronotum*. Moderately convex; covered with dense oval punctures; lateral pronotal carina entire.

*Elytra and wings*. Elytron with 10 visible striae; elytron carinate lateral to stria 8, carina with one ridge; elytron slanted inwards laterally to carina; striae 9 and 10 placed on pseudoepipleuron. Mesoscutellum concealed. Metathoracic wings absent.

*Legs*. Protibia with 3 teeth, distal margin oblique; male protibia with uncus; protibial spur absent in males, present in females; protarsus present in both sexes. Meso- and metatibiae strongly but evenly expanded proximo-distally. Tarsal claws not toothed.

*Ventral body surface*. Anterior hypomeral carina stretching towards lateral margin of hypomeron; posterior longitudinal hypomeral carina absent. Mesometaventral sulcus angular medially; metaventrite punctured, punctures smaller anteriorly.

*Tergite VIII*. Triangular, punctured.

*Genitalia*. Parameres asymmetrical, left paramere longer and with posterior apex expanded. Endophallites: FLP, SA and A sclerites present; LC absent; SRP with ring-shaped part.

**Distribution and biology**. The only known species of Dwesasilvasedini, *Dwesasilvasedis medinae* Deschodt & Scholtz, 2008, is restricted to Dwesa coastal forest in the Eastern Cape province of South Africa, where it was collected by sifting forest litter (Davis et al., 2020; Deschodt and Scholtz, 2008).

**Remarks**. *Dwesasilvasedis* was until recently considered a member of the Canthonini (Deschodt and Scholtz, 2008), and later treated as *incertae sedis* (Tarasov and Dimitrov, 2016). Our phylogeny recovered it within the Onthophagomorpha clade next to Macroderini and Onitini (Fig. 2), in substantial agreement with previous phylogenies (Mlambo et al., 2014, 2015; Tarasov and Dimitrov, 2016; Sole and Scholtz, 2010). The exact relationships between the three tribes, however, differ between different works.

Morphologically, Dwesasilvasedini are well characterised by the configuration of elytral striation, the presence of protarsi in both sexes, and the absence of protibial spur in males. The latter character, rare in Scarabaeinae, is shared with the closely allied Onitini (see in Onitini Remarks). This contributed to recovering Dwesasilvasedini and Onitini as sisters in the morphological analysis by Tarasov and Génier (2015). Consistently with molecules, the presence of 10 elytral striae, of which the 9th does not reach the anterior elytral margin, and the presence of a carina lateral to stria 8, also support a close relationship with Macroderini and Onitini (see also Discussion). Overall, multiple sources of evidence strongly justify the establishment of Dwesasilvasedini *trib. nov.* as a separate tribe.

### Elassocanthonini Davis, Deschodt & Scholtz, 2025

(Fig. 3B)

**Type genus**: *Elassocanthon* Kolbe, 1908.

**Genera included:**

*Ausmontins* Deschodt & Davis, 2018 (1 valid species; Afrotropical);

*Dicranocara* Frolov & Scholtz, 2003 (4 valid species; Afrotropical);

*Drogo* Deschodt, Davis & Scholtz, 2016 (1 valid species; Afrotropical);

*Namakwanus* Scholtz & Howden, 1987 (4 valid species; Afrotropical);

*Namaphilus* Deschodt & Davis, 2017 (5 valid species; Afrotropical);

*Versicorpus* Deschodt, Davis & Scholtz, 2011 (3 valid species; Afrotropical).

**Revised diagnosis**. The tribe is defined by the following unique combination of characters: 1) accessory endophallites comprising only A and BSc endophallites; 2) basal margin of spiculum gastrale with narrow process apically; 3) epipleuron very wide; 4) hind wings absent; 5) glossal flap bending inward, its tip covered with big blunt spurs.

**Remarks**. Elassocanthonini were previously known as Byrrhidiini Davis, Deschodt & Scholtz, 2019 (Davis et al., 2019), but later renamed due to homonymy of the type genus, *Byrrhidium* Harold, 1869, currently *Elassocanthon* Kolbe, 1908 (Davis et al., 2025). The monophyly of the tribe is well supported both morphologically and molecularly (Tarasov and Génier, 2015; Tarasov and Dimitrov, 2016; present analyses). The revised morphological diagnosis provided above enriches the original one with additional characters scored from the group synapomorphies found by Tarasov and Génier (2015). The tribe falls within the Coptorhinomorpha clade and is most closely related to Endroedyolini and Haroldiini (Fig. 1; see Discussion).

### Endroedyolini Davis, Deschodt & Scholtz, 2019 *sensu novo*

(Fig. 12)

**Figure 12.**
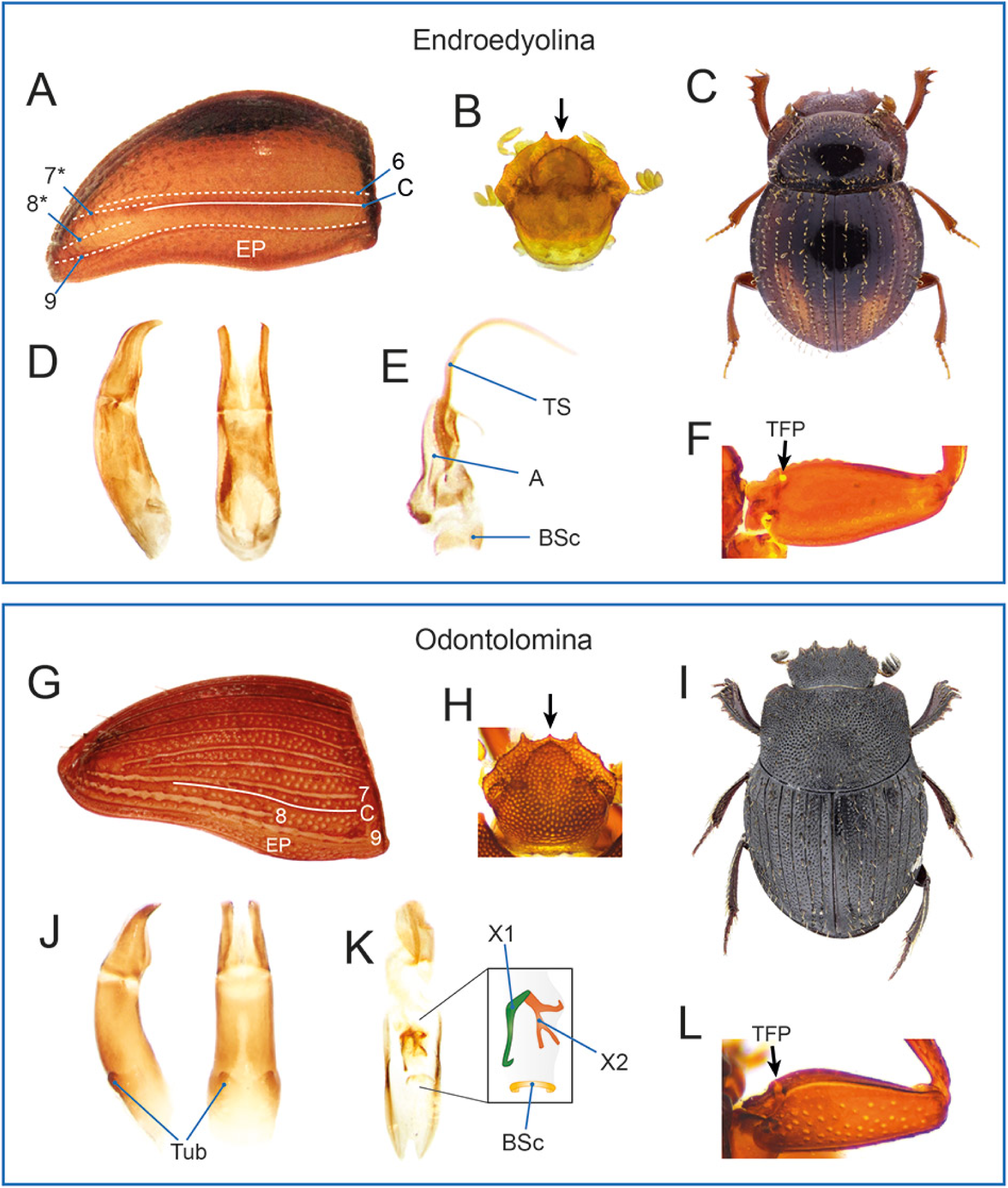
Diagnostic morphological characters of Endroedyolini *sensu novo*. Endroedoyolini Endroedyolina: (A) *Endroedyolus paradoxus*, elytron showing lateral striae (6–9; traces of striae 7 and 8 are marked with an asterisk) and carina (C; epipleural carina not marked); (B) *E. paradoxus*, head, arrow pointing at absence of medial clypeal tooth; (C) *E. paradoxus*, habitus (picture by C. Deschodt); (D) *E. paradoxus*, aedeagus in lateral (left) and dorsal (right) views; (E) *Nebulasilvius insularis*, endophallites; (F) *E. paradoxus*, profemur showing trochanterofemoral pit. Endroedyolini Odontolomina: (G) *Odontoloma* sp., elytron showing lateral striae (7–9) and carina (C; epipleural carina not marked); (H) *Odontoloma* sp., head, arrow pointing at presence of medial clypeal tooth; (I) *O. louwi*, habitus (picture by F. Génier); (J) *Odontoloma* sp., aedeagus in lateral (left) and dorsal (right) views, showing the presence of basal tubercles; (K) *Odontoloma* sp., endophallites; (L) *Odontoloma* sp., profemur showing trochanterofemoral pit.

**Type genus**: *Endroedyolus* Scholtz & Howden, 1987.

**Subtribes included:**

Endroedyolina Davis, Deschodt & Scholtz, 2019 *stat. nov*.

Odontolomina Davis, Deschodt & Scholtz, 2019 *stat. nov*.

**Diagnosis**. The tribe is defined by the following unique combination of characters: 1) parameres simple, tiny, apically acute (Figs. 12D, 12J); 2) BSc endophallite present, semicircularly shaped (Figs. 12E, 12K); 3) elytron with 9 striae, striae 7 and 8 sometimes indistinct (Figs. 12A, 12G); 4) elytron with elytral carina lateral to stria 7 (Figs. 12A, 12G); 5) proleg with trochanterofemoral pit (Figs. 12F, 12L).

**Description.**

Very small to small sized beetles (1.5–4.8 mm); moderately flattened to very convex.

*Head*. Clypeus with 2–4 bilateral teeth and, in Odontolomina, a small medial tooth; head margin at genoclypeal junctions with or without tooth; head surface without carinae or protrusions; frontoclypeal sulcus absent. Antennae with 9 antennomeres.

*Pronotum*. Variously punctuated, without protrusions; lateral pronotal carina entire.

*Elytra and wings*. Elytron with 9 visible striae, sometimes striae 7 and 8 reduced and inconspicuous; elytron carinated laterally to stria 7. Mesoscutellum concealed. Metathoracic wings present or absent.

*Legs*. Protibia with 3 teeth, distal margin more or less clearly transversely truncated; uncus not developed; proleg with trochantofemoral pit. Meso- and metatibiae evenly expanded proximo-distally. Tarsal claws not toothed.

*Ventral body surface*. Anterior hypomeral carina present, stretching to lateral angles of pronotum; posterior longitudinal hypomeral carina absent.

*Tergite VIII*. Foveate or not.

*Genitalia*. Parameres symmetrical, tiny, apically acute, with or without two tubercles proximo-dorsally. Endophallites (see also Odontolomina Remarks): accessory endophallites comprising only axial and BSc endophallites; axial endophallite with central lobe (TS) thin and long; BSc endophallite semicircularly shaped.

**Remarks**. Relatedness between the former Endroedyolini and Odontolomini was recovered by multiple earlier phylogenies (Mlambo et al., 2015; Sole and Scholtz, 2010; Tarasov and Génier, 2015; Tarasov and Dimitrov, 2016), although relationships with closely related tribes (especially Elassocanthonini and Haroldiini; see Discussion) differed between studies. Our phylogenetic reconstructions fully support a sister relationship between *Odontoloma* and *Endroedyolus*, consistently with the morphological analysis by Tarasov and Génier (2015). Morphologically, Endroedyolini and Odontolomini are quite similar and share many important synapomorphies in elytral striation, male genitalia, legs, and other traits (see Diagnosis). Additionally, their divergence is considerably shallower than that of most other sister tribes (Fig. 1).

On the other hand, the two groups maintain some degree of morphological differentiation. Main differences are in the head, elytra, endophallites and body shape. The head bears a tiny medial clypeal tooth and a tooth in correspondence with genoclypeal sutures in Odontolomini that are absent or inconspicuous in Endroedyolini. The elytra are sometimes subcarinate in Odontolomini with inconspicuous setae on the interstriae whereas they have clear setae and are evenly convex in Endroedyolini. Amongst the endophallites, the axial sclerites are differently shaped between the two tribes whereas the body is strongly convex in Endroedyolini but somewhat more flattened in Odontolomini (Fig. 12). Moreover, the two clades are relatively well differentiated ecologically: the Endroedyolini are restricted to relict forests of South Africa, while the Odontolomini are widespread in a variety of habitats in the Afrotropical realm.

Taken together, these results highlight that the most appropriate solution is to unite Odontolomini and Endroedyolini in order to avoid dispersion of morphologically homogeneous clades into separate small tribes. At the same time, since they retain some degree of morphological and ecological differentiation, we prefer to keep Odontolomina *stat. nov*. as a valid subtribe within Endroedyolini *sensu novo*.

### Endroedyolina Davis, Deschodt & Scholtz, 2019 *stat. nov*

(Fig. 12A–F)

**Genera included:**

*Aliuscanthoniola* Deschodt & Scholtz, 2008 (1 valid species; Afrotropical);

*Endroedyolus* Scholtz & Howden, 1987 (1 valid species; Afrotropical);

*Nebulasilvius* Deschodt & Scholtz, 2008 (2 valid species; Afrotropical);

*Outenikwanus* Scholtz & Howden, 1987 (1 valid species; Afrotropical);

*Parvuhowdenius* Deschodt & Scholtz, 2008 (1 valid species; Afrotropical);

*Peckolus* Scholtz & Howden, 1987 (3 valid species; Afrotropical);

*Silvaphilus* Roets & Oberlander, 2010 (2 valid species; Afrotropical);

*Upsa* Deschodt, Sole & Scholtz, 2020 (1 valid species; Afrotropical).

**Diagnosis**. The subtribe is defined by the following combination of characters, separating it from Odontolomina: 1) body very convex, covered with long, conspicuous setae (Fig. 12C); 2) metathoracic wings absent; 3) clypeus without small medial tooth, bearing 2 to 4 bilateral teeth (Fig. 12B); 4) phallobase without a pair of tubercles proximo-dorsally (Fig. 12D); 5) subaxial endophallite absent, central lobe of axial endophallite thin and long, surrounding lobe notched on right side (Fig. 12E).

**Description**. See details in Davis et al. (2019).

**Distribution and biology**. The genera of Endroedyolina are restricted to forest fragments in southern and southeastern South Africa, where they occur in leaf litter (Davis et al., 2020). An exception is one species of *Silvaphilus*, which has been collected in fynbos riparian vegetation in the Cederberg Mountains of South Africa (Daniel et al., 2022). Beyond these records, little is known about their nesting behavior, feeding preferences, or other aspects of natural history, except that *Silvaphilus* species have been collected in herbivore- and pig dung-baited traps (Daniel et al., 2022; Roets and Oberlander, 2010).

**Remarks**. The Endroedyolina comprise mono- or oligospecific genera of relict forest-specialised beetles. They are morphologically homogeneous, differing mainly in features of elytral striation, meso- and metaventrites, and hypomera. Notably, elytral striae 7 and 8 can be vestigial and reduced to the posterior region of elytron (*e.g.*, *Endroedyolus*, *Nebulasilvius*), or normally developed also anteriorly (*e.g.*, *Silvaphilus*). Contrarily to the Odontolomina, all Endroedyolina are wingless. Although the majority of species has only two clypeal teeth, some show two additional teeth (*Upsa*), or a small hint of a medial tooth (*Silvaphilus*), recalling the Odontolomina which usually have five teeth on the head margin.

### Odontolomina Davis, Deschodt & Scholtz, 2019 *stat. nov*

(Fig. 12G–L)

**Type genus**: *Odontoloma* Boheman, 1857.

**Genus included:**

*Odontoloma* Boheman, 1857 (20 valid species; Afrotropical).

**Diagnosis**. The subtribe is defined by the following combination of characters, separating it from Endroedyolina: 1) body convex, covered with short setae (Fig. 12I); 2) metathoracic wings usually present; 3) clypeus with a small medial tooth and 2 bilateral teeth, plus 2 bilateral teeth next to genoclypeal junctions and occasionally 2 others on the genae (Fig. 12H); 4) phallobase with a pair of tubercles proximo-dorsally (Fig. 12J); 5) subaxial endophallite present (but see below) (Fig. 12K).

**Description**. See details in Davis et al. (2019).

**Distribution and biology**. The Odontolomina are widely distributed in the Afrotropical realm, although with a strong bias towards southern Africa, where they occupy various habitats, from forest to savanna to arid mountainous regions. Some species have been attracted to carrion, herbivore and omnivore dung (Davis et al., 2008, 2020). Little is known about other life history traits.

**Remarks**. The Odontolomina comprise a single genus *Odontoloma*, that is well characterised among other genera historically placed within the Canthonini due to its habitus reminiscent of some species of Onthophagini. *Odontoloma* species are quite homogeneous morphologically, differing mostly in integument punctuation, clypeal denticulation, and male genitalia (Howden and Scholtz, 1987). Contrarily to Endroedyolina and other tribes within Coptorhinomorpha, which lack the subaxial endophallite, at least some odontolomines have two endophallites that are putatively homologous with the axial and subaxial endophallites of other Scarabaeinae, with the putative axial sclerite being relatively different from that of other coptorhinomorphans (Figs. 12K) (Tarasov and Génier, 2015). However, a comprehensive survey of genital morphology is required to assess if these features are shared by all *Odontoloma* species.

### Epactoidini Rossini, Grebennikov, Merrien, Miraldo, Viljanen & Tarasov, 2022

**Type genus**: *Epactoides* Olsoufieff, 1947.

**Genera included:**

*Epactoides* Olsoufieff, 1947 (38 valid species; Madagascan).

*Grebennikovius* Mlambo, Scholtz & Deschodt, 2019 (4 valid species; Afrotropical).

*Ochicanthon* Vaz-de-Mello, 2003 (54 valid species; Indomalayan).

**Remarks**. The tribe was recently described by Rossini et al. (2022) based on solid molecular and morphological evidence, so we avoid discussing it again here. According to multiple reconstructions, it is closely related to Epirinini and Sisyphini (Fig. 1; see Discussion).

### Epirinini Van Lansberge, 1874

(Fig. 3G)

**Type genus**: *Epirinus* Dejean, 1833.

**Genus included:**

*Epirinus* Dejean, 1833 (34 valid species; Afrotropical).

**Remarks**. The tribe was recently redefined based on a sound combination of molecular, morphological and biogeographical evidence (Daniel et al., 2020b, 2021a; Tarasov and Dimitrov, 2016), so we avoid discussing it again here. Our reconstruction confirms previous findings, recovering Epirinini as sister to Sisyphini (Fig. 1; see Discussion). Tarasov and Dimitrov (2016) proposed to merge the two groups, but they are now accepted as separate tribes (Daniel et al., 2021a).

### Gymnopleurini Streubel, 1846

(Figs. 13)

**Figure 13.**
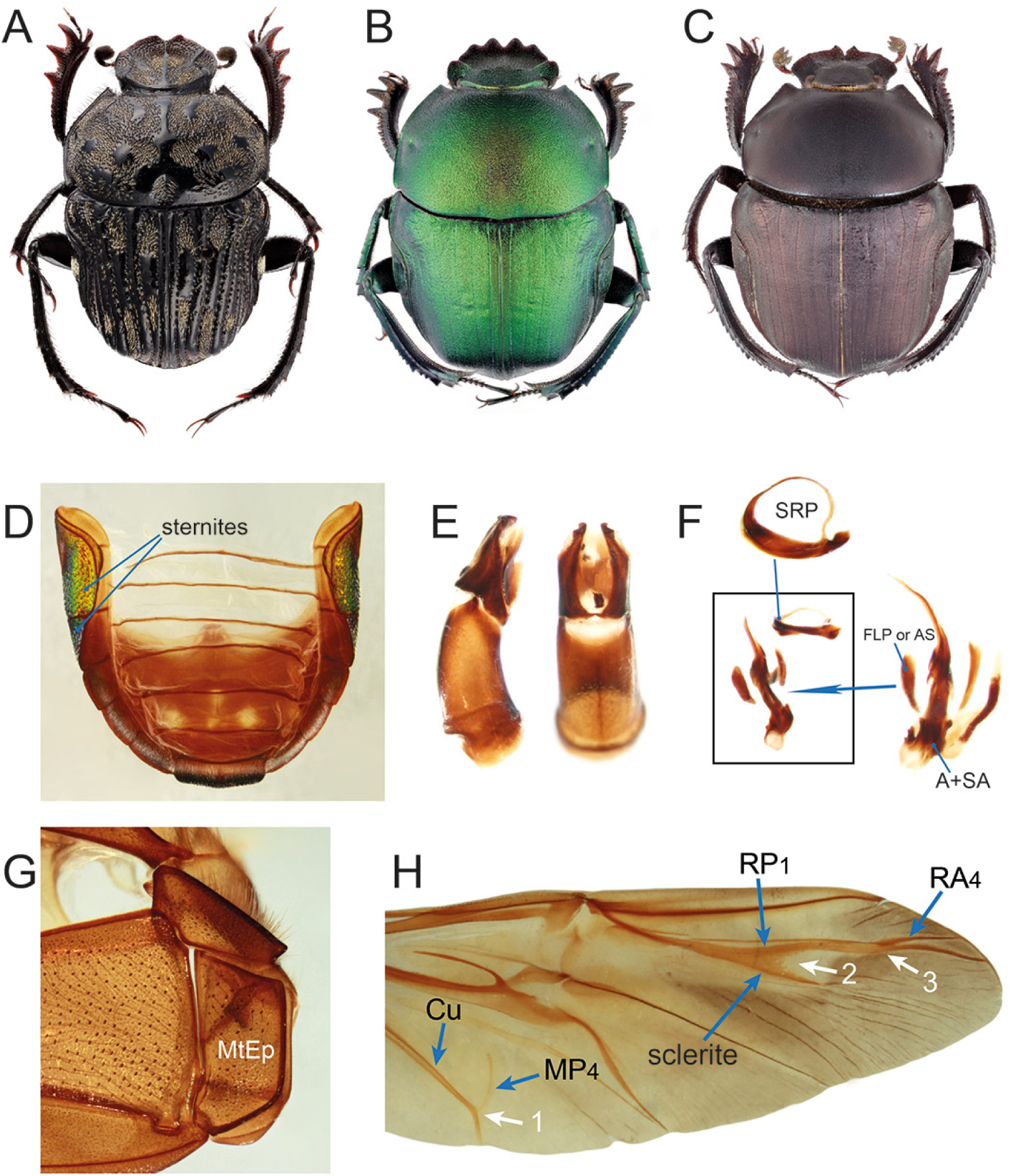
Diagnostic morphological characters of Gymnopleurini. (A) *Gymnopleurus koenigi*, habitus; (B) *Garreta wahlbergi*, habitus; (C) *Paragymnopleurus* sp., habitus; (D) *Gymnopleurus leei*, abdomen showing lateral surface of sternites 2–4 curved dorsally; (E) *Paragymnopleurus* sp., aedeagus in lateral (left) and dorsal (right) views; (F) *Paragymnopleurus* sp., endophallites; (G) *Paragymnopleurus* sp., lateral region of metathorax showing subrectangular, posteriorly rounded metanepisternum; (H) *G. leei*, detail of metathoracic wing showing MP4 vein curved and fused with Cu (arrow 1), RP1 vein and its posterior sclerite fused proximally but diverging distally (arrow 2) and RA4 vein gradually thinner medially and fusing with RP1 (arrow 3).

**Type genus**: *Gymnopleurus* Illiger, 1803.

**Genera included:**

*Allogymnopleurus* Janssens, 1940 (18 valid species; Afrotropical, Palaearctic, Indomalayan);

*Garreta* Janssens, 1940 (28 valid species; Afrotropical, Palaearctic, Indomalayan);

*Gymnopleurus* Illiger, 1803 (62 valid species; Afrotropical, Palaearctic, Indomalayan);

*Paragymnopleurus* Shipp, 1897 (13 valid species; Palaearctic, Indomalayan).

**Diagnosis**. The tribe is defined by the following unique combination of characters: 1) axial endophallite associated with multiple surrounding sclerites (Fig. 13F); 2) lateral elytral margin emarginate on anterior half; 3) wing with MP4 curved and fused with Cu (Fig. 13H); 4) wing with RA4 gradually thinner medially where it fuses with RP1 (Fig. 13H); 5) wing with RP1 and its posterior sclerite fused proximally but bifurcate distally, distinctly diverging from each other (Fig. 13H); 6) metanepisternum more or less rectangular with apical part rounded, in lateral view (Fig. 13G); 7) lateral surface of sternites 2–4 curved dorsally (Fig. 13D).

**Description.**

Small to large sized beetles (6.0–22.0 mm); moderately flattened.

*Head*. Clypeus with 2–6 teeth; head surface mostly without carinae or protrusions, sometimes with one or two longitudinal or oblique carinae; frontoclypeal sulcus absent. Antennae with 9 antennomeres.

*Pronotum*. Variously punctuated, without protrusions; posterior margin rarely with a medial pair of depressions; lateral pronotal carina entire.

*Elytra and wings*. Elytron with 9 visible striae; absence of elytral carinae; anterior half of lateral elytral margin strongly emarginate. Mesoscutellum concealed. Metathoracic wings present; RP1 and its posterior sclerite fused proximally but bifurcate distally, distinctly diverging from each other; MP4 curved and fused with Cu; RA4 gradually thinner medially where it fuses with RP1.

*Legs*. Protibia with 3 teeth; uncus absent, distoventral margin of male protibia often protruding; protibial spur sharply tapering in females, often modified in males. Meso- and metatibiae feebly expanded proximo-distally, elongated. Tarsal claws not toothed.

*Ventral body surface*. Anterior hypomeral carina present, stretching to anterior angles of pronotum; posterior longitudinal hypomeral carina absent. Metanepisternum more or less rectangular with apical part rounded. Lateral surface of abdominal sternites 2—4 curved dorsally, visible from above due to the sinuous emargination of the elytra; metepimeron either clearly visible from above in the lateral elytral emargination (*Allogymnopleurus*, *Garreta*, *Paragymnopleurus*) or not (*Gymnopleurus*).

*Tergite VIII*. Evenly flattened, triangular.

*Genitalia*. Parameres symmetrical. Endophallites: A and SA endophallites present; presence of additional sclerites around A+AS, one of which may be homologous to FLP (see Tarasov and Génier (2015)); LC absent; SRP ring-shaped.

**Distribution and biology**. The tribe comprises coprophagous rollers widely distributed in Afro-Eurasia, with diversity peaking in the Afrotropical realm. Gymnopleurini inhabit various ecosystems, from desert to rainforest (Davis et al., 2008).

**Remarks**. Due to its unique morphological features and easy diagnosis, the composition of Gymnopleurini has been very stable historically. The tribe was revised by Janssens (1940), who established the genera *Garreta* and *Allogymnopleurus* for two groups of species previously part of *Gymnopleurus*. Later, Kabakov (2006) erected the subgenus *Gymnopleurus* (*Metagymnopleurus*) Kabakov, 2006, recently synonimised with the nominotypical one (Moretto, 2025). Among Scarabaeinae, Gymnopleurini are very characteristic due to the surface of sternites 2–4, which is extended dorsally and exposed through the lateral emargination of the elytron. Here, we provided a more thorough definition of the tribe by adding additional characters, including unique synapomorphies in wing venation and male genitalia. Monophyly of the tribe is fully supported both morphologically (Tarasov and Génier, 2015) and molecularly (present analyses; Gunter et al., 2026, 2016; Mlambo et al., 2015; Monaghan et al., 2007; Tarasov and Dimitrov, 2016). Its relationships with other tribes, however, are discordant between phylogenies, so that no close relatives of Gymnopleurini can be reliably identified.

### Haroldiini *trib. nov*

(Fig. 14)

**Figure 14.**
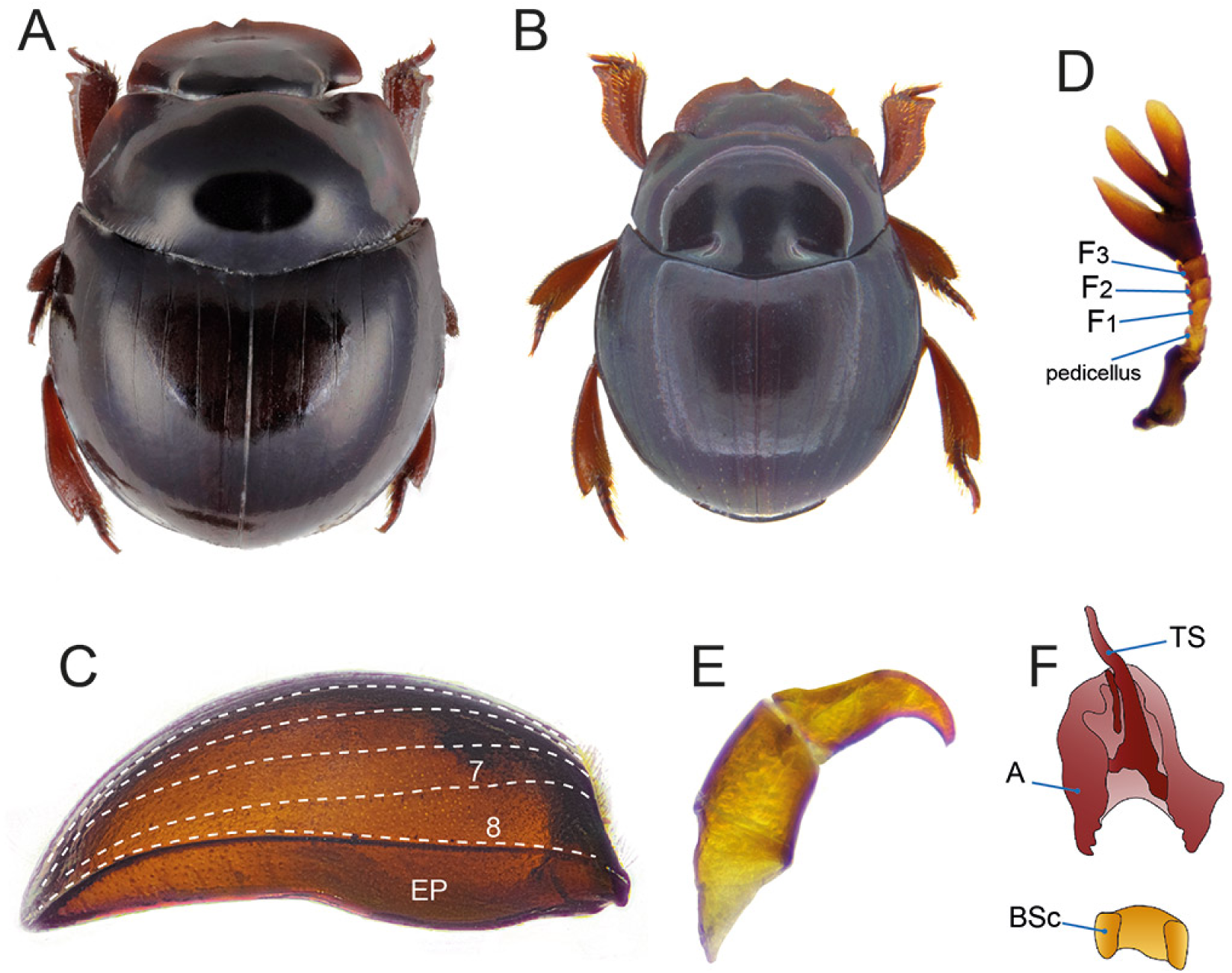
Diagnostic morphological characters of Haroldiini *trib. nov.* (A) *Haroldius brendelli*, habitus; (B) *H. ceylonicus* (=*Phaedotrogus ceylonicus*), habitus (picture by G. Cuccodoro); (C) *H. brendelli*, elytron showing lateral striae (7–8) and very wide epipleuron; (D) *H. brendelli*, antenna with 8 segments (F1–F3 indicate non-club flagellomeres, which in most other Scarabaeinae are 4 instead of 3); (E) *H. lassallei*, aedeagus in lateral view; (F) *H. lassallei*, endophallites.

**Type genus**: *Haroldius* Boucomont, 1914.

### Genus included

*Haroldius* Boucomont, 1914 (40 valid species; Afrotropical, Indomalayan, Palaearctic).

=*Afroharoldius* Janssens, 1949
=*Cyclotrogus* Wasmann, 1918
=*Formicdubius* Philips & Scholtz, 2000
=*Phaedotrogus* Paulian, 1985 ***syn. nov*.**
=*Ponerotrogus* Silvestri, 1924

**Diagnosis**. The tribe is defined by the following unique combination of characters: 1) accessory endophallites consisting of an axial endophallite with thin medial lobe and BSc (Fig. 14F); 2) elytron with 8 striae (Fig. 14C); 3) absence of elytral carinae except for the epipleural one (Fig. 14C); 4) epipleuron very wide (Fig. 14C); 5) antennae with 8 segments (Fig. 14D).

**Description.**

Very small sized beetles (ca. 1.5–3.3 mm); body lenticular, very convex.

*Head*. Anterior clypeal margin usually with 2 teeth, rarely without; frontoclypeal sulcus absent; head surface without protrusions. Antennae with 8 antennomeres.

*Pronotum*. Usually subtrapeziodal, convex, surface punctate or striolate; pronotum with or without trichomes below posterolateral angles; lateral pronotal carina entire.

*Elytra and wings*. Elytra with 8 striae; elytron not carinated except for epipleural carina; epipleuron most often very wide. Mesoscutellum concealed. Metathoracic wings present.

*Legs*. Protibia with 1 or 2 teeth; protibial distal margin often transversely truncated; protarsus present; uncus absent; fore leg without trochanterofemoral pit. Meso- and metatibiae strongly expanded proximo-distally. Tarsal claws not toothed.

*Ventral body surface*. Anterior region of hypomeron depressed; anterior hypomeral carina stretching towards lateral margin of hypomeron; posterior longitudinal hypomeral carina absent.

*Tergite VIII*. Wider than long, evenly convex.

*Genitalia*. Parameres symmetrical, strongly curved ventrally (Fig. 14E). Endophallites: A and BSc present; SA, FLP, SRP and LC absent; presence of raspulae basally.

**Distribution and biology**. The tribe is found in the Afrotropical (including Madagascar), Indomalayan and Palaearctic realms. Most species are myrmecophiles, with some of them possibly having termitophilous, saprophagous or mycophagous habits. They inhabit both forests and open habitats and are often found in the leaf litter and in ant nests (Daniel et al., 2021b; Krikken and Huijbregts, 2006; Krell, 2010; Montreuil, 2010).

**Remarks**. The tribal placement of *Haroldius* has been particularly troubled historically. The genus was placed within different tribes by different authors based on morphology: Onthophagini (Krell, 2010; Philips, 2016), including its synonym Alloscelini (see Onthophagini section) (Janssens, 1949; Balthasar, 1963); Canthonini (Krikken and Huijbregts, 2006; Branco, 1996); and Ateuchini (Montreuil, 2010). More recently, the genus was designated as *incertae sedis* as it does not fit any currently recognised tribe (Tarasov and Dimitrov, 2016; Daniel et al., 2021b). This systematic uncertainty was probably due to the derived morphology of the genus, which comprises very small, globose species adapted to myrmecophyly—*Haroldius* comprises the only known Scarabaeinae bearing trichomes, probably used to interact with their hosts (Krell, 2010).

Finally, recent molecular analyses shed light on the placement of the genus, with all evidence suggesting it is closely related to the coptorhinomorphan Elassocanthonini and Endroedyolini (see Discussion) (Fig. 1). Tarasov and Dimitrov (2016) recovered it as sister to Endroedyolina, although with low bootstrap support, while Odontolomina branched out more basally within the clade. Similarly, our UCE-based reconstructions found full support for *Haroldius* as sister to Endroedyolini+Elassocanthonini. Moreover, a re-evaluation of the morphology of *Haroldius* strongly confirms molecular findings.

The genus shares with its relatives the following unique combination of synapomorphies: 1) an axial endophallite with a thin medial lobe (modified in Odontolomina); 2) the presence of a BSc; 3) the absence of other accessory endophallites; 4) a very wide epipleuron. In particular, the first three characters are unique to all the members of the Coptorhinomorpha clade, also comprising Coptorhinini and Paraphytini. On the other hand, Haroldiini are well differentiated from their relatives by several characters: 1) the presence of 8 (instead of 9) elytral striae; 2) the presence of 8 (instead of 9) antennomeres; 3) the small, lenticular body; 4) an adaptation to myrmecophily or termitophily. Relying on these results, we confidently establish Haroldiini *trib. nov*.

Paulian (1985) described the genus *Phaedotrogus* Paulian, 1985 for a single species, *P. ceylonicus* (Balthasar, 1972), sharing several characters with *Haroldius* and *Ponerotrogus* Silvestri, 1924 (now considered a junior synonym of *Haroldius*). Despite noticing several similarities with *Haroldius*, Paulian (1985) stated that *P. ceylonicus* clearly differs from the latter, yet without giving clear arguments for this statement. Since then, the genus has virtually never been discussed by any author (Daniel et al., 2021b; Krell, 2010; Krikken and Huijbregts, 2006). We examined the external morphology of type specimens of *P. ceylonicus* and we could not find any relevant feature distinguishing it from *Haroldius*. Body and leg shape fall within the known variation of the genus, and elytral striation perfectly matches it. Therefore, we propose the following synonymy: *Haroldius* Boucomont, 1914 = *Phaedotrogus* Paulian, 1985 *syn. nov*. Some authors pointed out the need of re-evaluating the generic classification of *Haroldius* and its many synonyms, hinting at the possibility of re-splitting the genus (Daniel et al., 2021b; Krikken and Huijbregts, 2006). However, we argue that the group is almost surely a monophyletic lineage based on morphology. Overall, its current classification is satisfactory and does not require unnecessary oversplitting based on minor differences which most likely represent the species-specific outcomes of strong selective pressures related to myrmecophily.

### Heliocoprini *trib. nov*

(Figs. 15, 3L)

**Figure 15.**
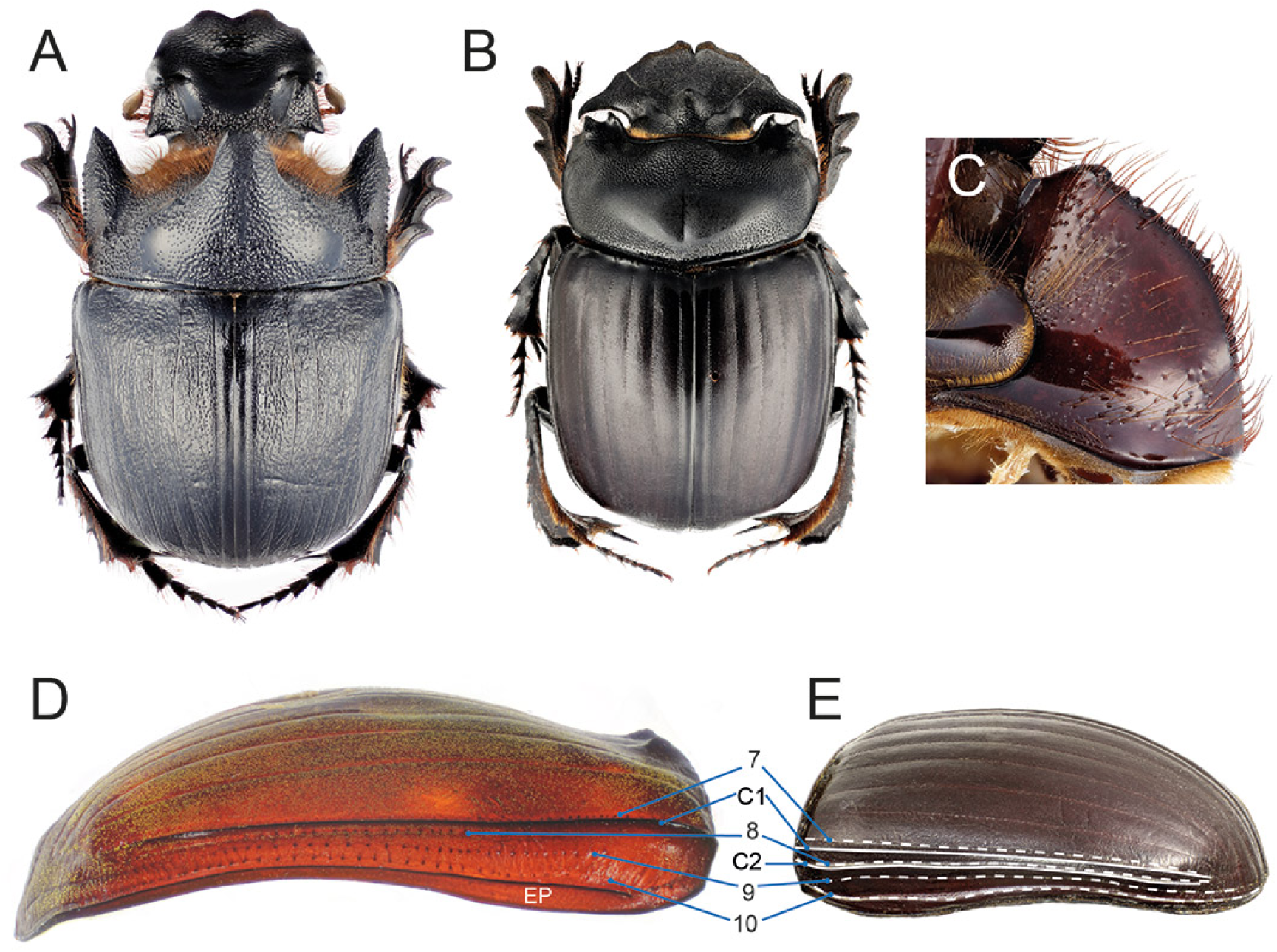
Diagnostic morphological characters of Heliocoprini *trib. nov.* (A) *Heliocopris gigas*, habitus; (B) *Synapsis tridens*; (C) *H. gigas*, hypomeron with no trace of anterior hypomeral carina; (D) *Heliocopris* sp. and (E) *S. tridens*, elytra showing lateral striae (7–10) and carinae (C1, C2; epipleural carina not marked).

**Type genus**: *Heliocopris* Hope, 1837.

**Genera incpluded:**

*Heliocopris* Hope, 1837 (57 valid species; Afrotropical, Indomalayan).

*Synapsis* Bates, 1868 (23 valid species; Indomalayan, Palaearctic).

**Diagnosis**. The tribe is defined by the following unique combination of characters: 1) elytron with 10 striae (Figs. 15D–E); 2) striae 8–10 placed on pseudoepipleuron (Figs. 15D–E); 3) anterior hypomeral carina absent (Fig. 15C) or, if carina present, elytron with a second carina between stria 8 and 9 (Fig. 15E).

**Description.**

Large to very large beetles (ca. 21–65 mm); body from moderately to very convex.

*Head*. Anterior clypeal margin with or without two teeth; head often carinated, tuberculated or horned in correspondence with frontoclypeal sulcus, which is absent; head vertex most often devoid of protrusions. Antennae with 9 antennomeres.

*Pronotum*. Granulated or punctured, either with conspicuous protrusions or carinae (*Heliocopris*), or evenly convex (*Synapsis*); mediolateral regions with a fovea; lateral pronotal carina entire, double in *Synapsis*.

*Elytra and wings*. Elytra with 10 visible striae; elytron carinate between striae 7 and 8; striae 8–10 placed on pseudoepipleuron; elytron carinated between striae 8 and 9 in *Synapsis*, not so in *Heliocopris*. Mesoscutellum concealed. Metathoracic wings present.

*Legs*. Protibia with 3 teeth; protibial distal end obliquely truncated; male uncus absent. Meso- and metatibiae strongly expanded proximo-distally; metatibiae strongly bent medially in some *Synapsis*; dorsal metatibial surface with two transverse carinae. Tarsal claws not toothed.

*Ventral body surface*. Anterior hypomeral carina present (a few species of *Synapsis*) or absent (*Heliocopris*, most *Synapsis*), if present stretching towards lateral margin of pronotum; posterior longitudinal hypomeral carina absent.

*Tergite VIII*. Wider than long, evenly convex.

*Genitalia*. Parameres symmetrical, elongated, triangular. Endophallites: A, SA, FLP and SRP present; LC absent.

**Distribution and biology**. *Heliocopris* is widespread in the Afrotropical realm with a few Indomalayan representatives, while *Synapsis* is mostly found in the Indomalayan realm with a few species reaching the Palaearctic (Pokorný et al., 2009; Zídek and Pokorný, 2010). *Heliocopris* are coprophagous tunnellers and the largest species are often associated with the dung of large non-ruminant mammals (Davis et al., 2008). Little is known about the biology of *Synapsis*, but they likely share similar habits with their sister genus.

**Remarks**. *Heliocopris* and *Synapsis* have long been included in the Coprini (Hope, 1837; Bates, 1868; Balthasar, 1963; Montreuil, 1998), although Edmonds and Halffter (1978) suggested their placement within the Dichotomiini, which was followed by some authors (Davis et al., 2008). The two genera were then classified as *incertae sedis* as multiple evidence showed that they are in fact unrelated to *Copris* and allies (present analyses; Gunter et al., 2016, 2026; Lopes et al., 2024; Tarasov and Génier, 2015; Tarasov and Dimitrov, 2016).

A close relation between *Heliocopris* and *Synapsis* was suggested by Génier (1996) and indicated in the morphological works by Bai et al. (2011) and Montreuil (1998), although no further evidence was added by the broad phylogeny of Tarasov and Solodovnikov (2011), which did not examine *Synapsis*. Here, we were able to include both genera in molecular analyses for the first time, which resulted in a fully supported sister relationship (Fig. 2) (see also Tarasov and Dimitrov, 2016).

A re-evaluation of morphological traits further corroborates this hypothesis. Both genera share the presence of 10 elytral striae of which the three most lateral (8—10) are placed on the pseudoepipleuron. This character is rare in the Scarabaeinae, being known only in the Australasian tribes Boletoscapterini and Mentophilini (genera *Amphistomus*, *Aptenocanthon*, *Diorygopyx*, *Labroma*, *Onthobium*, *Paronthobium*, *Saphobius*, *Tesserodon*) (Gunter et al., 2026). However, the two tribes are completely unrelated to Heliocoprini based on all available evidence (present analyses; Montreuil, 1998; Tarasov and Génier, 2015; Tarasov and Dimitrov, 2016), and can anyway be separated from Heliocoprini by the constant presence of an anterior hypomeral carina. The latter is present only in some species of *Synapsis* (Zídek and Pokorný, 2010), which can be easily distinguished from Boletoscapterini and Mentophilini by the presence of a second elytral carina between striae 8 and 9 and/or by the presence of a dense fringe of long setae along the margin of the hypomeral cavity.

Overall, the phylogenetic and morphological distinctiveness of Heliocoprini *trib. nov.* fully justifies its status as a separate tribe.

### Janssensantini *trib. nov*

(Fig. 16)

**Figure 16.**
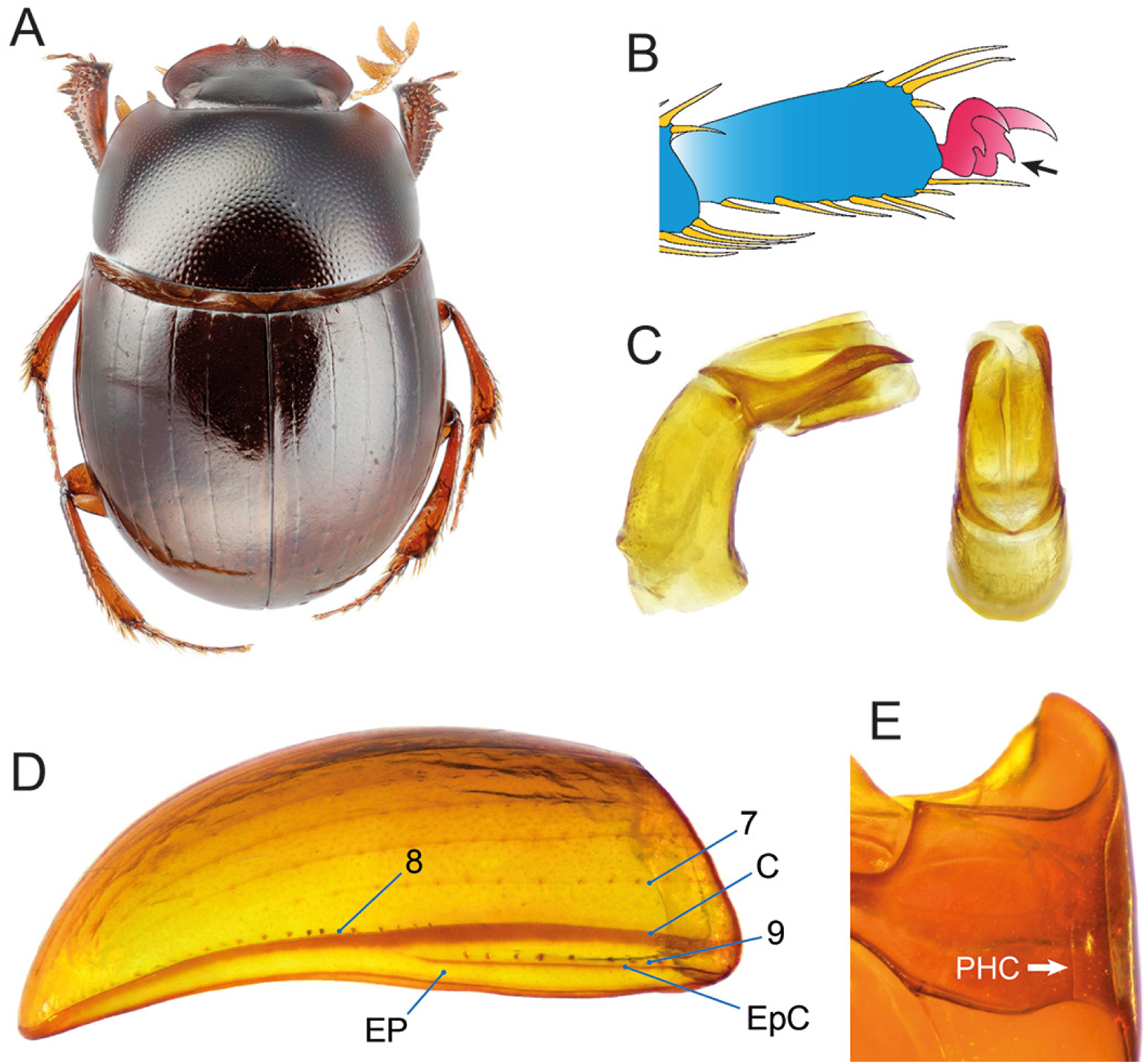
Diagnostic morphological characters of Janssensantini *trib. nov.* (A) *Janssensantus inexpectatus*, habitus; (B) *J. pauliani*, detail of mesotarsomere with arrow pointing at toothed claws; (C) *J. pauliani*, aedeagus in lateral (left) and dorsal (right) views; (D) *J. pauliani*, elytron showing lateral striae (7–9) and carinae (C1, C2, EpC); (E) *J. pauliani*, hypomeron showing posterior hypomeral carina.

**Type genus**: *Janssensantus* Paulian, 1976.

**Genus included:**

*Janssensantus* Paulian, 1976 (3 valid species; Afrotropical).

**Diagnosis**. The tribe is defined by the following unique combination of characters: 1) elytron with 9 visible striae, stria 9 located on pseudoepipleuron (Fig. 16D); 2) epipleural carina effaced posteriorly (Fig. 16D); 3) posterior longitudinal hypomeral carina present (Fig. 16E); 4) tarsal claws toothed (Fig. 16B).

**Description**.

Small sized beetles (3.5-5.0 mm); body oval, convex, integument shiny.

*Head*. Surface evenly punctate. Anterior margin of clypeus with two teeth, clypeus slightly notched laterally to teeth; head margin evenly rounded at genoclypeal suture; frontoclypeal sulcus absent. Antennae with 9 segments.

*Pronotum*. Evenly and rather densely covered with simple punctures; lateral pronotal carina entire.

*Elytra and wings*. Elytron with 9 visible striae; elytron carinate between striae 8 and 9; epipleural carina effaced posteriorly; elytron moderately slanted inwards laterally to the first carina. Mesoscutellum concealed. Metathoracic wings either present or absent.

*Legs*. Protibia with 3 teeth, distal margin approximately transversely truncated; uncus present in males; protarsus present in both sexes. Meso- and metatibiae evenly expanded proximo-distally. Ventral margin of male metafemur angulate on distal third of length. Tarsal claws toothed proximally.

*Ventral body surface*. Anterior region of hypomeron depressed; anterior hypomeral carina present, stretching towards lateral margin of hypomeron; posterior longitudinal hypomeral carina present. Metaventrite evenly convex; mesometaventral sulcus strongly arched.

*Tergite VIII*. Evenly convex.

*Genitalia*. Parameres either symmetrical or asymmetrical, with left paramere slightly longer. Endophallites: LC present; SA, A and FLP endophallites present; SRP with ring-shaped part.

**Distribution and biology**. The tribe is restricted to the Eastern Arc Mountains in Tanzania (*Janssensantus inexpectatus* Paulian, 1976 and *J. pauliani* Scholtz & Howden, 1987), where it occurs in rainforest (Scholtz and Howden, 1987), and to the South East African Montane Archipelago in Mozambique, where it is associated with loamy sand under dense woodland and bamboo forest in the foothills of Mount Mabu (Bayliss et al., 2024; Daniel et al., 2023). It is also known from degraded miombo in Zambia (*J. alatus* Josso, 2022) (Josso, 2022; J.-F. Josso, personal commmunication). A few specimens were attracted to pig and human dung, as well as to injured diplopods (Davis et al., 2008; Josso, 2022; GM, personal observation in Tanzania). An ongoing revision of the genus hints at the existence of additional undescribed, microendemic rainforest species (Montanaro *et al*., in preparation).

**Remarks**. The genus *Janssensantus* was traditionally placed in Canthonini (Paulian, 1976; Scholtz and Howden, 1987). The phylogenies by Mlambo et al. (2014) and Tarasov and Dimitrov (2016) found it to be closely related to another relictual rainforest specialist, *Tanzanolus* (Tanzanolini), yet this relationship is not confirmed by our reconstruction, which always found *Janssensantus* in very isolated positions as sister to large clades (see Fig. 1 and supplementary files S2–S3).

Despite its somewhat ordinary “canthonine” appearance, *Janssensantus* has a very distinctive morphology, being the only scarabaeine known to us in which the epipleural carina disappears completely on the posterior half of the elytron (Fig. 16), so that the epipleuron and pseudoepipleuron appear to be fused. Moreover, Janssensantini have a proximal denticle on tarsal claws, a character only shared, among Afro-Eurasian tribes, with the totally unrelated Bohepilissini. The creation of Janssensantini *trib. nov.* is therefore robustly justified both on a molecular and morphological basis.

### Macroderini *trib. nov*

(Figs. 17, 3C)

**Figure 17.**
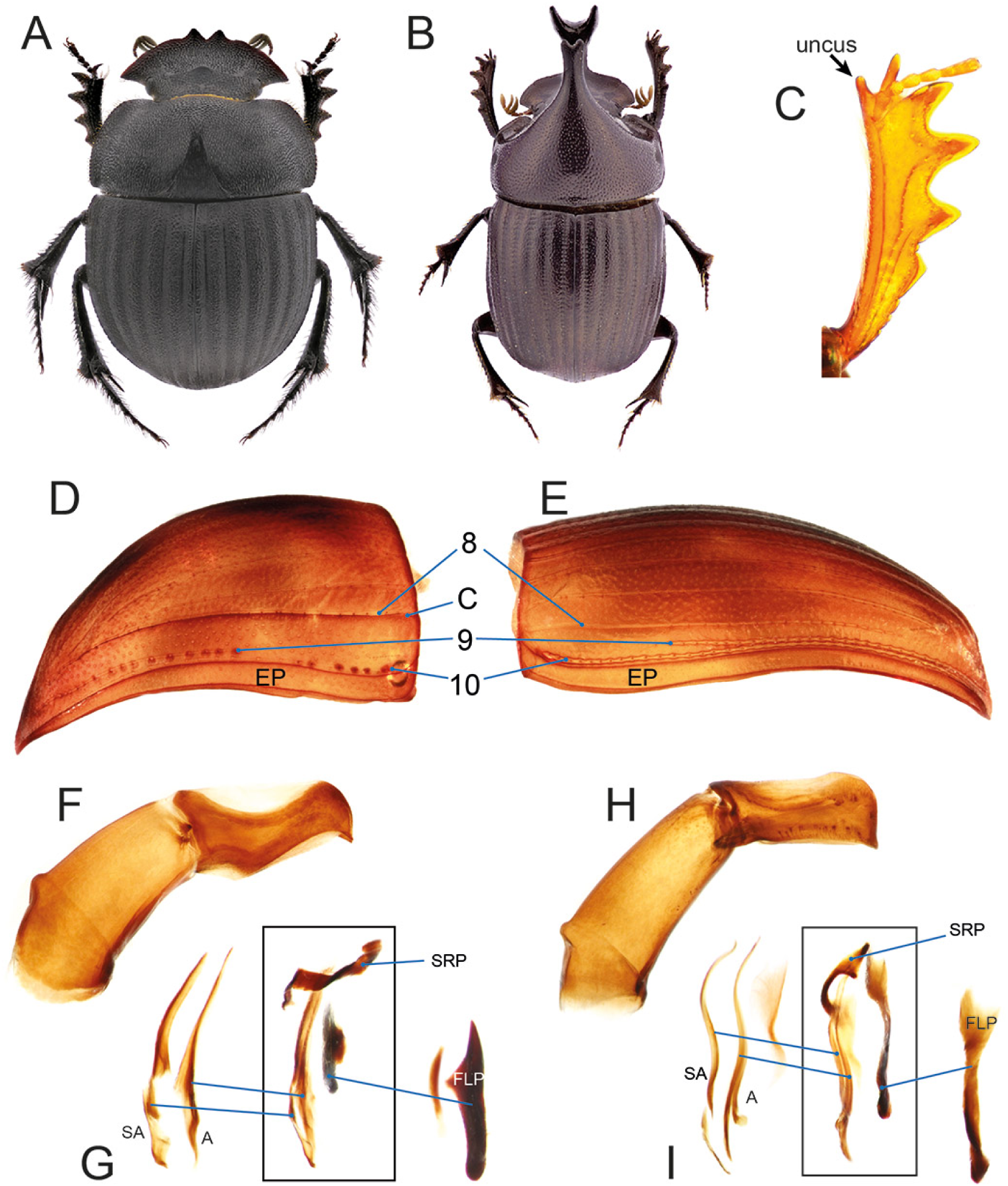
Diagnostic morphological characters of Macroderini *trib. nov.* (A) *Macroderes foveatus*, abitus (picture by C. Deschodt); (B) *Xinidium dewitzi*, habitus (picture by C. Deschodt); (C) *X. dentilabris*, male protibia showing uncus; (D) *M. mutilans* and (E) *X. dentilabris*, elytra showing lateral striae (8–10) and carinae (C; epipleural carina not marked); (F) *M. mutilans*, aedeagus in lateral view; (G) *M. mutilans*, endophallites; (H) *X. dentilabris*, aedeagus in lateral view; (I) *X. dentilabris*, endophallites.

**Type genus**: *Macroderes* Westwood, 1842.

**Genera included:**

*Macroderes* Westwood, 1842 (24 valid species; Afrotropical).

*Xinidium* Harold, 1869 (4 valid species; Afrotropical).

**Diagnosis**. The tribe is defined by the following unique combination of characters: 1) elytron with 10 visible striae (Figs. 17D–E); 2) elytral stria 9 not reaching the anterior margin of elytron (Figs. 17D–E); 3) parameres, relatively wide, convex dorsally and rounded apically, pointed distoventrally (Figs. 17F, 17H); 4) ventral part of axial endophallite significantly narrower than remaining portion of sclerite (Figs. 17G, 17I).

**Description.**

Medium sized beetles (7.0–13.7 mm); body convex, hemispherical to elongated.

*Head*. Anterior margin of clypeus with two teeth, strongly protruding to horn-like in *Xinidium* males; head margin evenly rounded on the whole; frontoclypeal sulcus carinate or tuberculate. Antennae with 9 segments.

*Pronotum*. Surface more or less densely punctate; with horn-like protrusions or carinae *Xinidium*, evenly convex or with shallow modifications in *Macroderes*; lateral pronotal carina entire.

*Elytra and wings*. Elytron with 10 striae; elytron carinate laterally to stria 8 in *Macroderes*, not carinate in *Xinidium*; stria 9 not reaching anterior margin of elytron. Mesoscutellum concealed. Metathoracic wings either present (*Xinidium*) or absent (*Macroderes*).

*Legs*. Protibia with 4 teeth; uncus present in males (Fig. 17C); protarsus present in both sexes. Meso- and metatibiae strongly and evenly expanded proximo-distally. Tarsal claws not toothed.

*Ventral body surface*. Anterior region of hypomeron not depressed; anterior hypomeral carina present, stretching towards lateral margin of hypomeron; posterior longitudinal hypomeral carina absent.

*Tergite VIII*. Evenly convex.

*Genitalia*. Parameres symmetrical, relatively wide, convex dorsally and rounded apically, pointed distoventrally. Endophallites: LC absent; SA and FLP endophallites present; A endophallite with ventral part significantly narrower than remaining portion of sclerite; SRP with ring-shaped part.

**Distribution and biology**. Macroderini are endemic to the Afrotropical realm, being largely restricted to South Africa, with only one species of *Xinidium* reaching eastern Zimbabwe (Abdalla et al., 2018; Cambefort, 1985). They are tunnellers feeding on dung with a few records from carrion for *Macroderes*. Apparently both genera comprise nocturnally active species (Davis et al., 2008).

**Remarks**. Historically, *Macroderes* and *Xinidium* have been placed in the highly polyphyletic tribe Dichotomiini due to the general “tunneller” habitus and the lack of transverse carinae on the dorsal margin of mesotibiae (Montreuil, 1998), although they were considered Coprini by some authors (Cambefort, 1985; Davis et al., 2008). More recently, the two genera were moved to *incertae sedis* as they did not match any described tribe (Tarasov and Dimitrov, 2016).

The morphological phylogeny by Tarasov and Génier (2015) found the two genera to be sisters and nested within a clade comprising Catharsiini, Onitini and Dwesasilvasedini. This is fully supported by our analyses, as well as by all previous molecular studies (Gunter et al., 2026; Mlambo et al., 2015; Sole and Scholtz, 2010; Tarasov and Dimitrov, 2016), which place *Macroderes* and *Xinidium* in the Onthophagomorpha clade (Fig. 2). On the other hand, earlier molecular works based on a few loci found that the two genera did not form a monophyletic group: Sole and Scholtz (2010) recovered the relationship *Macroderes*+(*Dwesasilvasedis*+*Xinidium*); Mlambo et al. (2015) found *Xinidium*+(*Aphengoecus*+*Macroderes*); and Tarasov and Dimitrov (2016) found them within an even more complicated topology. However, all of our UCEs-based reconstructions fully support a sister relationship between *Macroderes* and *Xinidium*.

Morphologically, the monophyly of the new tribe is corroborated by two unique synapomorphies: the shape of parameres, and the shape of the ventral part of the axial endophallite. The proximity of the Macroderini to the Dwesasilvasedini and Onitini is confirmed by molecular analyses, and it is further supported by a common configuration of elytral striation: the usual presence of 10 striae, of which the 9th does not reach the anterior elytral margin, and a pseudoepipleural carina between striae 8 and 9 (Figs. 11, 17 and 19). An exception to this pattern is found in *Xinidium*, which, similarly to some species of Onitini, lacks the pseudoepipleural carina. This genus is also well-characterised by having evolved a strong sexual dimorphism, with major males having conspicuous protrusions on pronotum and clypeus.

### Nesovinsoniini *trib. nov*

(Fig. 18)

**Figure 18.**
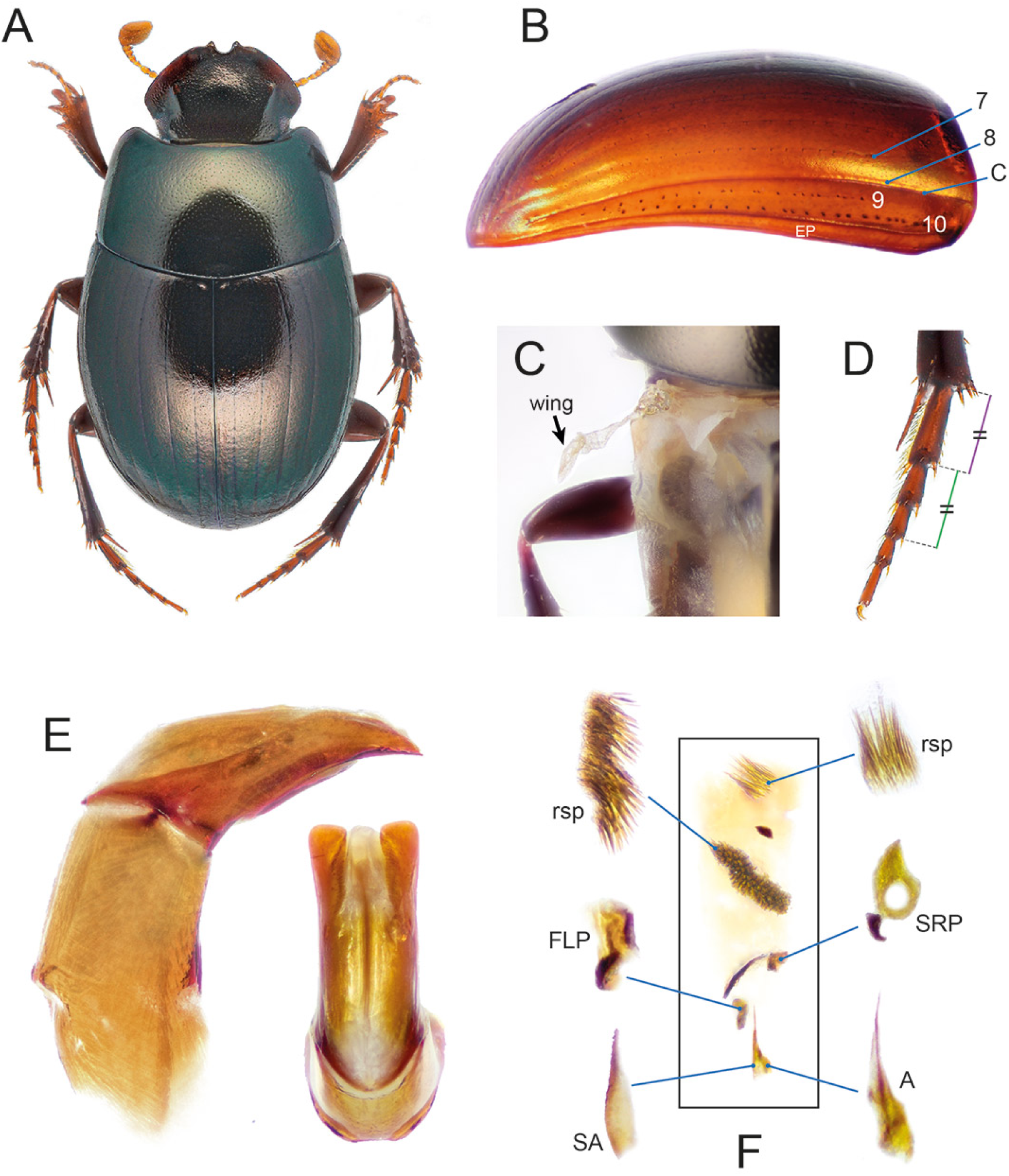
Diagnostic morphological characters of Nesovinsoniini *trib. nov.* *Nesovinsonia vinsoni*: (A) habitus; (B) elytron showing lateral striae (7–10) and carinae (C; epipleural carina not marked); (C) atrophied metathoracic wing; (D) metatarsus showing that the length of the first tarsomere equals the sum of the lengths of the following two segments; (E) aedeagus in lateral (left) and dorsal (right) views; (F) endophallites.

**Type genus**: *Nesovinsonia* Martínez & Pereira, 1958.

**Genus included:**

*Nesovinsonia* Martínez & Pereira, 1958 (1 valid species; Madagascan (Mauritius)).

**Diagnosis**. The tribe is defined by the following unique combination of characters: 1) elytron with 10 striae (Fig. 18B); 2) striae 9 and 10 placed on pseudoepipleuron (Fig. 18B); 3) protibiae with 3 external teeth (Fig. 18A); 4) protibial uncus absent (Fig. 18A); 5) metatarsomere 1 as long as 2 and 3 combined (Fig. 18D).

**Description.**

Small sized beetles (4.7–5.5 mm); body oval, convex.

*Head*. Anterior margin of clypeus with 2 teeth separated by a deep rounded notch and slightly notched laterally to the teeth; frontoclypeal sulcus absent; head surface without protrusions. Antennae with 9 antennomeres.

*Pronotum*. Subtrapeziodal, evenly punctate; lateral pronotal carina entire.

*Elytra and wings*. Elytron with 10 striae; carinated between striae 8 and 9 (pseudoepipleural carina). Mesoscutellum concealed. Metathoracic wings atrophied (Fig. 18C).

*Legs*. Protibia with 3 teeth; protibial distal margin obliquely truncated; protarsus present; uncus absent. Meso- and metatibiae evenly expanded proximo-distally; meso- and metatarsomeres 1 approximately as long as following two tarsomeres combined. Tarsal claws slightly toothed basally.

*Ventral body surface*. Anterior region of hypomeron depressed; anterior hypomeral carina stretching towards lateral margin of hypomeron; posterior longitudinal hypomeral carina absent.

*Tergite VIII*. Wider than long, evenly convex.

*Genitalia*. Parameres symmetrical, elongated, apex feebly curved ventrally (Fig. 18E). Endophallites (Fig. 18F): A, SA, FLP and SRP present; A and SA not fused; SRP with ring-shaped part; LC absent; presence of 2 raspulae and a small additional sclerite basally.

**Distribution and biology**. The only known species of the tribe, *Nesovinsonia vinsoni* (Paulian, 1939), is endemic to Mauritius in the Mascarene island archipelago. It was recently collected using pitfall traps baited with chicken dung (Lopes et al., 2023).

**Remarks**. *N. vinsoni* was originally described by Paulian (1939) as a species of the canthonine genus *Phacosoma* Boucomont, 1914 (now synonym of *Ochicanthon* Vaz-de-Mello, 2003, tribe Epactoidini). Martínez and Pereira (1959) placed it in a new monotypic genus, *Nesovinsonia*, differing from *Phacosoma* and related genera by the number of elytral striae and other characters. The genus was kept in Canthonini until Tarasov and Dimitrov (2016) transferred it to *incertae sedis*.

Morphologically, *Nesovinsonia* superficially resembles some Epactoidini, especially the genus *Ochicanthon* and *Epactoides giganteus* Rossini, Vaz-de-Mello & Montreuil, 2021, the latter sharing with *Nesovinsonia* the presence of 10 full elytral striae (usually 9 in Epactoidini; Rossini et al., 2021). However, *Nesovinsonia* has several characters that set it apart from Epactoidini, especially the configuration of endophallites: FLP present (absent in Epactoidini), and A and SA separated (fused in *Epactoides*) (Rossini et al., 2022). The collection of fresh *Nesovinsonia* specimens allowed us to sequence this genus for the first time. Molecular analyses give no evidence for a close relationship with Epactoidini, while they fully support a sister relationship with the relict African endemic *Tanzanolus* (Tanzanolini), with which *Nesovinsonia* forms a rather isolated clade (Fig. 1). Molecular divergence between Nesovinsoniini and Tanzanolini is deep and paralleled by the lack of meaningful morphological synapomorphies uniting the two groups. For these reasons, the creation of two distinct tribes for these taxa is largely justified.

### Onitini Laporte, 1840

(Fig. 19)

**Figure 19.**
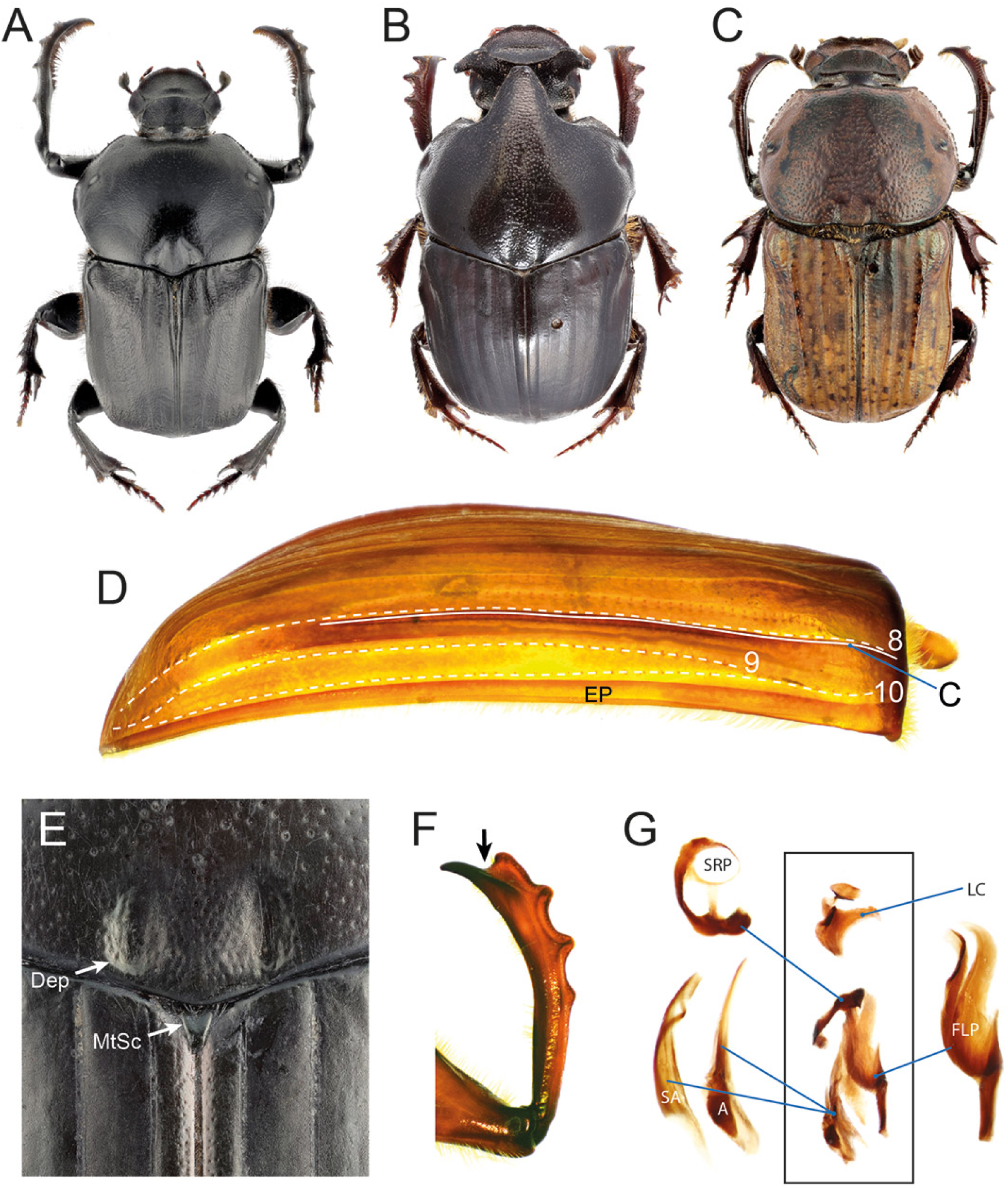
Diagnostic morphological characters of Onitini. (A) *Onitis belial*, habitus; (B) *Bubas bison*, habitus; (C) *Cheironitis pamphilus*, habitus; (D) *Onitis* sp., elytron showing lateral striae (8–10) and carina (C; epipleural carina not marked); (E) *Onitis subopacus*, detail of thorax showing pronotal posterior depressions (Dep) and scutellum (MtSc); (F) *O. subopacus*, male protibia; (G) *Onitis* sp., endophallites.

**Type genus**: *Onitis* Fabricius, 1798.

**Genera included:**

*Acanthonitis* Janssens, 1937 (4 valid species; Afrotropical);

*Allonitis* Janssens, 1936 (1valid species; Afrotropical);

*Anonychonitis* Janssens, 1950 (1valid species; Afrotropical);

*Aptychonitis* Janssens, 1937 (2 valid species; Afrotropical);

*Bubas* Dejean, 1833 (3 valid species; Palaearctic);

*Cheironitis* Lansberge, 1875 (24 valid species; Afrotropical, Indomalayan, Palaearctic);

*Gilletellus* Janssens, 1937 (2 valid species; Afrotropical);

*Heteronitis* Gillet, 1911 (5 valid species; Afrotropical);

*Janssensellus* Cambefort, 1975 (2 valid species; Afrotropical);

*Kolbeellus* Jacobson, 1906 (1 valid species; Afrotropical);

*Lophodonitis* Janssens, 1938 (2 valid species; Afrotropical);

*Megalonitis* Janssens, 1937 (2 valid species; Afrotropical);

*Neonitis* Péringuey, 1901 (2 valid species; Afrotropical);

*Onitis* Fabricius, 1798 (179 valid species; Afrotropical, Australian, Indomalayan, Palaearctic; introduced in Nearctic realm);

*Platyonitis* Janssens, 1942 (4 valid species; Afrotropical);

*Pleuronitis* Lansberge, 1875 (4 valid species; Afrotropical);

*Pseudochironitis* Ferreira, 1977 (2 valid species; Afrotropical);

*Tropidonitis* Janssens, 1937 (1 valid species; Afrotropical).

**Diagnosis**. The tribe is defined by the following unique combination of characters: 1) posterior margin of pronotum with two medial depressions (Fig. 19E); 2) mesoscutellum visible (Fig. 19E); 3) elytron with 10 striae, stria 9 sometimes indistinct (Fig. 19D); 4) protarsus absent in males (Fig. 19F); 5) protibial spur absent in males (Fig. 19F); 6) axial endophallite with sclerotised process extending ventrally (Fig. 19G).

**Description**.

Medium to very large sized beetles (10–40 mm); elongated, moderately convex to flattened.

*Head*. Head margin variously shaped, with variable number of clypeal teeth; clypeal carina either present or absent; frontoclypeal carina present; vertex carinate, tuberculate or without protrusions. Antennae with 9 antennomeres.

*Pronotum*. Variously punctured or granulated, with protrusions or not; posterior margin with two depressions; lateral pronotal carina entire.

*Elytra and wings*. Elytron with 10 visible striae, in some species stria 9 absent or effaced; elytron usually carinated laterally to stria 8, carina absent in *Aptychonitis* and some *Cheironitis*. Mesoscutellum visible. Metathoracic wings present.

*Legs*. Protibia with 4 teeth; uncus present in males; protibial spur absent in males (but see Remarks), present in females (fused with protibia in *Allonitis*); protarsus absent in males, present or absent in females. Meso- and metatibiae strongly and evenly expanded proximo-distally. Male legs often modified and with variously shaped protrusions. Tarsal claws not toothed.

*Ventral body surface*. Anterior region of hypomeron not depressed; anterior hypomeral carina absent; posterior longitudinal hypomeral carina absent. Metaventrite either evenly convex or with various depressions; prosternum either simple or with variously shaped protrusions.

*Tergite VIII*. Elongated in males, squeezing tergite VII medially.

*Genitalia*. Parameres either symmetrical (most genera) or asymmetrical (*Bubas*), elongated, posterior end usually sharp and bent ventrally. Endophallites: LC present; SA endophallite present; axial endophallite with sclerotised process extending ventrally; FLP endophallite usually big, long, comprising a variable number of distinct lobes, reduced to an ill-defined sclerotisation in some species of *Onitis*; SRP with ring-shaped part.

**Distribution and biology**. Onitini are widespread in Afro-Eurasia, with few species reaching Australasia and some others introduced into the Nearctic and Australasian realms. They occupy various ecosystems, ranging from dry habitats (*e.g.*, *Cheironitis*) to mountain rainforests (*e.g.*, *Lophodonitis*). Most species are attracted to herbivore or, rarely, omnivore dung, with a number more or less specialised to non-ruminant, herbivore dung, especially that of elephant and rhinoceros or even zebra (Davis et al., 2020; Moretto, 2010).

**Remarks**. Onitini are a molecularly strongly supported and morphologically well characterised tribe (present analyses; Ayivi et al., 2021; Breeschoten et al., 2016; Cheng et al., 2021; Gunter et al., 2026; Guo et al., 2022; Mlambo et al., 2015; Monaghan et al., 2007; Tarasov and Dimitrov, 2016; Tarasov and Génier, 2015; Villalba et al., 2002). Historically, the most important morphological characters used for its definition include the 4-toothed protibia, the absence of protarsus and protibial spur in males, the visible mesoscutellum and the symmetrical depressions on the posterior margin or pronotum (Davis et al., 2008; Janssens, 1937). The diagnosis provided here adds the peculiar shape of the axial endophallite, as well as elytral striation. It must be noticed that the correct number of elytral striae is usually 10, and not 8 as stated by previous authors (*e.g.*, Janssens (1937)), who did not count those placed on the pseudoepipleuron. Nevertheless, in some taxa stria 9 is poorly visible or completely effaced, for example in *Platyonitis* and some *Onitis*. Notably, the genus *Aptychonitis* and some *Cheironitis* (*e.g.*, *C. flabellatus* Boucomont, 1923 and *C. imitator* Balthasar, 1963) lack more or less completely the pseudoepipleural carina.

A remarkable feature of Onitini is the absence of protibial spur in males. This is a rare character among Scarabaeinae, to our knowledge only found in Dwesasilvasedini, in some species of Eurysternini (Génier, 2009), and occasionally in specimens of *Streblopus* Van Lansberge, 1874 (*incertae sedis*) (Cupello et al., 2020). However, the anatomical origin of this trait may be heterogeneous. While in some taxa (Eurysternini, *Streblopus*) this is likely due to a simple loss of the spur, in Onitini (and Dwesasilvasedini?) it may be caused by fusion of the spur with the protibial margin and the *uncus*. In Onitini, this phenomenon may have occurred twice in the genus *Allonitis*, whose females also have an “*uncus*” which, most probably, is in fact a fused protibial spur (Janssens, 1937).

Based on the similarity between the endophallites of Onitini and the American tribe Phanaeini, Zunino (1983) defined three subtribes within Onitini: Onitina, Phanaeina, and Gromphina (now Gromphadina Cupello and Vaz-De-Mello (2013)). This classification was later dismissed, as multiple evidence indicated that Onitina and the genera in the other two subtribes, now placed in the Phanaeini, are not closely related (Philips et al., 2004; Tarasov and Génier, 2015; Tarasov and Dimitrov, 2016; Gillett and Toussaint, 2020). Instead, all evidence indicates that Onitini fall within the Onthophagomorpha clade (Fig. 2).

Lastly, while monophyly of Onitini is well established, several works recovered paraphyly of *Onitis* with respect to other genera (present analyses; Breeschoten et al., 2016; Cheng et al., 2021; Gunter et al., 2026; Guo et al., 2022; Villalba et al., 2002). This is further supported by preliminary observations of male genitalia of several onitine lineages. Findings stress the need for a revision of the systematics of the tribe, hopefully, based on a dedicated, fine-scale phylogeny.

### Onthophagini Streubel, 1846 *sensu novo*

(Figs. 20, 3E)

**Figure 20.**
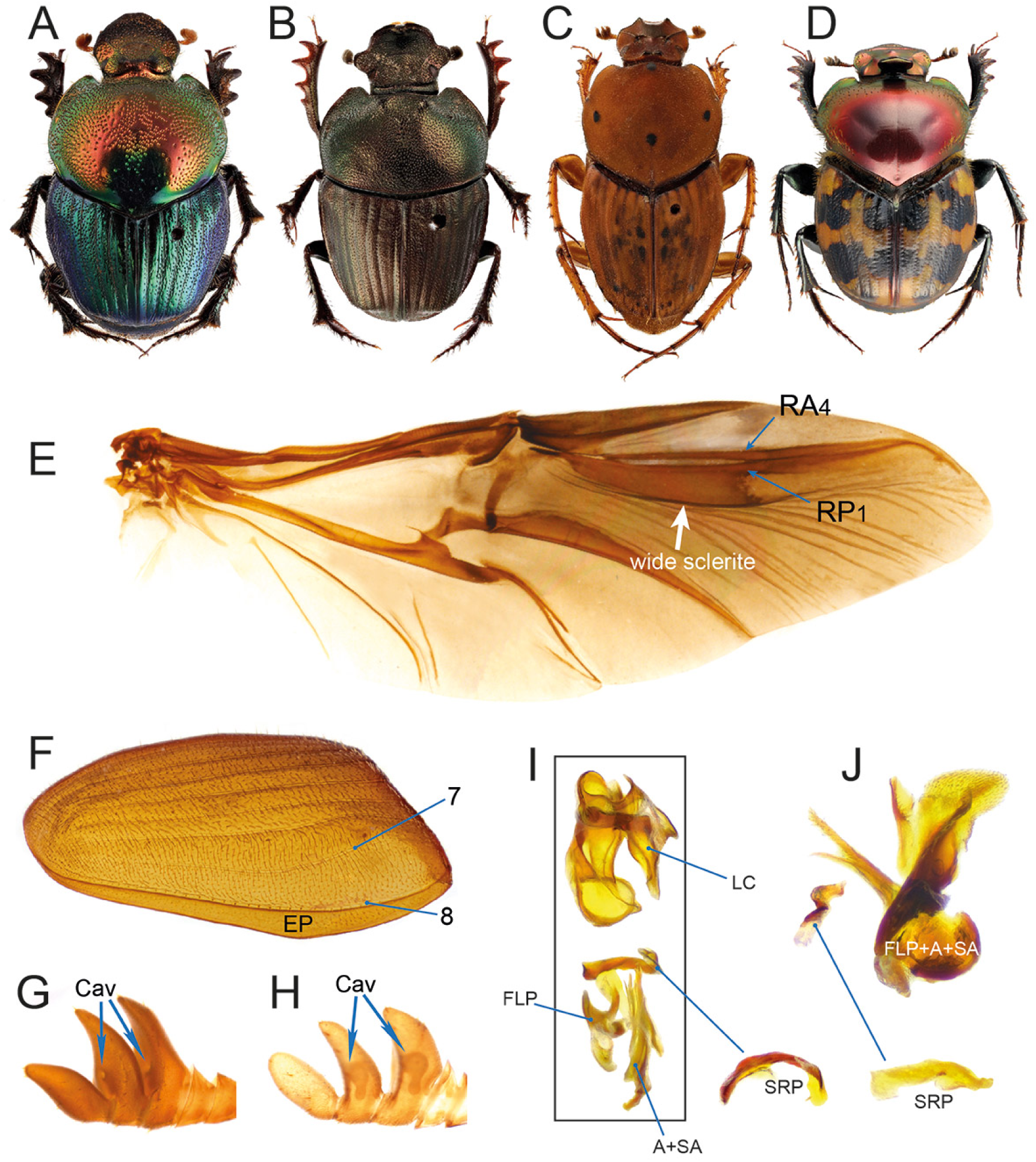
Diagnostic morphological characters of Onthophagini *sensu novo*. (A) *Proagoderus gemmatus*, habitus; (B) *Phalops boschas*, habitus; (C) *Yvescambefortius sarawacus*, habitus; (D) *Helictopleurus splendidicollis*, habitus; (E) *Onthophagus* sp., metathoracic wing showing the wide posterior sclerite of RP1 vein; (F) *Onthophagus* sp., elytron showing lateral striae (7–8); (G) *Diastellopalpus quinquedens* and (H) *Onthophagus* (*Serrophorus*) *seniculus*, antennal clubs showing cavities on distal surface of antennomeres (blue arrows); (I) *Diastellopalpus semirubidus* and (J) *Digitonthophagus* sp., endophallites.

**Type genus**: *Onthophagus* Latreille, 1802.

**Subtribes included:**

**Helictopleurina** Janssens, 1946 ***stat. nov.*** (1 genus, 59 valid species; Afrotropical (Malagasy));

**Oniticellina** Kolbe, 1905 ***stat. nov.*** (24 genera, 205 valid species; Afrotropical, Nearctic, Neotropical, Indomalayan, Palaearctic);

= Attavicinina Philips, 2016 **syn. nov.**
= Drepanocerina Lansberge, 1875 **syn. nov.**
= Liatongina Philips, 2016 **syn. nov.**

**Onthophagina** Streubel, 1846 (48 genera, 3014 valid species; Afrotropical, Australasian, Nearctic, Neotropical, Indomalayan, Palaearctic);

= Alloscelina Janssens, 1949 **syn. nov**.

**Diagnosis**. The tribe is defined by the following unique combination of characters: 1) wing with RP1 having wide posterior sclerite (Fig. 20E); 2) elytron with 8 striae, very rarely with a partial stria between stria 7 and 8 (Fig. 20F); 3) absence of elytral carinae except for the epipleural one (Fig. 20F); 4) first and, often, second club antennomeres with a cavity on their distal surface (Figs. 20G–H); 5) SRP not ring-shaped (Figs. 20I–J); 6) if posterior sclerite of RP1 vein reduced, then accessory endophallites consisting of FLP, SA, A and additional sclerites variously fused together, plus an unfused SRP (Fig. 20J).

**Distribution and biology**. With almost 3300 described species, Onthophagini are the are most diverse tribe of dung beetles. They are tunnelers found on all continents except Antarctica, and inhabit a broad variety of habitats. Trophic habits include coprophagy, necrophagy (including some taxa specialised on millipedes), saprophagy and mycophagy, with many species being rather generalist (Davis et al., 2008).

**Remarks**. Onthophagini and Oniticellini have long been considered separate tribes. However, there is now a large amount of evidence, both morphological (Tarasov and Solodovnikov, 2011; Bai et al., 2011) and molecular (present analyses; Ayivi et al., 2021; Breeschoten et al., 2016; Cheng et al., 2021; Guo et al., 2022; Lopes et al., 2024; Villalba et al., 2002; Stone et al., 2026), showing indisputably that Oniticellini are deeply nested within Onthophagini (Fig. 2).

The two characters traditionally used to separate the two tribes are the the presence, in Oniticellini, of a mesothoracic scutellum (absent in Onthophagini) and of 8-segmented antennae (9-segmented in Onthophagini). However, both these characters are also present in some Onthophagini, with a few species showing a mesoscutellum (Ochi and Kon, 2017) and many genera showing 8 antennomeres (Cambefort, 1979, 1981, 1986). On the other hand, the two groups share two important synapomorphies: 1) the presence of a wide sclerite posteriorly to RP1 vein (also present in the unrelated Sisyphini and Epirinini); and 2) the presence of cavities in the first and second club antennomeres (also present in other groups, see (Philips, 2016)). In addition to these characters, Onthophagini and Oniticellini have the same configuration of elytral striae, genital features (especially the widespread presence of a highly species-specific lamella copulatrix and the SRP not ring-shaped), as well as a very similar general appearance.

For these reasons, we down-rank Oniticellini to a subtribe of the Onthophagini, so that it becomes Oniticellina *stat. nov*.

Several subtribes were described within Oniticellini: Attavicinina Philips, 2016 (originally incorrectly spelled Attavicina), Drepanocerina Lansberge, 1875, Helictopleurina Janssens, 1946, Liatongina Philips, 2016, and Oniticellina Kolbe, 1905 (Philips, 2016). Helictopleurina was found to be sister to all remaining oniticellines by both morphology (Philips, 2016) and UCEs (present analyses; Lopes et al., 2024), although one mitogenomic study recovered it within Oniticellina (Breeschoten et al., 2016). Helictopleurina comprises a single genus *Helictopleurus* d’Orbigny, 1915, which is endemic to Madagascar and has a somewhat peculiar morphology compared to other Oniticellini, specifically the body less elongated on average and a characteristically wide elytral interstria 8. The group appears to be distinct phylogenetically, biogeographically and morphologically from the other oniticellines, so we keep it as a separate subtribe, Onthophagini Helictopleurina *stat. nov*.

As to the other subtribes, molecular (present analyses; Breeschoten et al., 2016; Stone et al., 2026) and morphological (Philips, 2016) phylogenies consistently showed that members of Drepanocerina and Liatongina are nested within a paraphyletic Oniticellina. Maintaining them as valid subtribes would require further splitting Oniticellina into several, morphologically poorly defined and mono- or oligogeneric taxa. We argue that this solution would only complicate classification without adding much biological meaning. Therefore, we propose the following two new synonymies: Oniticellina Kolbe, 1905 = Drepanocerina Lansberge, 1875 *syn. nov.* = Liatongina Philips, 2016 *syn. nov*. As to Attavicinina, its members (*Attavicinus* Philips & Bell, 2008 and *Paroniticellus* Balthasar, 1963) were never included in any molecular phylogeny, while morphological analyses found them to form a clade sister to Oniticellina (Philips, 2016). However, we believe that subtribal names should be maintained only when they are supported by a substantial amount of morphological, ecological and biogeographical evidence in addition to phylogenetic relationships. It is our opinion that these conditions are not met for Attavicinina, which we therefore synonymize with Oniticellina = Attavicinina *syn. nov*.

A much more complex situation involves the subtribe Onthophagina. With the subtribal classification proposed above, Onthophagina clearly emerges as a polyphyletic group composed of Oniticellina, Helictopleurina and several other lineages (Fig. 2). The only other named subtribe of Onthophagini is Alloscelina Janssens, 1949, originally described within Scarabaeini to accommodate myrmecophilous or termitophilous taxa sharing the presence of only 8 antennomeres and other highly homoplastic traits in leg and body shape: *Haroldius* (Haroldiini) and *Parachorius* Harold, 1873 (Parachoriini) and their synoynms, as well as the true Onthophagini *Alloscelus* Boucomont, 1923 and *Megaponerophilus* Ferreira, 1972 (now *Caccobiomorphus* Balthasar, 1964) (Janssens, 1949; Branco, 2011). The polyphyly of Alloscelina is now widely acknowledged, and the taxon—currently considered a subtribe of Onthophagini—is *de facto* unused (Krell, 2010; Philips, 2016). There is good morphological evidence that *Alloscelus* and *Caccobiomorphus* are not sister genera and are both deeply nested within the “true” Onthophagina lineage, with at least the former being probably allied to other myrmecophilous or termitophilous genera of the *Stiptopodius*-group, such as *Eusaproecius* Branco, 1989 (see Fig. 2) (Philips, 2016; Branco, 1996). We therefore formally propose the following synonymy: Onthophagina Streubel, 1846 = Alloscelina Janssens, 1949 *syn. nov*. The three valid onthophagine subtribes can be distinguished using the identification key provided below.

The taxa allied to the genus *Phalops* Erichson, 1848 (*Cacconemus* Jekel, 1872, *Digitonthophagus* Balthasar, 1959, *Hamonthophagus* Roggero, Dierkens, Barbero & Palestrini, 2017, *Kurtops* Roggero, Barbero & Palestrini, 2017 and several species currently placed in *Onthophagus*) form a clade sister to the remaining Onthophagini (present analyses; Ayivi et al., 2021; Breeschoten et al., 2016; Génier and Moretto, 2017; Stone et al., 2026). The group has highly modified accessory endophallites, with FLP, SA, A and some additional sclerites more or less extensively fused, and lacking a lamella copulatrix (Génier and Moretto, 2017; Tarasov and Solodovnikov, 2011). These features seem to represent a derived condition, as all the other members of the Onthophagomorpha clade have distinct accessory endophallites. On the other hand, the *Phalops*-group shows some putative onthophagomorphan plesiomorphic characters such as an often less developed, thinner posterior sclerite of the wing RP1 vein (*e.g.*, in *Digitonthophagus*, *Hamonthophagus*), the constant presence of a male protibial uncus and the presence, in most *Phalops* species, of a reduced, additional elytral stria between striae 7 and 8. Notwithstanding this combination of derived and plesiomorphic traits, the general morphology and ecology of these taxa is so similar to that of the remaining Onthophagini that we fully support the retention of these genera within the tribe.

Besides these considerations, resolving Onthophagini subtribal classification is a difficult endeavour due to the large number of genera involved, so we postpone it to a dedicated future paper. For now, all remaining Onthophagini shall be placed in Onthophagina *sensu lato*.

### Panelini Arrow, 1931 *stat. rev.* and *sensu novo*

(Fig. 21)

**Figure 21.**
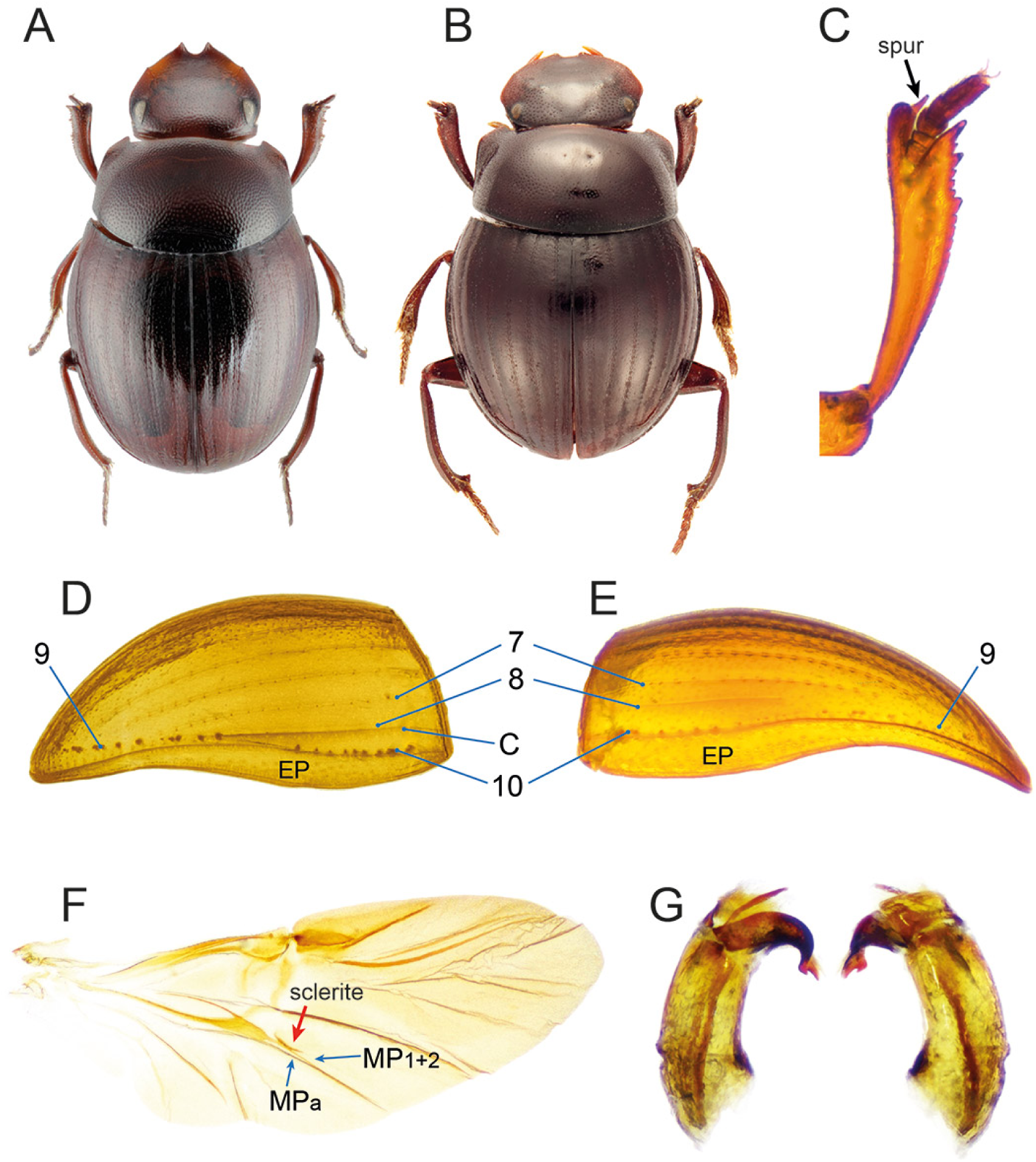
Diagnostic morphological characters of Panelini *sensu novo*. (A) *Panelus* (*Panelus*) sp., habitus; (B) *Panelus* (*Macropanelus*) *sumatrensis*, habitus; (C) *Panelus* sp., protibia with arrow showing spur inserted on digitiform process of protibial apex; (D) *Panelus* sp. and (E) *Panelus rhodesiensis*, elytra showing lateral striae (7–10) and carinae (C; epipleural carina not marked); (F) *Panelus* sp., metathoracic wing showing anterior sclerite of MP1+2 vein; (G) *P. rhodesiensis*, strongly asymmetric aedeagus in left and right lateral views.

**Type genus**: *Panelus* Lewis, 1895.

**Genus included:**

*Panelus* Lewis, 1895 (38 valid species; Afrotropical, Indomalayan, Palaearctic).

**Diagnosis**. The tribe is defined by the following unique combination of characters: 1) hind wing with anterior sclerite of MP1+2 vein curved posteriorly, its extremities not fused with the vein (Fig. 21F); 2) 8th or 9th elytral stria reduced to a series of points on the posterior part of elytron, either indistinct and restricted to the distalmost part of elytron (Fig. 21E), or aligned with pseudoepipleural carina (Fig. 21D); 3) protibial spur inserted into digitiform apical process of protibia (Fig. 21C); 4) parameres asymmetrical (Fig. 21G).

**Description.**

Very small to small sized beetles (ca. 1.5–5.5 mm); body oval, convex.

*Head*. Anterior margin of clypeus with 2 teeth; frontoclypeal sulcus absent; head surface without protrusions. Antennae with 9 antennomeres.

*Pronotum*. Subrectangular, convex, surface evenly punctate; lateral pronotal carina entire.

*Elytra and wings*. Elytron usually with 10 striae, stria 8 sometimes indistinct and/or subcarinate; pseudoepipleural carina immediately lateral to stria 8 present or not; second-to-last lateral stria (8 or 9) restricted to a row of points on the posterior half of elytron, usually aligned with the elytral carina or tightly adjacent to the lateralmost stria; epipleuron relatively wide. Mesoscutellum concealed. Metathoracic wings present; anterior sclerite of MP1+2 vein curved posteriorly, its extremities not fused with the vein. *Legs*. Protibia with 3 teeth; protibial spur inserted into digitiform apical process of protibia; protibial distal margin obliquely truncated; protarsus present; uncus absent. Meso- and metatibiae feebly expanded proximo-distally, more or less sinuous. Tarsal claws not toothed.

*Ventral body surface*. Anterior region of hypomeron depressed; anterior hypomeral carina stretching towards lateral margin of hypomeron; posterior longitudinal hypomeral carina absent.

*Tergite VIII*. Wider than long, evenly convex.

*Genitalia*. Parameres asymmetrical. Endophallites: A, SA, FLP and SRP present; LC absent.

**Distribution and biology**. The genus *Panelus* is widely distributed in the Indomalayan realm, with a few species reaching the Palaearctic and only one taxon, *P. rhodesiensis* Paulian, 1975, being endemic to the Afrotropical realm. It comprises two subgenera, *Panelus* and *Macropanelus* Ochi, Kon & Araya, 1998, the latter restricted to Indonesia (Sumatra) and with currently uncertain validity (Tarasov, 2017). *Panelus* species are small forest-dwellers; some species were found in dung baited-traps, but very little is known about other life history traits (Davis et al., 2008; Masumoto et al., 2011).

**Remarks**. Arrow (1931) described the tribe Panelini for a group of genera characterised by a “very compact form, with the middle coxae far apart and and nearly parallel, and with quite simple tarsi, of which the first four joints are nearly equal in size” (p. 404): *Delopleurus* (now in Coptorhinini), *Haroldius* (now in Haroldiini), *Panelus*, *Paraphytus* (now in Paraphytini), *Ponerotrogus* (currently synonym of *Haroldius*), and *Pycnopanelus* (now in Pycnopanelini). The tribe was soon considered a synonym of Canthonini, in which *Panelus* was included (Balthasar, 1963; Matthews, 1974; Scholtz and Howden, 1987; Davis et al., 2008) until Tarasov and Dimitrov (2016) classified it as *incertae sedis*.

Morphologically, the general habitus of *Panelus* resembles that of *Bohepilissus* (Bohepilissini), *Lepanus* (Mentophilini) and *Tanzanolus* (Tanzanolini), so that species of the latter two genera were initially placed in *Panelus* (Scholtz and Howden, 1987; Matthews, 1974). However, significant differences in elytral striation, wing venation and genitalia show that similarity is probably due to convergence. This is confirmed by our molecular reconstruction as well as that by Gunter et al. (2026), in which these genera are very distantly related, except for *Bohepilissus* which was often found close or sister to *Panelus*. Overall, both morphological and molecular evidence converge in identifying *Panelus* as a highly supported, rather isolated monogeneric tribe, Panelini *stat. rev.* and *sensu novo*.

Apparently, paneline morphology has remained largely similar during evolution. This is exemplified by the close similarity between the only known African species, *P. rhodesiensis*, and Asian species, which, in spite of a very deep divergence (Fig. 1), show substantial morphological homogeneity. Meaningful variation occurs mostly in elytral striation. In most of the Asian species examined here, elytra have a carina between striae 8 and 9, and stria 9 is composed by a series of points aligned with the posterior end of stria 8, so that it looks a continuation of that stria (Fig. 21D). However, a small shift between the position of the two striae suggests that they are in fact distinct. In *P. rhodesiensis*, the carina is absent (Fig. 21E), and in at least one unidentified species from Borneo even stria 8 is virtually undistinguishable, so that the total number of striae appears to be eight. In the latter two cases, striae 8 and 9 are not aligned and the latter is narrowly adjacent to the posterior part of stria 10.

### Parachoriini Tarasov, 2017

**Type genus**: *Parachorius* Harold, 1873.

**Genus included:**

*Parachorius* Harold, 1873 (20 valid species; Indomalayan).

**Remarks**. The tribe was described by Tarasov (2017) based on solid molecular and morphological evidence, so we avoid discussing it again here. We were not able to include *Parachorius* in our reconstruction, but previous works recovered a close relationship with Epactoidini, Epirinini and Sisyphini (Fig. 1; see Discussion).

### Paraphytini Scholtz, Davis & De Klerk, 2025

**Type genus**: *Paraphytus* Harold, 1877.

**Genus included:**

*Paraphytus* Harold, 1877 (14 valid species; Afrotropical, Indomalayan).

**Remarks**. The tribe was recently described by Scholtz et al. (2025) based on solid molecular and morphological evidence, so we avoid discussing it again here. The group falls within the Coptorhinomorpha clade, where it is most often found to be sister to Coptorhinini (Fig. 1; see Discussion).

### Pycnopanelini *trib. nov*

(Figs. 22, 3D)

**Figure 22.**
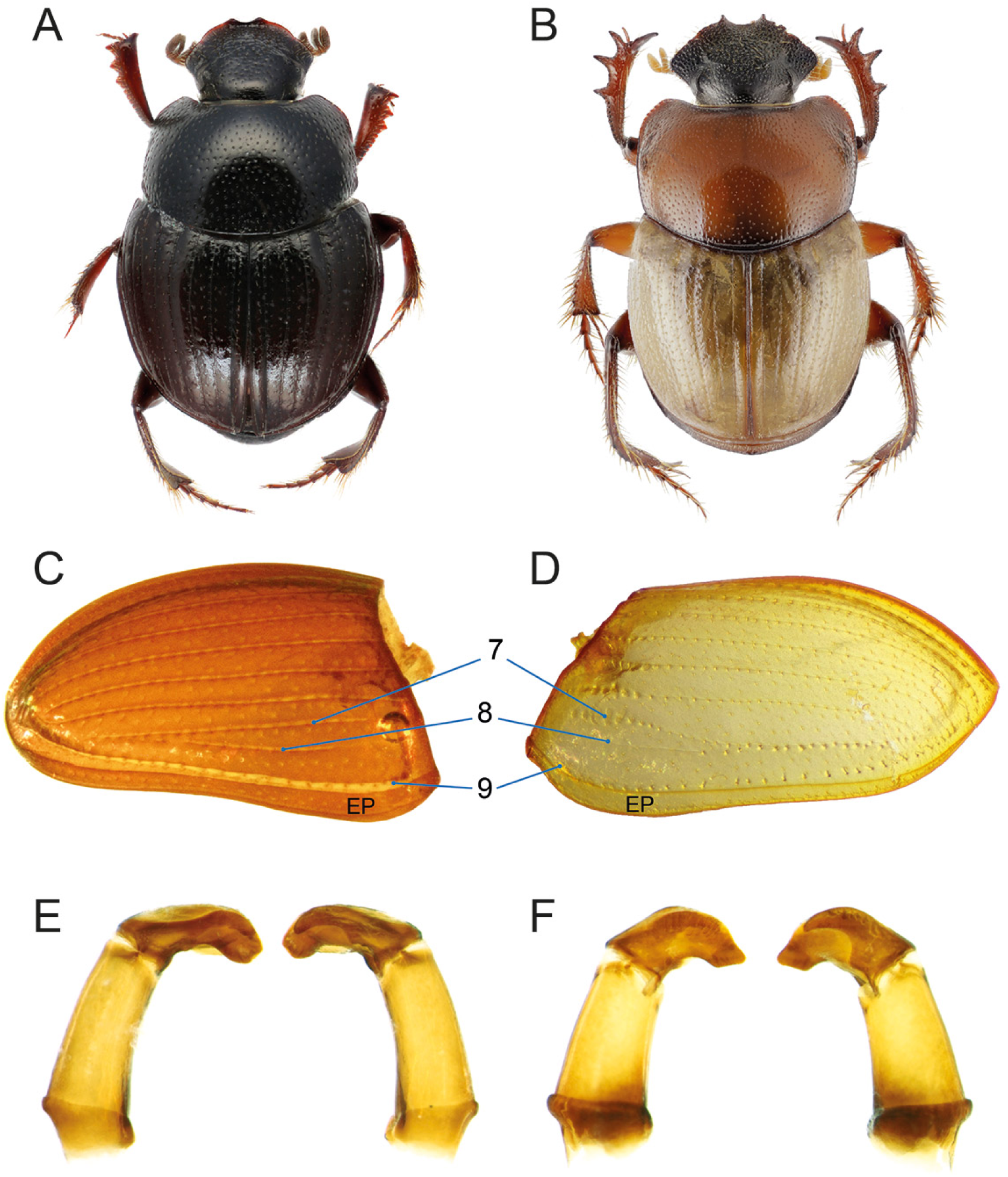
Diagnostic morphological characters of Pycnopanelini *trib. nov.* (A) *Pycnopanelus krikkeni*, habitus; (B) *Hammondantus psammophilus*, habitus; (C) *P. krikkeni* and (D) *H. psammophilus*, elytra showing lateral striae (7–9); (E) *P. krikkeni* and (F) *H. psammophilus*, aedeagi in left and right lateral views.

**Type genus**: *Pycnopanelus* Arrow, 1931.

**Genera included:**

*Pycnopanelus* Arrow, 1931 (3 valid species; Afrotropical, Indomalayan).

*Hammondantus* Cambefort, 1978 (1 valid species; Afrotropical).

**Diagnosis**. The tribe is defined by the following unique combination of characters: 1) second lateral stria restricted to medial portion of elytron, either effaced anteriorly and posteriorly (*Pycnopanelus*, Fig. 22C) or effaced anteriorly and fusing with stria 7 posteriorly (*Hammondantus*, Fig. 22D); 2) protibial uncus present in males; 3) parameres asymmetrical, left paramere widely and obtusely rounded, right paramere widely notched ventrally (Figs. 22E–F).

**Description.**

Small sized beetles (3.0–6.0 mm); moderately convex.

*Head*. Clypeus with two teeth; head surface without carinae or protrusions (except for *Pycnopanelus parvicollis*); frontoclypeal sulcus absent. Antennae with 9 antennomeres.

*Pronotum*. Finely punctured, without protrusions; lateral pronotal carina entire.

*Elytra and wings*. Elytron with 8 or 9 visible striae; second lateralmost stria restricted to medial portion of elytron, either effaced anteriorly and posteriorly (*Pycnopanelus*) or effaced anteriorly and fusing with stria 7 posteriorly (*Hammondantus*). Absence of elytral carinae except for the than the epipleural one. Mesoscutellum concealed. Metathoracic wings present.

*Legs*. Protibia with 3 teeth; uncus present in males. Mesotibiae and female metatibiae evenly expanded proximo-distally; male metatibiae curved at half length, strongly expanded distally. Tarsal claws present and not toothed in *Pycnopanelus*, absent in *Hammondantus*.

*Ventral body surface*. Anterior region of hypomeron depressed (*Pycnopanelus*) or not (*Hammondantus*); anterior hypomeral carina present (*Pycnopanelus*) or absent (*Hammondantus*); posterior longitudinal hypomeral carina absent.

*Tergite VIII*. More elongated in males than in females.

*Genitalia*. Parameres asymmetrical, left paramere widely and obtusely rounded, right paramere widely notched ventrally. Endophallites: LC absent; A, SA, FLP endophallites present; SRP with ring-shaped part.

**Distribution and biology**. The tribe presents a fragmented distribution. *Hammondantus* comprises only *H. psammophilus* Cambefort, 1978, a psammophilous species restricted to the dune fields of southern Namibia (Davis et al., 2008, 2020). *Pycnopanelus* comprises three species with disjunct Indoafrican distribution, being found in India (*P. rotundus* Arrow, 1931, the type species), and in northeastern (*P. parvicollis* (d’Orbigny, 1913)) and southwestern (*P. krikkeni* Cambefort, 1978) Africa. While *P. krikkeni* is relatively common, the other two species are known only from 2 specimens each (Cambefort, 1978).

**Remarks**. Arrow (1931) described the genus *Pycnopanelus* within his tribe Panelini together with genera currently assigned to other tribes (see Panelini section). Panelini soon became a synoynm of Canthonini, where *Pycnopanelus* was accommodated (Janssens, 1949; Scholtz and Howden, 1987). *Hammondantus* was also placed in Canthonini, and its affinities with *Pycnopanelus* were highlighted since its description (Cambefort, 1978; Scholtz and Howden, 1987). Both genera were officially excluded from Canthonini by Tarasov and Génier (2015) and treated as *incertae sedis*.

Our analyses fully support a sister relationship between the two genera and their placement within the Onthophagomorpha clade (Fig. 2; see Discussion), in agreement with previous molecular and morphological works (Tarasov and Génier, 2015; Sole and Scholtz, 2010; Tarasov and Dimitrov, 2016; Mlambo et al., 2015). Our tree 70p-1 found full support for a sister relationship between Pycnopanelini and Onthophagini, which was not recovered in the other 70p trees nor by previous works. However, such relationship is further supported by the observation that, unlike other onthophagomorphans, these two tribes lack elytral carinae except for the epipleural one, and have a reduced number of elytral striae (8 or 9 in Pycnopanelini; 8 or very rarely 9 in Onthophagini; usually 10 in the other Onthophagomorpha). Additionally, their general habitus is somehow similar, so that in the past *Pycnopanelus parvicollis* d’Orbigny, 1913 was described in the onthophagine genus *Caccobius* Thomson, 1859 (d’Orbigny, 1913). Nevertheless, the two lineages differ in many significant characters including wing venation, the degree of development of the penultimate lateral elytral stria, and features of male genitalia and legs (especially the metatibiae strongly curved and thickened in Pycnopanelini). Overall, the establishment of Pycnopanelini *trib. nov.* is fully justified both phylogenetically and morphologically.

Curiously, *Pycnopanelus* species show different degrees of development of elytral striae 7 and 8, as already observed by Cambefort (1978). *P. krikkeni* has a clearly visible stria 8, although limited to the middle of the elytron, while in *P. rotundus* the stria is reduced to a very short line. In *P. parvicollis*, stria 8 is completely missing and stria 7 is restricted to the middle of the elytron.

### Scarabaeini Latreille, 1802

(Fig. 23, 3I)

**Figure 23.**
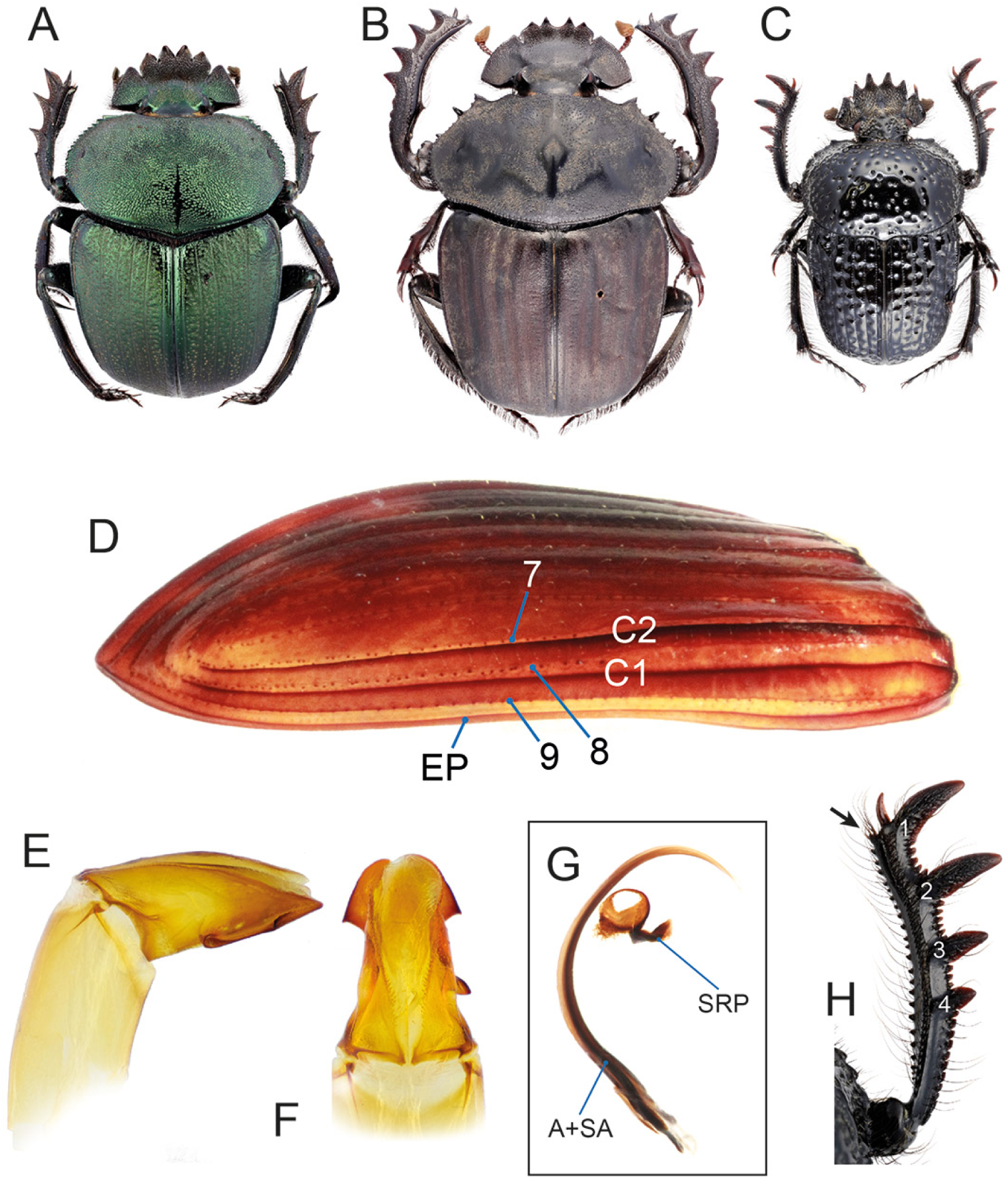
Diagnostic morphological characters of Scarabaeini. (A) *Kheper* sp., habitus; (B) *Pachylomera femoralis*, habitus; (C) *Scarabaeolus* sp., habitus; (D) *Scarabaeus* sp., elytron showing lateral striae (7–9) and carinae (C1, C2); *Kheper* sp., parameres in left lateral (E) and dorsal (F) views; (G) *Kheper aegyptiorum*, endophallites; (H) *Scarabaeolus* sp., protibia showing number of lateral teeth and absence of tarsus (black arrow).

**Type genus**: *Scarabaeus* Linnaeus, 1758.

**Genera included:**

*Ateuchetus* Bedel, 1892 (10 valid species; Afrotropical, Palaearctic);

*Escarabaeus* Zídek & Pokorný, 2011 (8 valid species; Afrotropical, Indomalayan, Palaearctic);

*Kheper* Janssens, 1940 (27 valid species; Afrotropical, Indomalayan);

*Mnematidium* Ritsema, 1888 (1 valid species; Palaearctic);

*Mnematium* MacLeay, 1821 (2 valid species; Palaearctic);

*Pachylomera* Griffith & Pidgeon, 1832 (2 valid species; Afrotropical);

*Pachysoma* MacLeay, 1821 (13 valid species; Afrotropical);

*Parateuchus* Shipp, 1895 (1 valid species; Afrotropical);

*Scarabaeolus* Balthasar, 1965 (45 valid species; Afrotropical, Palaearctic);

*Scarabaeus* Linnaeus, 1758 (66 valid species; Afrotropical, Indomalayan, Palaearctic);

*Sceliages* Westwood, 1837 (7 valid species; Afrotropical).

**Diagnosis**. The tribe is defined by the following unique combination of characters: 1) elytron with 9 striae, sometimes striae 8 and 9 indistinct (Fig. 23D); 2) elytron usually carinated between striae 7–8 and 8–9, carinae sometimes narrowly adjacent to fused (Fig. 23D); 3) protarsus absent in both sexes (Fig. 23H); 4) protibia with 4 teeth (Fig. 23H); 5) subaxial and axial endophallites long, occupying at least half of endophallus (Fig. 23G).

**Description.**

Moderately small to very large sized beetles (7.3–48.5 mm); body flattened to convex.

*Head*. Clypeal margin with 4 teeth, lateralmost ones sometimes reduced; genal margin more or less strongly toothed anteriorly; anterior head margin usually notched in correspondence with genoclypeal sulcus; head surface with or without protrusions; frontoclypeal sulcus either invisible or carinate or tuberculate. Antennae with 9 antennomeres.

*Pronotum*. Flattened to convex, without protrusions; variously punctate or granulate; lateral pronotal carina entire.

*Elytra and wings*. Elytron with 9 visible striae, sometimes 8th and/or 9th striae indistinct; carinated between striae 7–8 (except at least in *Ateuchetus catenatus* (Gerstaecker, 1871)) and 8–9, the two carinae sometimes narrowly adjacent to almost fused, with stria 8 hardly visible between them. Mesoscutellum concealed or not. Metathoracic wings present or absent.

*Legs*. Protibia with 4 teeth; uncus present or absent; protarsi absent in both sexes. Meso- and metatibiae feebly expanded proximo-distally. Tarsal claws not toothed, sometimes absent.

*Ventral body surface*. Anterior hypomeral carina absent; posterior longitudinal hypomeral carina absent. Metaventrite usually bulging anteriorly; flat to slightly concave in at least some *Pachysoma*, where it is strongly constricted between mesocoxae.

*Tergite VIII*. Evenly convex, subtriangular.

*Genitalia*. Parameres asymmetrical (Figs. 23E–F). Endophallites: LC absent; A and SA endophallites present, at least as long as half of endophallus; FLP present, reduced or absent; SRP ring-shaped; raspulae usually present.

**Distribution and biology**. Scarabaeini are widely distributed in the Afrotropical realm, where their diversity peaks, as well as in the Indomalayan and Palaearctic realms. They are rollers feeding mostly on various types of dung, although some species are attracted to carrion and the genus *Sceliages* is specialised in exploiting millipedes (Davis et al., 2008). The greatest majority of Scarabaeina occurs in open habitats, from savanna to desert, with a few exceptions inhabiting dry forests in Madagascar (Montanaro and Tarasov, 2024) or coastal dune forests in KwaZulu-Natal (South Africa) and southeastern Mozambique (*Scarabaeus bornemisszai* zur Strassen, 1980; Davis et al., 2020).

**Remarks**. The composition of Scarabaeini has been little questioned historically due to its morphological homogeneity. Its monophyly is largely confirmed on a molecular (present analyses; Tarasov and Dimitrov, 2016; Mlambo et al., 2015; Gunter et al., 2026; Scholtz et al., 2011; Monaghan et al., 2007) and morphological basis (Tarasov and Génier, 2015).

However, the current generic classification of the tribe is rather problematic, being largely affected by extensive paraphyly of *Scarabaeus*, whose species are widely interspersed among other genera (present analyses; Harrison, 2003; Forgie et al., 2005, 2006; Scholtz et al., 2011). This is reflected in the fact that most genus-group names have oscillated between subgeneric and generic rank (Forgie et al., 2005). All those taxa are now considered genera, with the exception of *Scarabaeus* (*Pachylosoma*) Zídek & Pokorný, 2008 and *Sceliages*, formally considered a subgenus of *Scarabaeus* but regarded as a full genus by some authors (here, we adopt the latter interpretation) (Davis et al., 2020; Deschodt et al., 2015). Overall, a dedicated, fine scale investigation of the relationships between the genera is urgently needed to fix the classification of Scarabaeini (Montanaro and Tarasov, 2024).

Our analyses found a fully supported sister relationship with the monotypic genus *Circellium* (Circelliini *trib. nov.*), in agreement with some previous studies (Tarasov and Génier, 2015; Tarasov and Dimitrov, 2016). However, the deep divergence between the two lineages, as well as significant morphological and biogeographical differences between them, justify their accommodation into separate tribes (see Circelliini section).

### Sisyphini Mulsant, 1842

**Type genus**: *Sisyphus* Latreille, 1807.

**Genera included:**

*Indosisyphus* Barbero, Palestrini & Zunino, 1991 (1 species; Indomalayan);

*Nesosisyphus* Vinson, 1946 (5 valid species; Madagascan (Mauritius));

*Sisyphus* Latreille, 1807 (105 valid species; Afrotropical, Neotropical, Indomalayan, Palaearctic).

**Remarks**. This morphologically homogeneous tribe has been quite stable historically. More recently, Tarasov and Dimitrov (2016) proposed to merge it with the sister tribe Epirinini based on close morphological and molecular results, but Daniel et al. (2020b) separated them again, which is the currently accepted solution (see also Epirinini section). A sound definition of Sisyphini was already given by those authors, and further consolidated by Daniel (2019), Daniel et al. (2020a) and Losacco et al. (ress), so we avoid discussing it here. However, the generic delimitation of the tribe is still in progress (Daniel, in preparation). Our reconstructions fully confirm the sister relationship with Epirinini (Fig. 1; see Discussion).

### Tanzanolini *trib. nov*

(Fig. 24)

**Figure 24.**
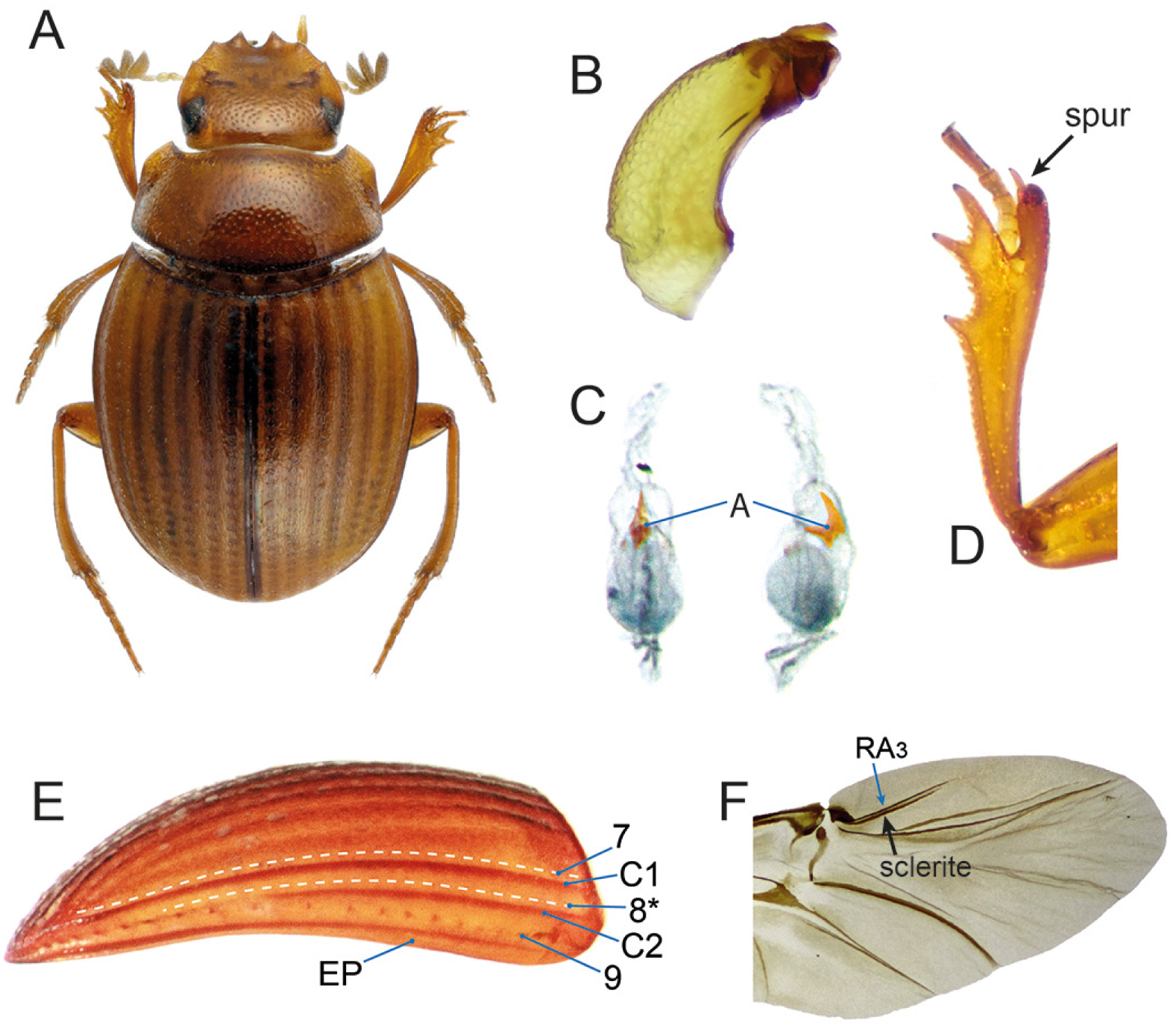
Diagnostic morphological characters of Tanzanolini *trib. nov.* *Tanzanolus leleupi*: (A) habitus; (B) aedeagus in lateral view; (C) endophallus; (D) protibia showing spur inserted on digitiform process of tibial apex; (E) elytron showing striae (7–9) and carinae (C1, C2; epipleural carina not marked); (F) metathoracic wing showing elongated and thin posterior sclerite of RA3 vein.

**Type genus**: *Tanzanolus* Scholtz & Howden, 1987.

**Genus included:**

*Tanzanolus* Scholtz & Howden, 1987 (1 valid species; Afrotropical).

**Diagnosis**. The tribe is defined by the following unique combination of characters: 1) posterior sclerite of RA3 thin and elongated, extending from base of RA3 to its median part (Fig. 24F); 2) SA, FLP, SRP endophallites and LC absent (Fig. 24C); 3) elytron with 9 striae (Fig. 24E); 4) elytron carinated between striae 7–8 and 8–9 (Fig. 24E).

**Description.**

Very small sized beetles (2.0–3.0 mm); body oval, elongated, moderately convex.

*Head*. Anterior margin of clypeus with 4 teeth, genae toothed next to genoclypeal sulcus; frontoclypeal sulcus absent. Antennae with 9 antennomeres.

*Pronotum*. Moderately convex, densely punctured; lateral pronotal carina entire.

*Elytra and wings*. Elytra with 9 visible striae, stria 8 reduced to a rather indistinct row of punctures; elytron carinated between striae 7–8 and 8–9, lateralmost carina becoming effaced on posterior half. Mesoscutellum concealed. Metathoracic wings present; posterior sclerite of RA3 vein thin and elongated, extending from base of RA3 to its median part.

*Legs*. Protibia with 3 teeth; protibial distal end obliquely truncated, distoventral margin strongly produced into a digitiform process bearing protibial spur; protarsus present. Meso- and metatibiae evenly and feebly expanded proximo-distally. Tarsal claws not toothed.

*Ventral body surface*. Anterior region of hypomeron depressed; anterior hypomeral carina stretching towards lateral margin of hypomeron; posterior longitudinal hypomeral carina present; in males, 5th visible abdominal sternite extremely short medially, constricted by long 6th sternite.

*Tergite VIII*. Wider than long, convex, longitudinally subcarinated medially.

*Genitalia*. Parameres symmetrical. Endophallites: A endophallite present; SA, FLP, SRP and LC absent.

**Distribution and biology**. The only described species, *Tanzanolus leleupi* (Balthasar, 1960), is known from the rainforest of the Uluguru Mountains in Tanzania (Scholtz and Howden, 1987). It has been collected in forest litter and, in one occasion, attracted to a trap baited with carrion and millipedes (GM, unpublished); nothing else is known about its life history. Additional, undescribed populations are known to occur elsewhere in the Tanzanian Eastern Arc Mountains (Montanaro *et al*., unpublished).

**Remarks**. *Tanzanolus* was originally placed in the Canthonini (Scholtz and Howden, 1987) and later moved to *incertae sedis* (Tarasov and Dimitrov, 2016). Previous molecular works found it as sister to the genus *Janssensantus* (Janssensantini) (Mlambo et al., 2014) or even nested within it, possibly due to contamination (Tarasov and Dimitrov, 2016). Our reconstructions place it in a very isolated position together with *Nesovinsonia* (Nesovinsoniini). Both genera are relics occurring in isolated rainforests: the former in the Tanzanian Eastern Arc Mountains, the latter on Mauritius in the middle of the Indian Ocean. *Tanzanolus* can be readily distinguished from all other Scarabaeinae by its unique wing configuration, particularly the shape of the posterior sclerite of RA3. Morphological features of elytral carination, endophallites, legs and general habitus resemble those of *Bohepilissus* (tribe Bohepilissini *trib. nov.*) and, to a lesser extent, *Panelus* (tribe Panelini). However, the three genera seem completely unrelated based on all available studies (see remarks on Bohepilissini for further details). No meaningful synapomorphies could be found between *Tanzanolus* and its sister, *Nesovinsonia*. Thus, phylogenetic and morphological evidence strongly supports the establishment of Tanzanolini *trib. nov*.

## IDENTIFICATION KEY TO AFRO-EURASIAN TRIBES AND SUBTRIBES

**1.** Elytron with 10 clear striae; striae 8–10 placed laterally to medialmost elytral carina (*i.e.*, placed on pseudoepipleuron) (Figs. 15D–E) ……….. **Heliocoprini**

**–.** Elytron with 10 striae or less; lateralmost 3 striae never all placed laterally to medialmost elytral carina (*i.e.*, never all placed on pseudoepipleuron). Instead, either pseudoepipleural carina absent, or only 1 or 2 striae placed on pseudoepipleuron ……….. **2**

**2.** Epipleural carina effaced on posterior half (Fig. 16D). Posterior longitudinal hypomeral carina present (Fig. 16E). Tarsal claws toothed (Fig. 16B) ……….. **Janssensantini**

**–.** Epipleural carina not effaced on posterior half. Posterior longitudinal hypomeral carina present or absent; if present, tarsal claws not toothed ……….. **3**

**3.** Anterior clypeal margin with 6 teeth (Fig. 4A). Medialmost elytral carina (pseudoepipleural carina) adjoining epipleural carina at half elytral length (Fig. 16D). Lateral elytral margin not emarginated on anterior half (Fig. 16D) ……….. **Aphengoecini**

**–.** Anterior clypeal margin usually with less than 6 teeth; if with 6 teeth, then pseudoepipleural carina absent and lateral elytral margin emarginated on anterior half ……….. **4**

**4.** Lateral elytral margin clearly emarginate on anterior half ……….. **5**

**–.** Lateral elytral margin not emarginate on anterior half, at most slightly concave ……….. **6**

**5.** Lateral surface of sternites 2–4 curved dorsally, visible from above in the lateral elytral emargination (Fig. 13D). Metanepisternum subrectangular, more or less straight dorsally (Fig. 13G). Elytron with 9 striae; hind wings normally sclerotised, not darkened ……….. **Gymnopleurini**

**–.** Lateral surface of sternites 2–4 not curved dorsally, invisible from above. Metanepisternum expanded dorsally into the elytral emargination. Elytron with 10 striae; hind wings strongly sclerotised, darkened **Coptorhinini**

**6.** Epipleuron very wide, at least as wide as the two lateralmost interstriae taken together (*e.g.*, Fig. 14C). Fore leg without trochanterofemoral pit. Endophallites comprising only BSc and A sclerite with elongated spur-shaped medial lobe (TS) (*e.g.*, Fig. 14F) ……….. **7**

**–.** Epipleuron usually not very wide, usually less wide than the two lateralmost interstriae taken together; if epipleuron very wide, fore leg with trochanterofemoral pit (Fig. 12F). Endophallites variously shaped**8**

**7.** Antennae with 8 segments (Fig. 14D). Small-sized (ca. 1.5–3.3 mm), lenticular beetles.. **Haroldiini**

**–.** Antennae with 9 segments. Medium-sized (4.9–10.8 mm), convex but never lenticular beetles **Elassocanthonini**

**8.** Fore leg with trochanterofemoral pit (Figs. 10F, 12F, 12L) ……….. **9**

**–.** Fore leg without trochanterofemoral pit ……….. **11**

**9.** Elytron with 10 striae, without carinae except for the epipleural one (Fig. 9B). Posterior region of hypomeron depressed or grooved (Fig. 10E). Aedeagus strongly asymmetric (Fig. 10G). Body elongated, heavily punctate (Fig. 10I). Body length: 5.0–15.0 mm ……….. **Coprini Pedariina**

**–.** Elytron with 9 striae, sometimes striae 8 and 9 poorly visible; elytron carinated between striae 7 and 8 (Figs. 12A, 12G). Posterior region of hypomeron not depressed nor grooved. Aedeagus symmetric. Body rounded, variously punctate. Body length: 1.5–4.8 mm ……….. **10 (Endroedyolini)**

**10.** Body very convex, covered with long, conspicuous setae (Fig. 12C). Clypeus almost always without small medial tooth (Fig. 12B). Hind wings absent. Endophallites comprising only BSc and A sclerite with elongated spur-shaped lobe (TS) (Fig. 12E) ……….. **Endroedyolina**

**–.** Body less convex, covered with short setae (Fig. 12I). Clypeus always with small medial tooth (Fig. 12H). Hind wings usually present. Endophallites comprising BSc and two additional sclerites (Fig. 12K) **Odontolomina**

**11.** Distal margin of protibia with groove accommodating protarsus (*e.g.*, Fig. 10A) ……….. **12**

**–.** Distal margin of protibia without groove accommodating protarsus ……….. **13**

**12.** Last tarsomeres expanded and concealing tarsal claws (Fig. 10B). Elytron with carina lateral to stria 8 (Fig. 10C) ……….. **Coprini Onychothecina**

**–.** Last tarsomeres not expanded and not concealing tarsal claws. Elytron without carina lateral to stria 8 **Paraphytini**

**13.** Posterior region of pronotum with two symmetrical depressions (Fig. 19E). Metathoracic scutellum visible between elytra (Fig. 19E). Male protibia without spur and protarsus, with an uncus (Fig. 19F) **Onitini**

**–.** Posterior region of pronotum without two symmetrical depressions. Metathoracic scutellum visible or not between elytra. Male protibia either with or without spur, protarsus and uncus ……….. **14**

**14.** Mesanepisternum carinated dorsoventrally (Fig. 6C). Elytron with medialmost elytral carina lateral to stria 8 (Figs. 6D–E). Hind wing with J vein well developed and AP3+4 reduced (Fig. 6F).. **Catharsiini**

**–.** Mesanepisternum not carinated dorsoventrally. Elytron with or without medialmost elytral carina lateral to stria 8. Hind wing venation different from above ……….. **15**

**15.** Elytron with 10 clear striae; striae 9–10 placed laterally to medialmost elytral carina (pseudoepipleural carina) (*e.g.*, Figs. 11C, 11D, 18B). Protibial spur not placed on digitiform distal process of protibia. **16**

**–.** Elytron either without pseudoepipleural carina, or with less than 10 striae. Protibial spur placed on digitiform distal process of protibia (*e.g.*, Fig. 24D) or not ……….. **19**

**16.** Elytron carinated posteriorly between striae 7 and 8. Anterior margin of clypeus with 4 teeth. Putatively extinct species from La Réunion (Mascarene archipelago)***Epactoides giganteus* (Epactoidini)**

**–.** Elytron not carinated posteriorly between striae 7 and 8. Anterior margin of clypeus with 2 teeth or less, occasionally with a feeble bulging laterally to the main 2 teeth. Distribution different ……….. **17**

**17.** Protibia with 4 teeth (*e.g.*, Fig. 17C) ……….. ***Macroderes* (Macroderini)**

**–.** Protibia with 3 teeth (*e.g.*, Fig. 11E) ……….. **18**

**18.** Elytron clearly setose (Figs. 11A–B). In males, protibia with uncus, without spur (Figs. 11E). In females, anterior margin of clypeus between clypeal teeth sharp; lateral margin of genae pointed (Fig. 11B). Endemic to Dwesa Forest (South Africa) ……….. **Dwesasilvasedini**

**–.** Elytron not setose (Fig. 18A). In males, protibia without uncus, with spur. In females, anterior margin of clypeus between clypeal teeth rounded; lateral margin of genae rounded. Endemic to Mauritius (Mascarene archipelago) ……….. **Nesovinsoniini**

**19.** Elytron with 8 visible striae, with two carinae converging posteriorly between striae 7 and 8 (Fig. 5D). Tarsal claws toothed proximally; ventro-distal margin of male protibia with scale-like setae (Fig. 5E). Endophallus comprising only A, SRP and LC endophallites (Fig. 5B). Small beetles (1.7–3.1 mm) **Bohepilissini**

**–.** Elytron with 7 or more visible striae, with at most one carina between striae 7 and 8 (*e.g.*, Fig. 7D); if two narrowly adjacent carinae present, then both striae 8 and 9 placed on the pseudoepipleuron. Tarsal claws not toothed proximally; ventro-distal margin of male protibia with or without scale-like setae. Composition of endophallites different. Body size variable ……….. **20**

**20.** Protibial spur placed on digitiform apical process of protibia (Figs. 21C, 24D) ……….. **21**

**–.** Protibial spur not placed on digitiform apical process of protibia ……….. **22**

**21.** Elytron with 9 striae (stria 8 sometimes indistinct); carinated betwen striae 7–8 and 8–9 (Fig. 24E). Fore wing with RA3 vein having a thin, elongated sclerite extending from base of vein to its median part (Fig. 24F). Endophallites comprising only A sclerite (Fig. 24C) ……….. **Tanzanolini**

**–.** Elytron with 10 striae (stria 8 sometimes indistinct); stria 9 visible only posteriorly, where it seems a continuation of stria 10; elytron with at most one carina between striae 8 and 10 (Figs. 21D–E). Fore wing without thin, elongated sclerite of RA3 vein (Fig. 21F). Endophallites comprising A, SA, FLP and SRP sclerites ……….. **Panelini**

**22.** Absence of elytral carinae other than the epipleural one (*e.g.*, Figs. 9A, 22C–D) ……….. **23**

**–.** Elytron with one or two carinae in addition to the epipleural one (*e.g.*, Figs. 7D, 23D) ……….. **30**

**23.** Elytron with 10 clearly visible striae, striae 9–10 sometimes closely adjacent but distinct at least posteriorly (Figs. 9A, 17E). Protibia with 4 teeth ……….. **24**

**–.** Elytron with 8 or 9 striae, penultimate stria sometimes restricted to the middle of elytral length (*e.g.*, Figs. 22C–D). Protibia with 4 teeth or less ……….. **25**

**24.** Dorsal side of metatibia with short transverse carina. Male protibia without uncus (Fig. 9D). Posterior longitudinal hypomeral carina usually present (Fig. 9G). Hind wing with posterior sclerite of RP1 vein present (Fig. 9F) ……….. **Coprini Coprina**

**–.** Dorsal side of metatibia without short transverse carina. Male protibia with uncus (Fig. 17C). Posterior longitudinal hypomeral carina absent. Hind wing with posterior sclerite of RP1 vein absent.. ***Xinidium* (Macroderini)**

**25.** Middle and hind legs very elongated, length of metatibia+metatarsus longer than elytron+pronotum. Surface of posterolateral region of hypomeron flat, approximately coplanar, perpendicular to frontal body plane ……….. **Sisyphini**

**–.** Middle and hind legs relatively short, length of metatibia+metatarsus shorter than elytron+pronotum. Surface of posterolateral region of hypomeron sinuous, not coplanar, not perpendicular to frontal body plane ……….. **26**

**26.** Metaventrite with anterior longitudinal carina connected to mesometaventral suture. Phallobase with a pair of tubercles anterodorsally. Protibia with 3 teeth ……….. **Parachoriini**

**–.** Metaventrite without anterior longitudinal carina connected to mesometaventral suture. Phallobase without pair of tubercles anterodorsally (*e.g.*, Fig. 9E). Protibia with 4 teeth or less ……….. **27**

**27.** Elytron with 9 or, in one case, 8 striae; penultimate stria shortened, not reaching either the anterior or posterior elytral margin (Figs. 22C–D). Hind wing without large posterior sclerite of RP1 vein. Male protibia with uncus; male metatibia curved inwards distally. Parameres asymmetrical (Figs. 22E–F) **Pycnopanelini**

**–.** Elytron with 8 or, very rarely, 9 striae (Fig. 20F); penultimate stria not shortened, except the 8th one in some species of *Phalops*. Hind wing usually with large posterior sclerite of RP1 vein (Fig. 20E). Male protibia with or without uncus; male metatibia not curved inwards distally. Parameres most often symmetrical ……….. **28 (Onthophagini)**

**28.** Antennae with 8 antennomeres. Mesothoracic scutellum usually present, if absent then body elongated, flattened, dorsal integument strongly sculptured and setose, and basolateral plate of parameres present**29**

**–.** Antennae with 8 or 9 antennomeres. If mesothoracic scutellum present, antennae with 9 antennomeres. If antennae with 8 antennomeres and scutellum absent, then body shape, sculpture and parameres different from above ……….. **Onthophagina (*sensu lato*)**

**29.** Elytral interstria 8 at least twice as wide as interstria 7. Mesothoracic scutellum always present. Dorsal margin of tergite VIII not carinated. Endemic to Madagascar ……….. **Helictopleurina**

**–.** Elytral interstria 8 not or slightly wider than interstria 7. Mesothoracic scutellum present or, rarely, absent. Dorsal margin of tergite VIII carinated or not. Absent from Madagascar ……….. **Oniticellina**

**30.** Elytron with only one stria laterally to pseudoepipleural carina, stria short and sometimes indistinct. Anterior clypeal margin with only 2 sharp teeth. Metaventrite between mesocoxae normally developed, not narrowly constricted. Hind wing, when present, with large posterior sclerite of RP1 vein (similar to Fig. 20E) ……….. **Epirinini**

**–.** Elytron with two striae laterally to pseudoepipleural carina (*e.g.*, Fig. 8D), and/or anterior clypeal margin with 4 sharp teeth (*e.g.*, Figs. 23A–C), and/or mesocoxae narrowly adjacent and metaventrite strongly constricted between them. Hind wing, when present, without large posterior sclerite of RP1 vein ……….. **31**

**31.** Protibia with 4 teeth or more (Fig. 23H) ……….. **Scarabaeini**

**–.** Protibia with 3 teeth (*e.g.*, Figs. 7E, 8E) ……….. **32**

**32.** Body size smaller (2–9 mm). Protarsi present ……….. **Epactoidini**

**–.** Body size bigger (11–50 mm). Protarsi absent ……….. **33**

**33.** Elytron relatively short, approximately as long as 1.3 times length of pronotum. Protibia without uncus or uncus-like projections (Fig. 8A). Endophallites comprising very long, separable A and SA sclerites, putative FLP reduced to an inconspicuous small sclerotisation (Fig. 8C). Body strongly convex, almost hemispherical (Fig. 8A) ……….. **Circelliini**

**–.** Elytron relatively long, at least 1.7 times length of pronotum, except in *Canthodimorpha* where it is 0.9–1.5 times pronotal length. Protibia with uncus or uncus-like projections both in males and females (Fig. 7E). Endophallites comprising tightly wrapped, almost inseparable A and SA sclerites and a well developed FLP (Figs. 7F, 7H). Body less strongly convex, not hemispherical (Figs. 7A–C)**Chalconotini**

## DISCUSSION

Our UCE-based phylogenetic reconstructions recovered 27 fully supported clades that we interpret as tribe-rank taxa, encompassing the vast majority of Afro-Eurasian dung beetle lineages. Parachoriini and Aphengoecini were not sampled in our dataset, but independent molecular evidence supports their recognition as separate tribes (Tarasov, 2017; Tarasov and Dimitrov, 2016). By integrating molecular results with morphology and previous phylogenetic studies, we delimit 13 newly described tribes and re-evaluate several others. This increases the number of valid scarabaeine tribes from 24 to 36 and provides a classification for virtually all Afro-Eurasian genera previously designated as *incertae sedis*, thereby resolving long-standing uncertainties in dung beetle classification. The only genus still unplaced is the fossil *Lobateuchus*, whose phylogenetic placement is unclear.

Our dataset did not include most of the American and Australasian taxa. However, available molecular studies indicate that those genera are unlikely to be closely related to the Afro-Eurasian lineages treated herein (Gunter et al., 2026; Lopes et al., 2024; Tarasov and Dimitrov, 2016). Several American *incertae sedis* genera appear to occupy rather isolated positions (*e.g.*, *Bdelyrus*, *Canthonella*+*Canthochilum*, *Tesserodoniella*, *Uroxys*), whereas others (*e.g.*, *Ontherus*, *Canthidium*) belong to an endemic American clade comprising Dichotomiini, Eucraniini, Homocoprini, Phanaeini, and part of Ateuchini (Gunter et al., 2026; Tarasov and Dimitrov, 2016). These results suggest limited impact of unsampled non-Afro-Eurasian taxa on the tribal definitions established here. We therefore expect the classification proposed in this study to remain largely stable when a comprehensive global phylogeny of Scarabaeinae (in prep.) including American endemic genera becomes available. Even if some Neotropical genera are unexpectedly recovered within Afro-Eurasian tribes, they can be readily accommodated within the classification proposed here.

### Major Afro-Eurasian dung beetle clades

Our multiple analytical approaches consistently recovered the same tribal-level clades, whereas relationships among tribes remained less well resolved. Several tribes shifted positions across alternative topologies, precluding robust hypotheses of their higher-level relationships. Nevertheless, several deeper clades were recovered with strong support.

One of them is Coptorhinomorpha (Fig. 1) (70p-1: UFBoot=99.8, SH-aLRT=99). This group is the most basally derived in the whole subfamily and is united by a unique configuration of endophallic sclerites, comprising only a BSc and an axial sclerite usually bearing a thin medial lobe. The only exception is the genus *Odontoloma*, which seems to have a somewhat more derived endophallite configuration comprising sclerites with unclear homology. Coptorhinomorpha is further structured into Paraphytini+Coptorhinini (70p-1: UFBoot=100, SH-aLRT=100), sharing a more or less notched lateral elytral margin and a relatively narrow epipleuron; and Haroldiini+(Elassocanthonini+Endroedyolini) (70p-1: UFBoot=100, SH-aLRT=100), sharing a very wide epipleuron. Support for the monophyly and position of this clade is confirmed by morphology (Tarasov and Génier, 2015), although earlier molecular works recovered Coptorhinini+Paraphytini and Haroldiini+Elassocanthonini+Endroedyolini as sequentially diverging from the root of the tree (Gunter et al., 2026; Mlambo et al., 2015; Monaghan et al., 2007; Sole and Scholtz, 2010; Tarasov and Dimitrov, 2016; Tarasov, 2017).

A second well-supported clade (70p-1: UFBoot=100, SH-aLRT=99) is Onthophagomorpha (Fig. 2), which was previously identified by other studies (Gunter et al., 2026; Mlambo et al., 2015; Tarasov and Génier, 2015; Tarasov and Dimitrov, 2016). Its members share the presence of unci, a clear plesiomorphic character retained throughout the group and lost only within the extensive radiation of Onthophagini *sensu novo* and occasionally in Catharsiini. Onthophagomorpha most likely has an Afrotropical origin, as all included genera are either endemic to or centred in Africa.

A clade comprising Sisyphini, Epactoidini and Epirinini is also fully supported (70p-1: UFBoot=100, SH-aLRT=100) by our analyses and previous phylogenies (Gunter et al., 2026; Tarasov and Dimitrov, 2016). Four out of 10 of our UCE trees recovered *Bohepilissus* as either nested within or sister to the clade, with *Panelus* being often sister to the whole cluster. Tarasov and Dimitrov (2016) and Tarasov (2017) found that the lineage also contains the Parachoriini and Eurysternini, although Gunter et al. (2026) did not support the inclusion of the latter tribe and Rossini et al. (2022) found Onthophagini nested within it. Unfortunately, morphology does not provide further evidence. Given its incompletely resolved composition, the clade will require further consolidation in broader future phylogenetic analyses.

Chalconotini and Circelliini+Scarabaeini form a clade in three of our trees (50p-5, 70p-4, 70p-5), consistently with the results of Gunter et al. (2026), Lopes et al. (2024), and Tarasov and Génier (2015). This relationship is also supported by morphology: all tribes lack protarsi in both sexes, share a similar pattern of elytral striation (identical in Chalconotini and Circelliini), and possess asymmetrical parameres. In addition, they comprise medium- to large-sized roller species with predominantly African distributions (entirely so in Circelliini and Chalconotini). Intriguingly, Tarasov and Génier (2015) recovered the Neotropical genus *Streblopus* within this clade, as sister to Circelliini+Scarabaeini. This placement is morphologically plausible: *Streblopus* shares the same elytral configuration as Chalconotini and Circelliini, has asymmetrical parameres and the endophallites comprising only SRP and elongated A and SA, similarly to Scarabaeini and Circelliini. Its general habitus is also notably similar to that of Chalconotini. If confirmed, this relationship would lend support to the biogeographic hypothesis of Cupello et al. (2020), which proposed a transoceanic dispersal of *Streblopus* from Africa. However, the UCE phylogeny of Gunter et al. (2026) did not recover the same relationships, so that a more comprehensive analysis will be needed to test them.

### Relict Afrotropical taxa

Seven tribes are here newly established or revalidated for former Canthonini lineages. Most are restricted to the Afrotropics, where they occur either in (rain)forest (Bohepilissini, Dwesasilvasedini, Janssensantini, Tanzanolini) or in more open habitats (Aphengoecini), while others are endemic to the Mascarene archipelago (Nesovinsoniini) or shared between the Afrotropical and Indomalayan realms (Panelini).

Among these, Dwesasilvasedini and Aphengoecini are best interpreted as relict onthophagomorphan lineages, nowadays restricted to very small areas in South Africa (Davis et al., 2020). The remaining tribes occupy isolated phylogenetic positions and likely represent the last remnants of once more widespread forest-specialised clades. Several of them—particularly Tanzanolini, Panelini, and Bohepilissini—share a superficially similar morphology, being small-bodied forest dwellers with comparable body and leg shape. This resemblance raises the question of whether it reflects convergence driven by similar ecological constraints or the retention of an ancestral forest-dwelling body plan that became extensively modified in other dung beetle lineages which adapted to different habitats. Although the latter possibility is appealing, the occurrence of a comparable ground plan in the Australasian *Lepanus* (Menthophilini), where it is likely derived (Gunter and Weir, 2019; Gunter et al., 2026), points instead toward convergence as the most parsimonious explanation.

Additionally, these forest taxa exhibit compelling biogeographic patterns. Panelini, which are diverse and widespread in the Indomalayan realm, are represented in Africa by a single described species, *P. rhodesiensis* Paulian, 1975. Despite their conserved morphology, the African and Asian lineages are deeply divergent molecularly, suggesting that *P. rhodesiensis* is a relict of a formerly more widespread Afrotropical paneline clade. A similar scenario emerges from the relationship between Tanzanolini and Nesovinsoniini, which are restricted to the Eastern Arc Mountains and the Mascarene islands, respectively. These groups may represent remnants of an ancestral African lineage that later dispersed eastward into the Mascarenes. Both these hypotheses are consistent with an African origin of the clades, followed by dispersal into other regions and then extensive extinction across their original home ranges. This pattern would perfectly coincide with the history inferred for Epactoidini, which, from East Africa, colonised Asia, Madagascar and subsequently the Mascarene islands, with African forms now surviving only in rainforest patches in the Eastern Arc Mountains, exactly like Tanzanolini (Montanaro et al., 2024b; Rossini et al., 2022).

## CONCLUSIONS

This study substantially revises the tribal classification of Scarabaeinae dung beetles by integrating UCE-based phylogenomics with morphology. We redefine previous tribal concepts, describe 13 new tribes, and provide the first nearly complete tribal classification of Afro-Eurasian dung beetles, with the exception of a few *incertae sedis* genera from Madagascar. This work increases the number of recognised tribes from 24 to 36 and reduces the number of *incertae sedis* genera from 45 to 22, leaving only six Madagascan genera, 15 American genera, and the Eocene fossil genus *Lobateuchus* unplaced. We also recovered two well-supported higher-level clades—Coptorhinomorpha and Onthophagomorpha—that clarify deeper relationships within the subfamily and provide a phylogenetic basis for future studies of scarabaeine evolution. Finally, we showed that many former Afro-Eurasian Canthonini represent phylogenetically isolated, relictual lineages, likely of Afrotropical origin, highlighting their complex evolutionary histories. Our results provide a foundation for a comprehensive global phylogeny and stable tribal classification of Scarabaeinae, facilitating future revisions of the remaining unplaced taxa and enabling investigations of dung beetle evolution across time and space.

## ACKNOWLEDGMENTS

We are sincerely thankful to many colleagues: Giulio Cuccodoro (Muséum d’histoire naturelle, Geneva, Switzerland), Jiří Hájek (National Museum, Prague, Czech Republic), Keita Matsumoto and Max Barclay (Natural History Museum, London, UK), Jaakko Mattila (Finnish Museum of Natural History, Helsinki, Finland), Olivier Montreuil (Muséum National d’Histoire naturelle, Paris, France) and Philippe Moretto (Toulon, France) for facilitating access to specimens under their care; Vasily Grebennikov (Canadian Food Inspection Agency, Ottawa, Canada) and Werner P. Strümpher (Ditsong National Museum of Natural History, Pretoria, South Africa) for providing valuable specimens of rare canthonines; Elina Laiho (Finnish Museum of Natural History) for help with DNA extractions; Hennie De Klerk (University of Pretoria, Pretoria, South Africa) for the wonderful pictures of live beetles, and Christian Deschodt (University of Pretoria, Pretoria, South Africa), Giulio Cuccodoro and François Génier (Canadian Museum of Nature, Ottawa, Canada) for those of preserved specimens (respective credits are in figure captions); Fernando Vaz-de-Mello (Universidade Federal de Mato Grosso, Cuiabá, Brazil) for sharing his extensive scarab bibliography; Alberto Ballerio (Brescia, Italy) for valuable nomenclatural advice. We thank the Malagasy Institut pour la Conservation des Écosystèmes Tropicaux (MICET) and the Mauritian National Parks and Conservation Service (NPCS) for providing permits to collect samples for DNA analyses in Madagascar and Mauritius. Gimo M. Daniel is supported by the Instituto Nacional de Coleoptera (INCol) and the CNPq National Institute of Science and Technology (INCT) (grant no. 408430/2024-9).

## SUPPLEMENTARY MATERIAL

All supplementary files are accessible at https://osf.io/f4kxu.

## ADDITIONAL INFORMATION AND DECLARATIONS

## Competing Interests

The authors declare that they have no competing interests.

## Author Approvals

All authors have seen and approved the manuscript, which has not been accepted or published elsewhere.

## Funding

This work was supported by the Research Council of Finland (369299 and 331631), and the National Science Foundation (NSF 1942193).

## Author Contributions

ST, GM, FLop and NG conceived the work. All authors contributed to data acquisition. FLop performed molecular analyses. GM developed tribal diagnoses. GM and FLos acquired figures. ST and NG provided the project resources, supervision and administration. GM wrote the initial draft with support from FLop and ST. All authors validated the work and contributed to reviewing and editing the final draft.

## Notes

https://osf.io/f4kxu

### Competing Interest Statement

The authors have declared no competing interest.

